# CSsingle: A Unified Tool for Robust Decomposition of Bulk and Spatial Transcriptomic Data Across Diverse Single-Cell References

**DOI:** 10.1101/2024.04.07.588458

**Authors:** Wenjun Shen, Cheng Liu, Yunfei Hu, Yuanfan Lei, Hau-San Wong, Si Wu, Xin Maizie Zhou

## Abstract

We introduce CSsingle, a novel method that enhances the decomposition of bulk and spatial transcriptomic (ST) data by addressing key challenges in cellular heterogeneity. CSsingle applies cell size correction using ERCC spike-in controls, enabling it to account for variations in RNA content between cell types and achieve accurate bulk data deconvolution. In addition, it enables fine-scale analysis for ST data, advancing our understanding of tissue architecture and cellular interactions, particularly in complex microenvironments. We provide a unified tool for integrating bulk and ST with scRNA-seq data, advancing the study of complex biological systems and disease processes. The benchmark results demonstrate that CSsingle outperforms existing methods in accuracy and robustness. Validation using more than 700 normal and diseased samples from gastroesophageal tissue reveals the predominant presence of mosaic columnar cells (MCCs), which exhibit a gastric and intestinal mosaic phenotype in Barrett’s esophagus and esophageal adenocarcinoma (EAC), in contrast to their very low detectable levels in esophageal squamous cell carcinoma and normal gastroesophageal tissue. We revealed a dynamic relationship between MCCs and squamous cells during immune checkpoint inhibitors (ICI)-based treatment in EAC patients, suggesting MCC expression signatures as predictive and prognostic markers of immunochemotherapy outcomes. Our findings reveal the critical role of MCC in the treatment of EAC and its potential as a biomarker to predict outcomes of immunochemotherapy, providing insight into tumor epithelial plasticity to guide personalized immunotherapeutic strategies.

## Introduction

Deciphering cellular heterogeneity is crucial when investigating the microenvironment of a tissue relevant to disease, as it plays a crucial role in identifying specific cell populations that are potential therapeutic targets.^1, 2^ High-throughput sequencing technologies have revolutionized cellular research and offer unparalleled insight into the complexity and dynamics of living systems. Conventional gene expression profiling technologies, such as microarrays or bulk RNA-sequencing (RNA-seq) have successfully measured an enormous number of bulk samples due to technical simplicity and low cost. However, bulk RNA-seq technologies, which measure averaged expression across cell populations, hinder the elucidation of cellular signatures driving tumor initiation and progression. The advent of single-cell RNA sequencing (scRNA-seq) technologies has revolutionized the study of cellular heterogeneity by offering unprecedented resolution and genome-wide range.^3–5^ However, the high cost and the requirement for high quality tissues impede the use of this approach in clinical investigations that typically involve a considerable cohort of participants.^6, 7^ Spatial transcriptomics (ST), such as 10x Genomics Visium^8^ and Slide-Seq,^9^ captures the entire transcriptome while preserving spatial context but lacks singlecell resolution.^10^ Over the last two decades, numerous computational methods have been developed to infer cell-type composition from cell-type mixtures, commonly known as cell-type deconvolution. Integrating scRNA-seq with bulk RNA-seq and ST data provides a more holistic view of cellular diversity and tissue organization. This synergy enhances the study of complex biological systems and disease processes, advancing both basic and translational research.^11–13^ Accurate decomposition of cell type mixtures relies on four key components (i) a specialized signature matrix of celltype-specific gene expression profiles (GEPs), (ii) the ability to account for both technical (e.g., discrepancies between sequencing methods) and biological (e.g., disease-related variations) differences between the mixture and reference data, (iii) the incorporation of cell size factors when mixing cell types with significantly different RNA contents, and (iv) the ability to filter noise from unrelated cell types in the reference data, particularly for deconvolving ST data.

To build a signature matrix, many existing deconvolution methods^14–18^ utilized raw gene counts, which can be read counts or UMI (Unique Molecular Identifier) counts generated by scRNA-seq.^19^ This kind of method does not adequately address the significant impact of sequencing depth (total number of reads or UMI molecules per cell) on gene expression measurement. When the sequencing depth varies significantly between cells in the single-cell reference data, the resulting estimates of cell type proportions may not be accurate. On the other hand, many deconvolution methods have used single-cell GEPs measured by transcripts per million (TPM) or counts per million (CPM)-normalized values to account for variable sequencing depth.^20–24^ However, both TPM and CPM-based methods were designed to address library size heterogeneity through count depth scaling, assuming all cells initially contain equal RNA contents.^25^ Bulk transcriptomic technologies measure the average gene expression across a population of cells, which inherently includes contributions from cells of varying sizes. Larger cells contribute more RNA to the bulk sample. Hence, using such a signature matrix to deconvolve bulk mixtures of cell types with significantly different sizes would yield inaccurate cell-type proportion estimates without cell-size correction. To our knowledge, the impact of cell size differences on deconvolution accuracy has not been systematically evaluated.

Existing bulk deconvolution methods focus primarily on addressing the influence of gene feature selection in reference profiles.^14, 15, 18, 20^ Empirical evidence indicates that gene feature selection or gene weighting strategies result in a higher deconvolution accuracy. However, only a few studies have focused on addressing the technical and biological disparities between bulk and reference data. Bisque^24^ learns gene-specific transformations of bulk data to correct for technical biases between the bulk and reference data. CIBER-SORTx,^23^ an extension of CIBERSORT,^20^ employs a batch correction method to reduce cross-platform variation between bulk mixtures and the signature matrix. In particular, CIBERSORTx requires at least three bulk mixtures, with a recommendation of ten, to effectively perform the batch correction procedure. Resolving technical and biological variations between a single bulk mixture and the signature matrix remains a critical challenge.

Existing spatial deconvolution methods^26–32^ have made significant progress in inferring the distribution of cells within tissue samples and enabling spatial mapping of cellular heterogeneity. Despite these achievements, they face notable limitations. These methods often fail to fully account for technical and biological discrepancies between spatial and reference data, significantly affecting the accuracy of estimates.^13^ Furthermore, effective filtering of noise from unrelated cell types, particularly at the level of individual spatial spots, remains a significant challenge. These limitations highlight the need for more accurate, robust, and high-resolution deconvolution techniques in spatial transcriptomics.

To address these prevailing challenges, we introduce CSsingle (Cross-Source SINGLE cell decomposition), an innovative deconvolution method for accurate, robust and efficient estimation of cell type composition from bulk and ST data across diverse single-cell references. By explicitly addressing both biological and technical variations inherent in individual cell-type mixtures and the signature matrix, CSsingle significantly outperforms current state-of-the-art methods for deconvolution tasks involving cross-source heterogeneity. Moreover, CSsingle is the first tool to apply cell size correction using ERCC spike-in controls, enabling it to account for variations in RNA content between cell types and achieve accurate bulk data deconvolution. In this study, we systematically investigate the role of cell size correction in enhancing deconvolution accuracy. Additionally, by precisely identifying enriched cell types at single-spot resolution, CSsingle facilitates high-resolution analytical capabilities for spatial transcriptomics data. We demonstrate the effectiveness of CSsingle through comprehensive analyzes of multiple single-cell, bulk and ST data sets, including GEPs from the mouse brain cortex, human pancreas, peripheral blood mononuclear cells, whole blood, normal squamous-columnar junction, Barrett’s esophagus, and esophageal carcinoma. The results underscore the versatility and robustness of CSsingle, positioning it as a powerful tool for integrative analyses of single-cell, bulk and ST data across diverse biological and clinical contexts.

## Results

### Existing deconvolution methods fail to consider differences in cell size

The deconvolution problem is typically cast in the form of a linear equation system: **y** = **S** × **X**, where **X** denotes the cell type composition of the cell type mixture **y**, and **S** denotes a signature matrix. Computational methods aim to deconvolve cell type proportion **X** by using the signature matrix **S**. To build a signature matrix, existing deconvolution methods normally use single-cell GEPs measured by read / UMI counts or TPM / CPM normalized values.^20–23^ We observed that these deconvolution methods, which formulate **y** = **S** × **X** to estimate cellular fractions for the bulk sample **y**, often introduce a systematic bias in estimation when the bulk sample consists of cell types with significantly different cell sizes, commonly referred to as the absolute cellular RNA contents. To illustrate this, we considered a real data set containing External RNA Controls Consortium (ERCC) spike-ins^33, 34^ introduced by Konstantin et al.,^35^ which was a compilation of 14 mixtures from two cell types (HEK and Jurkat cells) of markedly different cell sizes. ERCC spike-in controls were added to each mixture to eliminate systematic technical differences in expression between the mixtures, e.g. systematic variation introduced during the library preparations or sequencing of the samples. Each mixture in this data set, consisting of 10^6^ cells, is commonly assumed to contain equivalent quantity of mRNA molecules by existing deconvolution methods employing TPM- or CPM-normalized values. However, we observed that the absolute RNA contents of the mixtures after spike-in normalization were significantly correlated with the proportions of HEK cells (Fig.1a), suggesting distinct cell sizes between HEK and Jurkat cells. In this data set, Samples 1, 2 and Samples 13, 14 are mixtures of pure HEK and Jurkat cells, respectively. Fig.1b indicated that HEK cells had markedly larger cell sizes than Jurkat cells after spike-in normalization. Without spike-in normalization, the estimated cell size difference is less pronounced between HEK and Jurkat cells than with spike-in normalization (Fig.1c versus 1b), which is due to technical effects introduced during RNA extraction, amplification or sequencing.

**Figure 1.**
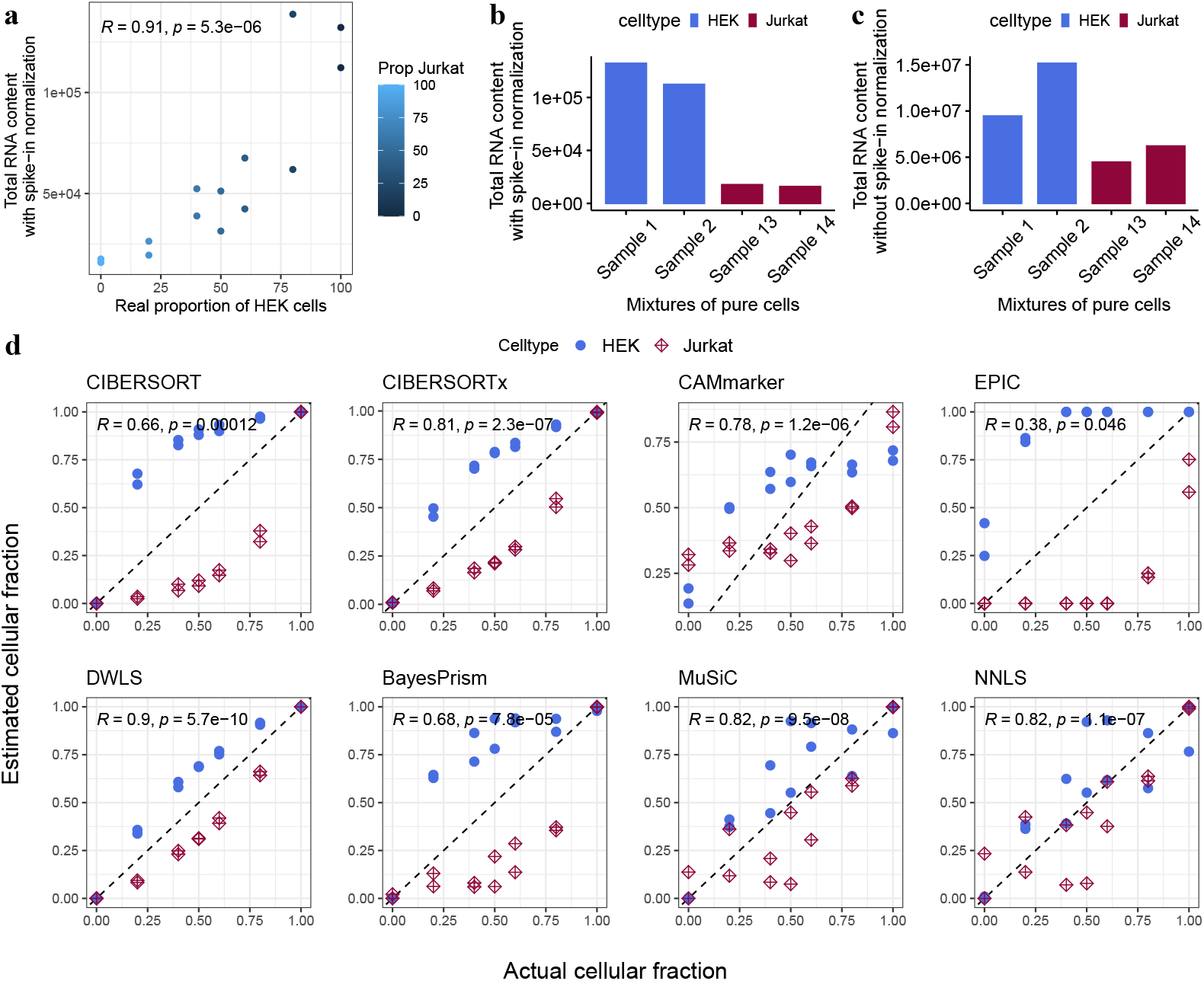
Cell type composition in mixtures of two cell types of markedly different total RNA contents: HEK and Jurkat cells. **a** Scatter plot showing Pearson correlation between absolute RNA contents and real proportions of HEK cells for bulk mixtures. **b** Absolute RNA contents estimated using ERCC spike-in controls for each cell type. **c** RNA contents estimated by summing gene counts for each cell without ERCC spike-in normalization. **d** Performance of existing bulk deconvolution methods on the mixtures of HEK (blue circles) and Jurkat (red squares) cells. The estimated cell type proportions are plotted against the actual cell type proportions. Reported ‘R’ corresponds to Pearson’s correlation coefficient and *p*-values indicate the significance of these correlations.

For this real data set, we then used mixtures of pure HEK and pure Jurkat cells to generate cell-type-specific GEPs to construct the signature matrix. One mixture of pure HEK cells, i.e. Sample 1 was excluded from the signature matrix construction due to technical bias (Fig.1b). All 14 bulk mixtures were employed for benchmarking existing deconvolution methods against the ground truth. CIBERSORT, CIBER-SORTx, CAMmarker,^22^ and EPIC^21^ constructed signature matrices by normalizing the gene counts to CPM or TPM, while DWLS, BayesPrism,^17^ MuSiC and NNLS^14^ used raw gene counts. As expected, all methods yielded estimated cellular fractions that systematically deviated from the actual cellular fractions (Fig.1d). Specifically, for the deconvolution methods using a CPM or TPM-normalized signature matrix, cellular fractions of the HEK cell type, characterized by larger cell size, were consistently overestimated. Conversely, cellular fractions for the Jurkat cell type, with smaller cell size, were consistently underestimated (Fig.1d, top panels). The potential source of this bias in cellular fraction estimations is often overlooked because of the commonly held, though rarely explicitly stated, assumption in deconvolution that the absolute amount of total mRNA is equivalent across different cell types. Both CPM and TPM-based methods intend to correct for library size using count depth scaling and assume that all cells are initially characterized by an equivalent quantity of mRNA molecules.^25^ As a result, they produce an erroneous estimation of cellular fractions due to the marked difference in cell sizes between the two cell types. Alternatively, we also tested deconvolution methods using signature matrices of raw read counts. Since these methods do not correct for bias arising from “technical” library size differences, cell sizes cannot be accurately estimated. As a result, the estimated cellular fractions did not align with the actual cellular fractions (Fig.1d, bottom panels). These results highlighted the importance of incorporating cell size factors in computational deconvolution.

### An iteratively reweighted least-squares approach to robustify deconvolution estimates

CSsingle tackles cellular heterogeneity challenges by (i) integrating cell size coefficients into deconvolution to correct cell size bias, and (ii) managing technical and biological variations between individual mixtures and the signature matrix. An overview of CSsingle is shown in Fig.2a. In the context of spatial transcriptomics, CSsingle enhances the high-resolution analytical potential by accurately identifying enriched cell types at single-spot resolution (Fig.2b).

**Figure 2.**
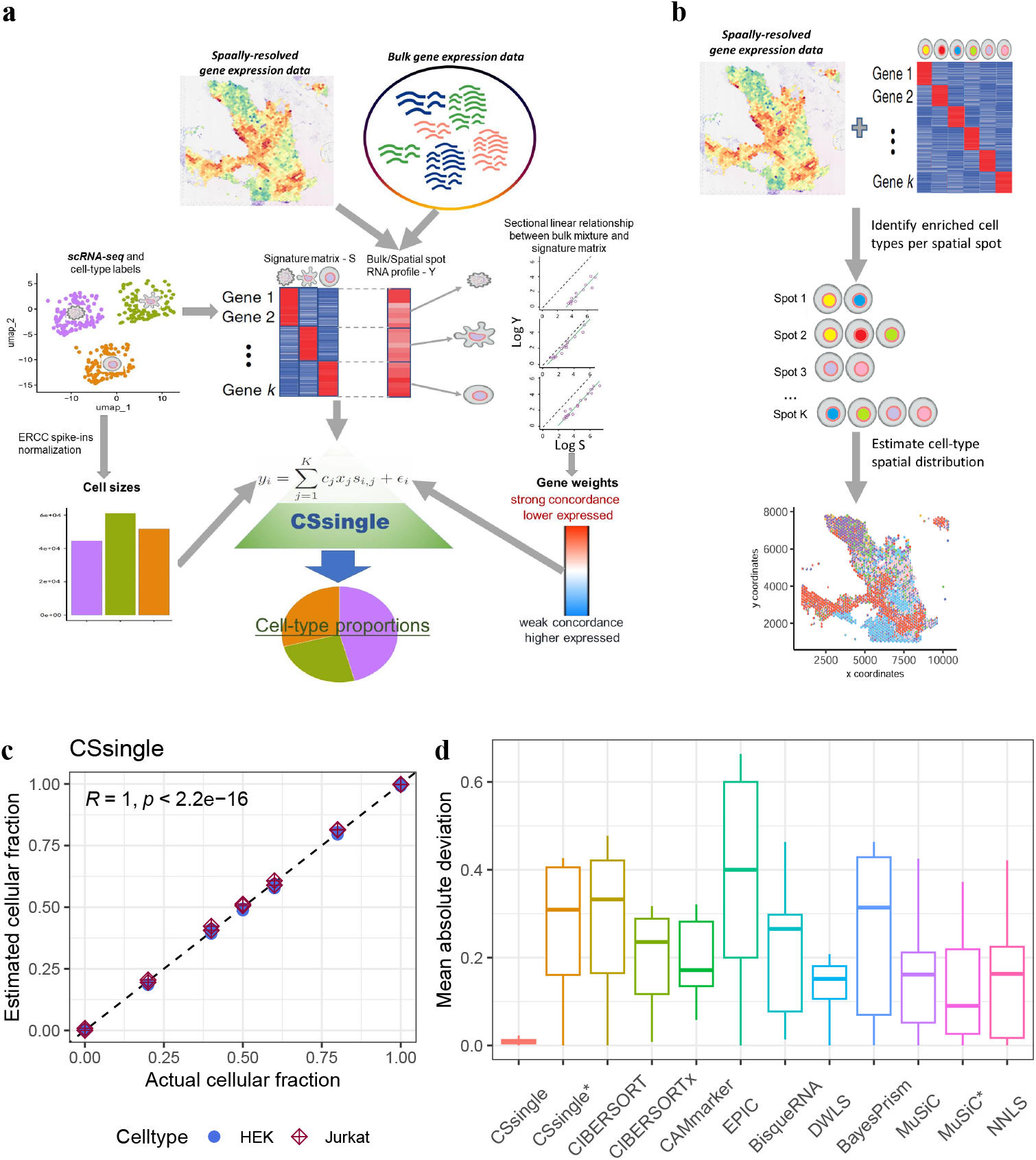
Schematic representation of the CSsingle workflow and performance validation. **a** CSsingle decomposes spatial and bulk transcriptomic data into a set of predefined cell types using the scRNA-seq or flow sorting reference. Within CSsingle, the cell size coefficients are estimated by using ERCC spike-in controls which allow the absolute RNA expression quantification. CSsingle is a robust deconvolution method based on the iteratively reweighted least squares approach. CSsingle assigns lower weights to genes with higher expression and/or weaker concordance between individual mixtures and the signature matrix to mitigate the influence of highly expressed genes in the least squares fitting procedure and enhance cross-platform performance. **b** CSsingle enables fine-scale analysis by identifying enriched cell types at single-spot resolution for spatial transcriptomic data. **(c**,**d)** Application of CSsingle to the deconvolution of bulk mixtures of HEK and Jurkat cells. **c** The plot illustrates the estimated cell type proportions by CSsingle compared to the actual cell type proportions. Reported ‘R’ corresponds to Pearson’s correlation coefficient and *p*-values indicate the significance of these correlations. **d** Boxplot depicting mean absolute deviation (mAD) between estimated and actual cell type proportions, with colors differentiating benchmark methods. The box encompasses quartiles of mAD, and whiskers span 1.5x the interquartile range.

To recapitulate the effect of cell size differences on deconvolution, the gene expression of marker gene *I* in cell type mixture *Y* is modeled within CSsingle by 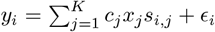 (Eq.1). Here, *c*_*j*_ denotes the cell size coefficient of cell type *j, x*_*j*_ represents the fraction of cell type *j* in mixture *Y, s*_*i,j*_ represents the column-normalized cell-type-specific expression level of marker gene *i* within cell type *j*, and *ϵ*_*i*_ models measurement noise and other possible un-modeled factors. Within CSsingle, cell size coefficients derived from scRNA-seq data were estimated by using a library of 96 external RNA spike-in controls developed by the External RNA Controls Consortium (ERCC).^33, 34^ This is achieved by leveraging ERCC spike-in controls as technology-independent standards for absolute RNA quantification, ensuring that cell size estimates are robust to technical variability. Our approach for inferring cell sizes using ERCC spike-in controls relies on two assumptions (i) the same amount of spike-in RNA is added to each cell; and (ii) technical effects should affect the spike-ins in the same way as they do the endogenous genes.^36, 37^ Under these assumptions, the ERCC spike-ins can act as technology-independent controls for cell size quantification.

Furthermore, when constructing a signature matrix from scRNA-seq data for deconvolution, it is important to take into account the technical and biological variations between the cell type mixture and the signature matrix, as previously highlighted.^23^ To appropriately use single-cell-derived cellular information to deconvolve bulk data, we designed a novel weighting scheme to solve cell type deconvolution as a minimization problem via 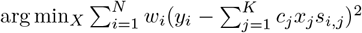 (Eq.2). This scheme aims to appropriately adjust the contribution of each gene, with the goal of enhancing cross-platform performance and mitigating the influence of highly expressed genes in the least squares fitting procedure.

CSsingle is a robust deconvolution method based on the concept of iteratively reweighted least squares (IRLS). To initialize an efficient and robust set of weights to solve the minimization problem, we relied on an important property of marker genes: there exists a sectional linear relationship between individual cell type mixtures and the signature matrix (Fig.2a). To illustrate this, we examined two fundamental assumptions underlying deconvolution methods: (i) expression profiles from each cell type are linearly additive (Eq.1), and (ii) cell-type-specific genes are exclusively or restrictively expressed in only one cell type within a cell type mixture. In an ideal scenario, genes exclusively expressed in a single cell type exhibit a strictly linear relationship with their expressions in a cell type mixture. Then Eq.1 can be expressed as log *y*_*i*_ = *u*_*j*_^*^ + log *s*_*i,j*_^*^ (Eq.3), where *u*_*j*_^*^ = log(*c*_*j*_^*^ *x*_*j*_^*^) and marker gene *i* is exclusively expressed in cell type *j*^*^. This equation highlights the “sectional linear relationship” between individual cell type mixtures and the signature matrix, allowing us to identify this key property of marker genes. Specifically, for each cell type *j*, we employed a linear regression model to fit the cell type mixture *Y* and each cell-type-specific GEP. This model was represented as log *y*_*i*_ = *u*_*j*_ + log *s*_*i,j*_ with the slope of exactly one, where *i* ∈ ℳ^*j*^ and ℳ^*j*^ is a finite set of marker genes for cell type *j*. Therefore, we determined the estimate *u*_*j*_ for each cell type *j* using the respective fitted linear regression model. Once *u*_*j*_ is estimated, CSsingle defines 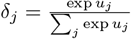, representing the estimated proportion of RNA content derived from cell type *j* in *Y*. In this formula, exp *u*_*j*_ depends on *c*_*j*_*x*_*j*_ which represents the estimated RNA content derived from cell type *j* in the mixture *Y*. We then created an estimated cell type mixture based on *δ*_*j*_ as *Y* ^*^ = (*t*_1_, *t*_2_, …, *t*_*N*_)^*T*^ where 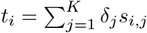. Finally, we estimated the initial weight for each gene *i* by considering both the difference between *y*_*i*_ and *t*_*i*_, and the magnitude of *t*_*i*_. Specifically, CSsingle downweights genes that have a large difference between *Y* and *Y* ^*^, a.k.a. weak concordant genes between *Y* and *Y* ^*^ Down-weighting weak concordant genes is vital since this kind of genes are sensitive to the technical and biological differences between the bulk and reference data. This approach offers a promising perspective for addressing technical and biological variations between mixture and single-cell reference data. The detailed method for further solving the minimization problem through IRLS is provided in the Methods section.

To reveal and employ the important “sectional linear” property of marker genes in real data by CSsingle, we first examined the linear relationship of marker genes in the aforementioned mixtures of HEK and Jurkat cells on a per-cell-type basis. We used mixtures of pure HEK or Jurkat cells to generate cell-type-specific GEPs to construct the signature matrix. Within the signature matrix, cell-type-specific marker genes were identified by using likelihood-ratio test. All bulk, spatial and single-cell data sets used in this study are summarized in Table 1. In Fig.S1a, we specifically fitted a linear regression model with Eq.3 (log *y*_*i*_ = *u*_*j*_ + log *s*_*i,j*_) for each cell type (solid lines). Importantly, for the five mixtures with different proportions of HEK and Jurkat cells, we observed a strong linear relationship between the HEK-specific GEPs and the bulk mixture using HEK-specific marker genes (*R >* 0.9; Fig.S1a, top panels, Samples 04, 06, 08, 10 and 12). We also observed a strong linear relationship between the Jurkat-specific GEPs and the bulk mixture using Jurkat-specific marker genes (*R >* 0.8; Fig.S1a, bottom panels). As a result, in these five bulk mixtures, we did reveal the sectional linear relationship between individual bulk mixtures and the signature matrix.

**Table 1:**
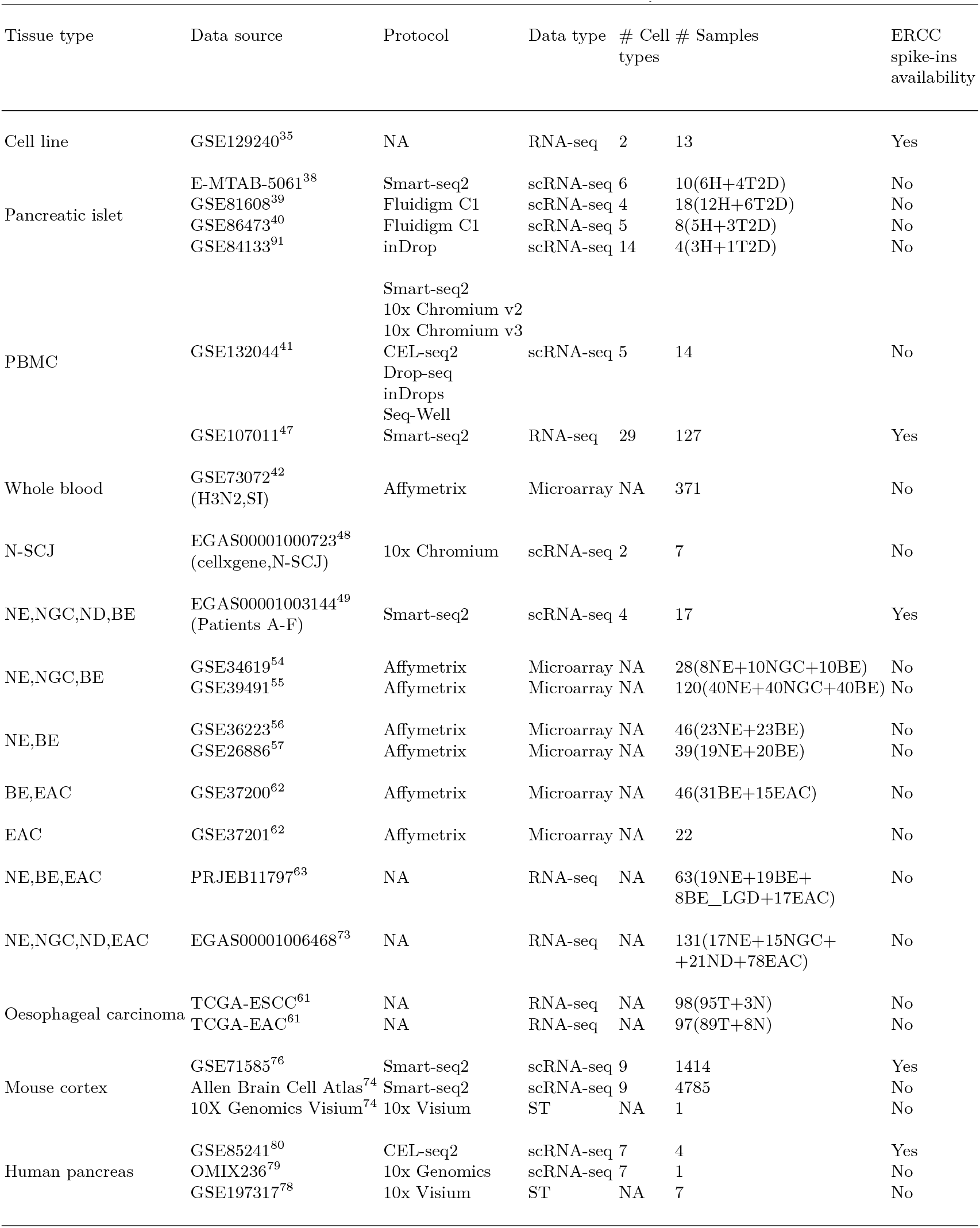
Details of the data sets used in the study.

To deconvolve the cell type proportions in 14 different bulk mixtures, CSsingle leverages the sectional linear relationship property to derive a robust and efficient set of initial weights (Fig.S1b). Additionally, CSsingle employs ERCC spike-ins to estimate cell sizes of HEK and Jurkat cells (Fig.1b) and integrates these size coefficients into its deconvolution framework. This methodology effectively accounts for variations in RNA content between cell types, significantly enhancing the accuracy of the estimation of cell type proportion in bulk samples (Fig. S2a,b). As shown in Fig.2c,d, CSsingle provides accurate estimates of cellular fractions in various mixtures of HEK and Jurkat, significantly outperforming 10 conventional methods.

### CSsingle improves cross-platform deconvolution

The pancreatic islet is a well-studied tissue with existing deconvolution methods.^14, 16, 18, 23^ We further benchmarked the performance of CSsingle using an artificial pancreatic islet data set, where the bulk data and reference data were generated on distinct platforms and under varying disease conditions. Specifically, we constructed the artificial bulk pancreatic islet mixtures by aggregating single cells of six well-characterized cell types (alpha, beta, gamma, delta, acinar, and ductal) for each healthy and diseased sample from Segerstolpe et al., where single cells were profiled by SMART-Seq2.^38^ We constructed the signature matrix using scRNA-seq data from healthy donors, sourced from two studies: four cell types (alpha, beta, delta, and gamma) from Xin et al.^39^ and five cell types (alpha, beta, delta, acinar, and ductal) from Lawlow et al.^40^ Both sets of single cells were profiled by Fluidigm C1. In this context, the signature matrix was built by integrating single cells from the two studies to obtain a more complete picture of the diverse cell types that make up the tissue. For each cell type, we observed that their specific GEPs within the signature matrix were well-correlated with the healthy or T2D bulk mixture by applying linear regression to their marker genes in log-scale via Eq.3 (*p <* 0.05; Fig.S3a and Fig.S4a). This analysis revealed that the bulk mixtures remained sectional linear with the signature matrix, despite being derived from different sequencing methods and different states of the disease. However, considerable variations were observed for many genes. We therefore hypothesized that these observed variations were, to some extent, shaped by technical and biological differences between the bulk mixture and signature matrix derived from different sources. To take advantage of the good concordant genes, we generated a set of initial coefficients by up-weighting marker genes with lower expression and strong concordance and down-weighting genes with higher expression and weak concordance between the bulk mixture and signature matrix (Fig.S3b and Fig.S4b). The results showed that CSsingle achieved higher accuracy in terms of three metrics (root mean square deviation, mean absolute deviation, and Pearson correlation) over all other methods (Table S1). Although the signature matrix was only built from healthy samples, we found that CSsingle proved adept at accurately estimating the fractions of major and minor cell types for both healthy and T2D bulk mixtures (Fig.3a, and Fig.S5). Specifically, only CSsingle, DWLS, and BayesPrism could accurately estimate the proportion of ductal cells; other methods consistently underestimated its presence.

**Figure 3.**
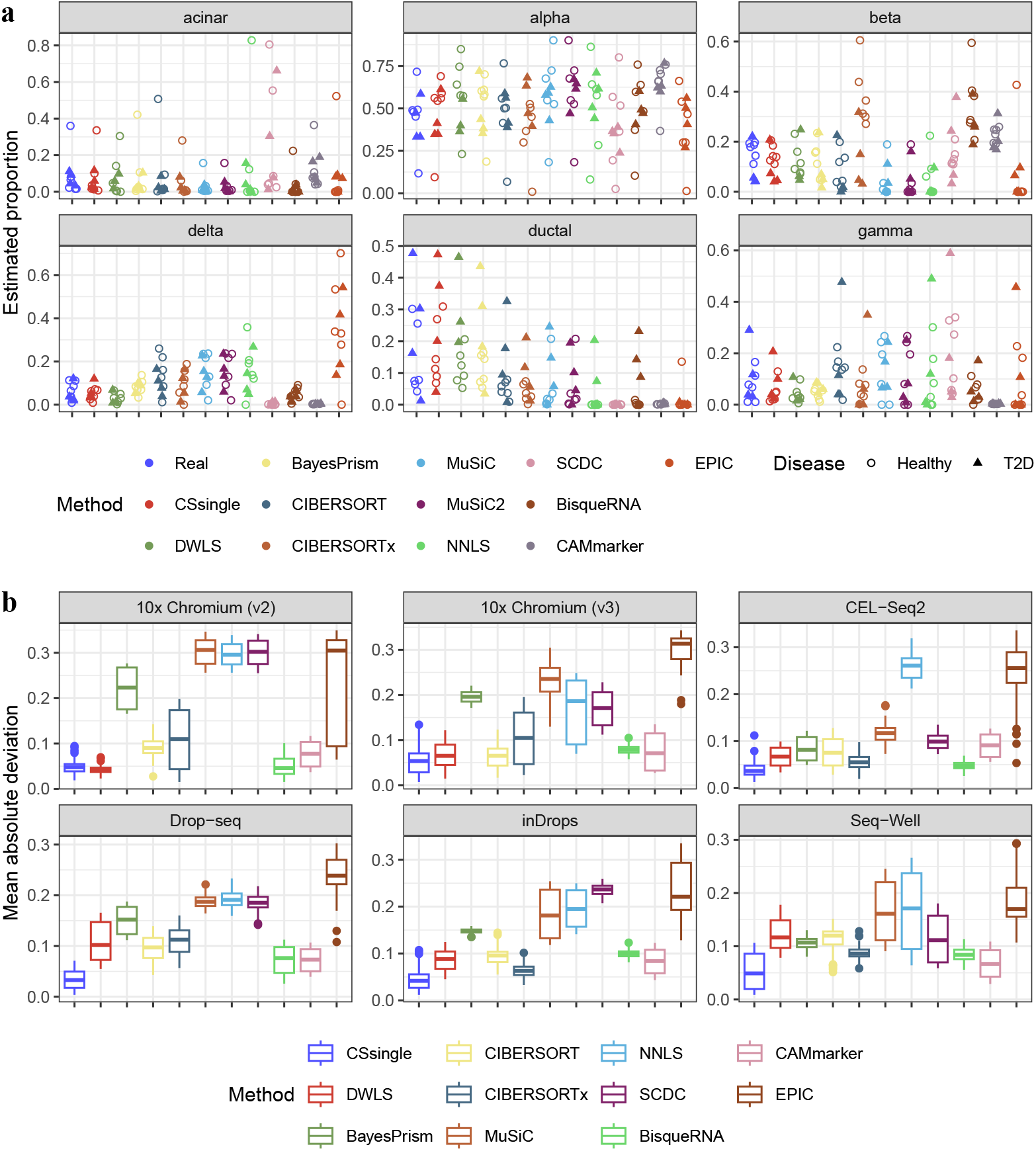
Decomposition benchmark in human pancreatic islet and PBMC. **a** Jitter plots displaying true and estimated cell type proportions in pancreatic islet. Each color represents a benchmarked method. Healthy subjects are denoted as dots while T2D subjects are denoted as triangles. **b** Decomposition benchmark of human PBMC using scRNA-seq reference data derived from six distinct scRNA-seq methods (10x Chromium v2, 10x Chromium v3, CEL-seq2, Drop-seq, inDrops, and Seq-Well).

ScRNA-seq has become a pivotal tool for profiling cellular heterogeneity, with rapid development of new and improved laboratory protocols and associated transcript quantification tools. Thus, there is a need to assess the performance of deconvolution methods for single-cell reference data generated from different laboratory protocols. To benchmark different deconvolution methods in this challenging but realistic scenario, we considered a benchmark data set introduced by Ding et al.,^41^ who were among the first to systematically examine how well scRNA-seq methods captured biological information. This data set profiled human peripheral blood mononuclear cells (PBMCs) samples employing two low-throughput plate-based methods, Smart-seq2 and CEL-Seq2, alongside five high-throughput methods: 10x Chromium v2, 10x Chromium v3, Drop-seq, Seq-Well and inDrops. SMART-Seq2 prepares full-length libraries, which is essentially identical to the process used in bulk RNA-seq. We created artificial bulk RNA-seq data using single cells from PBMCs profiled by the SMART-Seq2 protocol. Subsequently, we built signature matrices using scRNA-seq data derived from six distinct scRNA-seq methods (10x Chromium v2, 10x Chromium v3, CEL-seq2, Drop-seq, inDrops, and Seq-Well), all of which incorporate UMIs. We compared the Pearson correlation and mean absolute deviation between actual and estimated cell type proportions across all 11 methods. CSsingle achieved the best accuracy, while other methods performed substantially worse (Fig.3b, Fig.S6 and Table S2). The results demonstrate the versatility and robustness of CSsingle in handling reference data derived from various single-cell sequencing methods.

Finally, to assess the effectiveness of our initialization strategy employing the sectional linear property, we also introduced a comparable step within CSsingle, where initial coefficients were estimated with constant gene weights. The results indicated a significant performance improvement when CSsingle up-weighted strong concordant genes and down-weighted weak concordant genes, as opposed to the initialization strategy using constant gene weights (Fig.S7).

### CSsingle accurately estimates neutrophil-to-lymphocyte ratio from whole blood samples

We further benchmarked these bulk decomposition methods using real clinical samples. Specifically, we analyzed bulk transcriptomic data from a study involving influenza challenge in healthy adults aged 18-45, who were vaccinated with the A/Wisconsin/67/2005 (H3N2) strain. Genome-wide gene expression was evaluated in their peripheral blood using Affymetrix microarrays before the challenge and on days 2-7 after the challenge.^42^ Prior studies have shown that neutrophil-to-lymphocyte ratio (NLR) was associated with disease severity and mortality for influenza and Covid-19 patients.^43–45^ Thus, the ability to accurately deconvolve neutrophils and lymphocytes in blood samples has significant implications in clinical applications.

To evaluate the performance of deconvolution methods, we assessed the concordance between their estimates of neutrophil and lymphocyte proportions and standard laboratory-derived measurements, with the latter obtained from a pre-existing study.^46^ These laboratory-derived measurements were recorded daily from day 1 to day 7, along with a baseline measurement taken before inoculation. For deconvolution, we used the PBMC data set from an RNA-seq study^47^ as the reference. This reference data set contained profiles for 110 immune cells, grouped into three major cell populations: neutrophils, lymphocytes, and myeloids. Cell size estimates were obtained using ERCC spike-in controls, and a Kruskal-Wallis rank sum test revealed significant differences in cell sizes between the three cell populations (*p <* 0.0001; Fig.S8).

We then compared the accuracy of all 12 methods in deconvolving neutrophils and lymphocytes using blood samples from symptomatic infected (SI) adults, who exhibited noticeable signs of infection. Among the tested methods, CSsingle demonstrated the highest concordance with the laboratory measurements (Fig.S9-S10). While most methods consistently underestimated neutrophil proportions and overestimated lymphocyte proportions, CSsingle was able to correct this systematic bias. Notably, the CAMmarker tool, in contrast to other methods, underestimated both neutrophil and lymphocyte proportions (Fig.S9-S10).We further benchmarked temporal changes in the NLR, given its strong association with the progression of influenza, as previously reported.^46^ The temporal alterations in the NLR estimated by CSsingle showed excellent agreement with laboratory-derived measurements (*R* = 0.93, *p <* 0.001; Fig.4). Although other methods also displayed positive correlations between the estimated NLRs and laboratory measurements (*R >* 0.8, *p <* 0.05; Fig.4), their NLRs were consistently underestimated. These findings demonstrated that CSsingle accurately estimated neutrophil and lymphocyte proportions, as well as the NLR, from whole blood samples, highlighting its potential utility in clinical applications.

**Figure 4.**
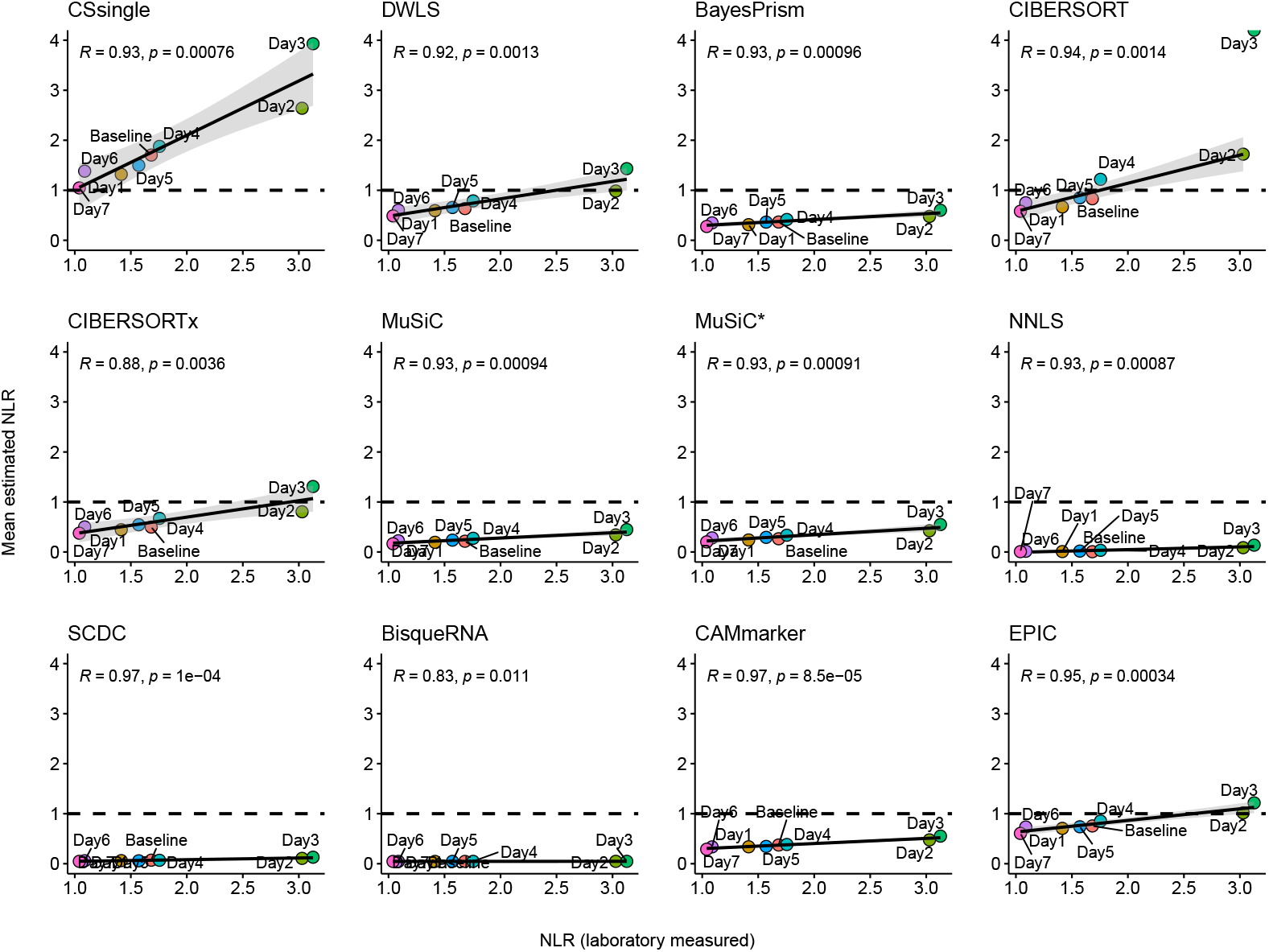
Correlation between laboratory-measured and estimated NLR in symptomatic infected (SI) group of influenza H3N2. Data points are labeled by days post-inoculation, with the baseline representing pre-inoculation and day 1 indicating the inoculation day. The ‘R’ value represents Pearson’s correlation coefficient, while *p*-values quantify the statistical significance of the observed correlations.

### CSsingle differentiates epithelial compartments spanning the gastroesophageal junction

We extended our analysis to the epithelial compartments spanning a complex anatomic area: the gastroesophageal junction. The native esophagus and gastric cardia are characterized as squamous and columnar, respectively. The squamous-columnar junction (SCJ) delineates the boundary between esophageal and gastric epithelial cells.^48^ To characterize the squamous and columnar compartments of the SCJ, we built a signature matrix using a scRNA-seq data set derived from Barrett’s esophagus (BE), normal esophagus (NE), normal gastric cardia (NGC) and normal duodenum (ND).^49^ Individual cells were classified into four epithelial cell populations: squamous, gastric columnar, intestinal columnar, and mosaic columnar (Fig.5a). To achieve this categorization, the cellular components were first characterized by two cell types: squamous cells, identified by their marker genes TP63 and KRT5, and columnar cells, identified by the marker gene KRT8^48^ (Fig.5b). The columnar cells were further categorized into gastric, intestinal, and mosaic columnar cells. Gastric and intestinal columnar cells were identified based on restricted expression of markers such as PGC, MUC5AC, and TFF1^50, 51^ for gastric columnar cells and OLFM4, GPA33, and TFF3^48^ for intestinal columnar cells. The mosaic columnar cells (MCCs) were defined by the co-expression of gastric and intestinal markers. We estimated cell sizes by using ERCC spike-in controls, and found that a statistically significant difference exists between cell sizes of four epithelial cell populations (*p <* 0.0001, Kruskal-Wallis rank sum test; Fig.5c).

**Figure 5.**
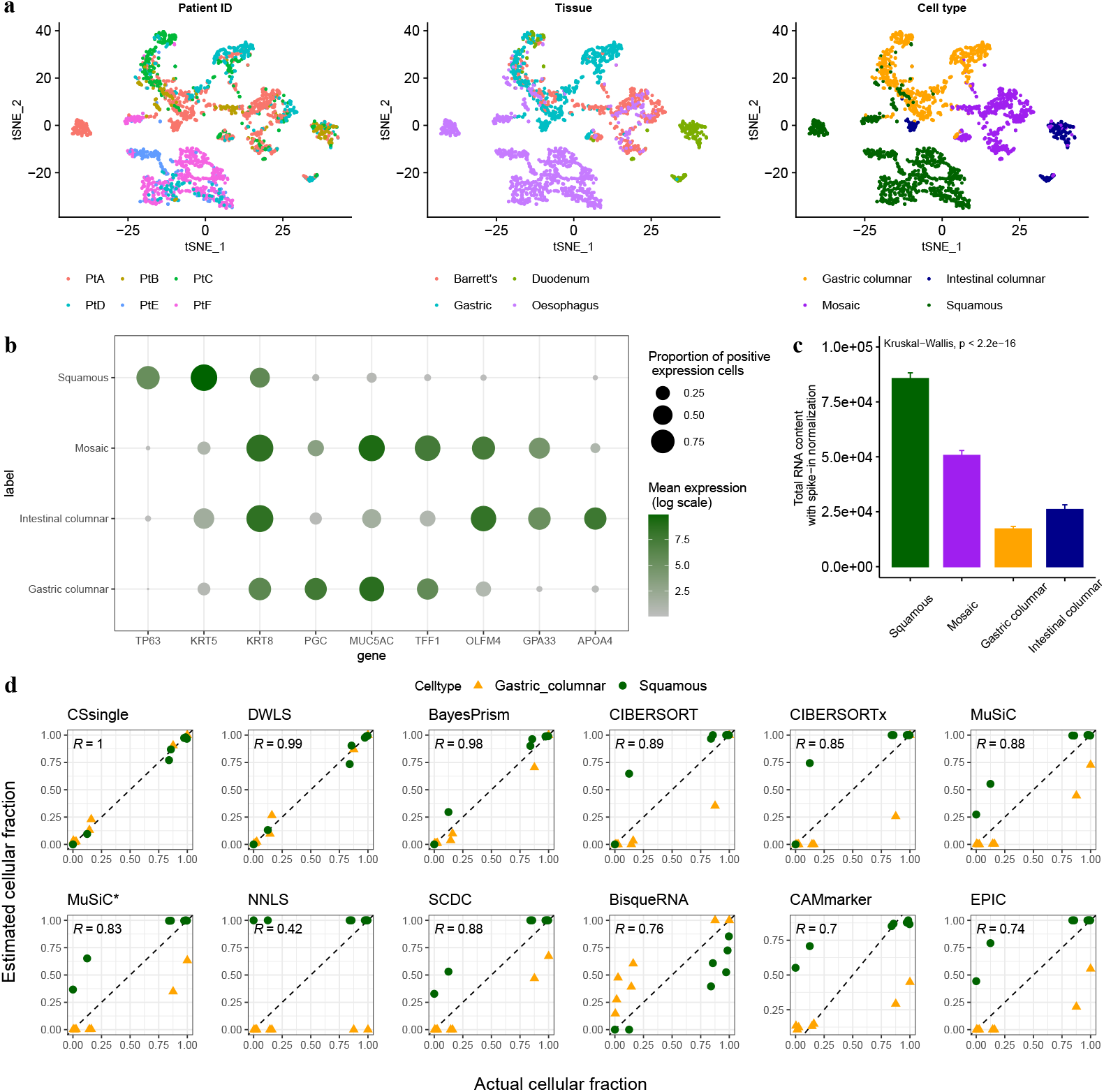
Decomposition benchmark in human normal squamous-columnar junction (N-SCJ) tissue. **a** t-SNE projection of scRNA-seq data from samples of normal squmous esophagus (NE), normal gastric cardia (NGC), normal intestinal (ND) and Barrett esophagus (BE), and color coded by the patient ID (left panel), tissue (middle panel) and cell type (right panel). **b** Bubble chart of selected marker gene expression among different cell types. **c** Cell sizes estimated using ERCC spike-in controls for each cell type. Error bars represent standard deviation. Kruskal-Wallis rank sum test was used to test whether the estimated cell sizes of four epithelial cell populations are statistically different. **d** Decomposition benchmark on the simulated bulk data of N-SCJ. The plot compares estimated versus actual cell type proportions, with colors representing squamous (green) and gastric columnar (orange) cell types. The ‘R’ value represents Pearson’s correlation coefficient, while *p*-values quantify the statistical significance of the observed correlations.

To evaluate CSsingle’s performance in analyzing the squamous and columnar compartments of normal SCJ (N-SCJ), we created seven artificial bulk mixtures using a scRNA-seq data set,^48^ derived from N-SCJ samples of seven healthy subjects. For deconvolution, the signature matrix constructed using scRNA-seq data shown in Fig.5a was refined to include only the squamous and gastric columnar cell populations. Notably, the squamous cells are larger in size compared to gastric columnar cells (Fig.5c). By applying cell size correction, CSsingle precisely compensates for variations in RNA content between the two cell types, leading to more accurate estimates of their proportions in bulk samples of N-SCJ (Fig.S11). Furthermore, our results demonstrated that the cellular fractions estimated by CSsingle exhibited a perfect correlation with the actual cellular fractions, outperforming DWLS and BayesPrism, while other methods showed significantly inferior performance (Fig.5d). Specifically, most of the other methods exhibited a consistent tendency to overestimate the cellular fractions of squamous cells with larger cell size while underestimating those of gastric columnar cells characterized by smaller cell size.

### CSsingle identifies mosaic columnar cells from bulk data of Barrett’s esophagus and esophageal adenocarcinoma

Barrett’s metaplasia, characterized by the transformation of squamous epithelium into columnar epithelium, serves as a precursor lesion for the development of esophageal adenocarcinoma (EAC).^52^ In the case of BE, metaplasia occurs at the SCJ.^53^ Prior studies have reported that individual columnar cells of BE show a gastric-intestinal mosaic phenotype as defined by the co-expression of gastric and intestinal markers, while this mosaic phenotype is absent in normal gastric or intestinal tissues.^48^ Within our previously established signature matrix, we specifically identified a cluster of mosaic columnar cells (MCCs) exhibiting co-expression of gastric markers (PGC, MUC5AC, and TFF1) alongside intestinal markers (OLFM4, GPA33, and TFF3). In particular, this co-expression pattern was not observed in the normal gastric or intestinal cell clusters (Fig.S12a).

We challenged CSsingle and 11 other deconvolution methods using 233 epithelium samples generated from NE, NGC, and BE tissues derived from four studies^54–57^ (Table 1). Given that the NE and NGC tissues are composed of squamous and columnar epithelium, respectively, CSsingle, DWLS, CIBER-SORT, CIBERSORTx, and SCDC correctly identified the predominant cell type as squamous in the NE and gastric columnar in the NGC epithelium (Fig.6a and Fig.S12b). In contrast, other methods either underestimated the squamous cells in the NE or the gastric columnar cells in the NGC (Fig.S12b). Notably, CSsingle, DWLS, CIBERSORT, and CIBERSORTx also detected a significant increase in the proportion of MCC in BE samples compared to those in NE and NGC (*p <* 0.05, one-sided Wilcoxon-test relative to NE and NGC, respectively; Fig.S13). These findings align with clinical and histological observations,^48, 58^ reinforcing the presence of gastric and intestinal phenotypic mosaicism in Barrett’s metaplasia.

**Figure 6.**
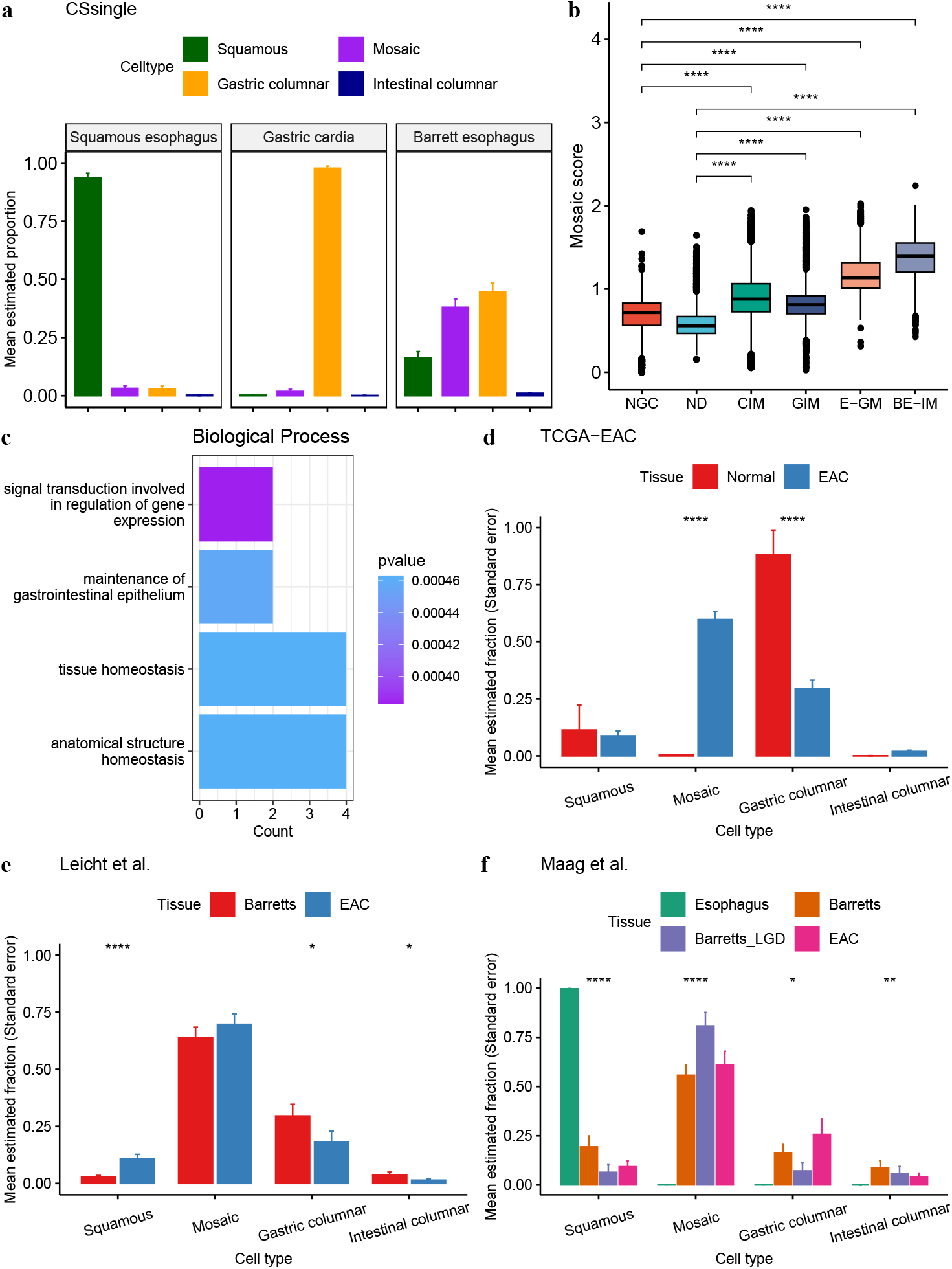
The mosaic columnar cells predominantly present in Barrett’s esophagus and esophageal adenocarcinoma. **a** Comparison of decomposition estimates by CSsingle for 233 epithelium samples derived from normal squmous esophagus (NE), normal gastric cardia (NGC) and Barrett esophagus (BE). **b** Comparison of mosaic scores for columnar cells across esophageal (E-GM, BE-IM) and gastric (CIM, GIM) intestinal metaplasia, as well as normal gastric cardia (NGC) and normal intestinal (ND) tissues. **c** GO enrichment analysis for the top 25 markers of mosaic columnar cells. **d**,**e**,**f** The relative abundance of four epithelial cell populations for normal and diseased tissues from (**d**) TCGA-EAC, (**e**) Leicht et al.^62^ and (**f**) Maag et al.,^63^ colored by tissue of origin. Error bars represent standard deviation.

To further characterize phenotypic mosaicism in esophageal and gastric intestinal metaplasia compared with normal tissues, we investigated scRNA-seq data from these tissues.^48^ We calculated the mosaic score, defined as the mean of log-normalized expressions of the top 25 markers for MCC (Table.S3). Our analysis indicated that the mosaic scores in columnar cells of esophageal and gastric intestinal metaplasia were significantly higher than in normal gastric and intestinal tissues (Fig.6b). Assessing the top 25 markers of the MCC through gene ontology (GO) enrichment analysis revealed strong associations with processes involved in signal transduction, regulation of gene expression, maintenance of gastrointestinal epithelium, tissue homeostasis, and anatomical structure homeostasis (Fig.6c). While these biological processes are critical for adaptation in response to chronic inflammatory or injurious stress, their dysregulation is a common feature in early neoplastic progression.^59, 60^ Collectively, these findings indicate that esophageal intestinal metaplasia is characterized by upregulation of epithelial markers specific to novel cell identities (e.g., MCC), driven by transcriptional reprogramming in response to chronic inflammatory or injurious stress. This adaptive mechanism enables epithelial plasticity to meet functional demands while simultaneously increasing susceptibility to neoplastic transformation.

Esophageal cancer is classified by histology as squamous cell carcinoma (ESCC) or adenocarcinoma (EAC), both of which share an anatomic site.^52^ We applied CSsingle to dissect the epithelial composition from the transcriptomes of 184 bulk oesophageal tumors and 11 adjacent normal tissues profiled by TCGA.^61^ In this TCGA data set, about 70% tumor samples were collected from the distal esophagus. For the deconvolution of tumor-adjacent normal tissues, we observed that gastric columnar cells were consistently dominant in both the EAC and ESCC cohorts, followed by squamous cells, while the presence of mosaic and intestinal columnar cells was rare (Fig.S14a). Additionally, no significant differences were observed in the proportions of all four cell populations between the EAC and ESCC cohorts in tumor-adjacent normal tissues (Fig.S14a). For the deconvolution of ESCC tumors, we found that squamous cells were predominant (mean estimated proportions *>* 50%; Fig.S14b). In contrast, the MCCs were most abundant in EAC tumors (mean estimated proportions *>* 50%; Fig.S14b and Fig.6d), which shared the same characteristics as BE.

Furthermore, analysis of two additional published RNA-seq data sets^62, 63^ revealed that MCC remained predominant in Barrett’s esophagus (with or without low-grade dysplasia (LGD)) and EAC, but was rare in normal esophagus (Fig.6e,f). These findings highlight the potential involvement of MCC in the progression from metaplasia to dysplasia and ultimately to invasive carcinoma. This aligns with previous studies,^48, 52^ which demonstrated that the metaplastic epithelium in EAC progresses through a stage of dysplasia to invasive cancer. Our results demonstrate significant differences in epithelial composition between ESCC and EAC tumors, with MCCs predominantly present in BE and EAC but notably absent in ESCC and normal esophagus, highlighting their specific association with the pathogenesis of EAC.

### Mosaic-related signatures predict immunotherapy response and patient survival in esophageal adenocarcinoma

Given the association between MCC and the pathogenesis of EAC, we explored their role in immunotherapy response. Epithelial-to-mesenchymal plasticity (EMP) refers to the reversible transition of cancer/epithelial cells between epithelial-like and mesenchymal-like states.^64, 65^ EMP encompassing epithelial-to-mesenchymal transition (EMT), has been implicated in immune escape and resistance to cancer immunotherapy.^66–71^ We assessed the EMT status in four epithelial compartments (squamous, gastric columnar, intestinal columnar, and mosaic columnar cells) by calculating EMT scores using 145 epithelial marker genes and 170 mesenchymal marker genes from a previous study.^72^ The EMT scores were calculated as the average log-normalized expression of mesenchymal or epithelial marker genes. In MCC, epithelial marker expression is significantly elevated compared to the other three epithelial compartments (Fig.S15a, top panel), while no significant change in mesenchymal marker expression is observed (Fig.S15a, bottom panel). However, “TGF-*β* receptor signaling in EMT” pathway is significantly enriched in MCCs (Fig.S15b). The results indicate that MCCs exhibit a unique pre-EMT characteristics, where epithelial cells are preparing for a potential transition but have not yet initiated the full EMT program. Moreover, we observed that intestinal stem cell-related markers are significanlty up-regulated in MCCs compared to gastric columnar and squamous cells, indicating a potential progenitor-like functionality or retained plasticity that may enable MCCs to contribute to epithelial regeneration (Fig.S15c and Fig.S16).

Currently, there are fewer useful predictive biomarkers to identify EAC patients who would benefit the most from immune checkpoint inhibitors (ICI) treatment combined with or without concurrent chemotherapy.^73^ To investigate the role of MCCs in response to ICI in EAC, we applied CSsingle to dissect the epithelial composition using esophageal transcriptome data from the phase I/II LUD2015-005 trial.^73^ This clinical trial involved patients with inoperable esophageal cancers, predominantly EAC, who received ICI immunotherapy for a four-week window (ICI-4W) before immunochemotherapy (ICI + CTX). Patients with irRECIST outcomes of complete or partial responses were classified as responders, while others were non-responders. Additionally, patients achieving 12 months of progression-free survival were considered to have long-lasting clinical benefit (CB), while the remaining were classified as no clinical benefit (NCB).^73^ Specifically focusing on the epithelial compartments in inoperable EAC, we compared the cell composition estimates between EAC tumor samples and paired normal gastrointestinal (GI) tissue biopsies (esophagus, gastric and duodenum), all collected prior to ICI treatment (PreTx). Our analysis revealed a significant increase in the proportion of MCC in EAC compared to paired normal GI tissue biopies (Fig.S15d). Further analysis of changes in epithelial composition at three time points - before (PreTx), during (ICI-4W), and after (PostTx) treatment - revealed a progressively decrease in the proportion of MCC and a corresponding increase in squamous cell in patients who achieved CB after ICI treatment combined with chemotherapy (Fig.7a), suggesting a change in epithelial cellular dynamics linked to therapeutic response. This observation suggests that the reduction of MCCs alongside the expansion of squamous cells after ICI + CTX treatment may play an important role in mediating or reflecting the effectiveness of immunochemotherapy in EAC.

**Figure 7.**
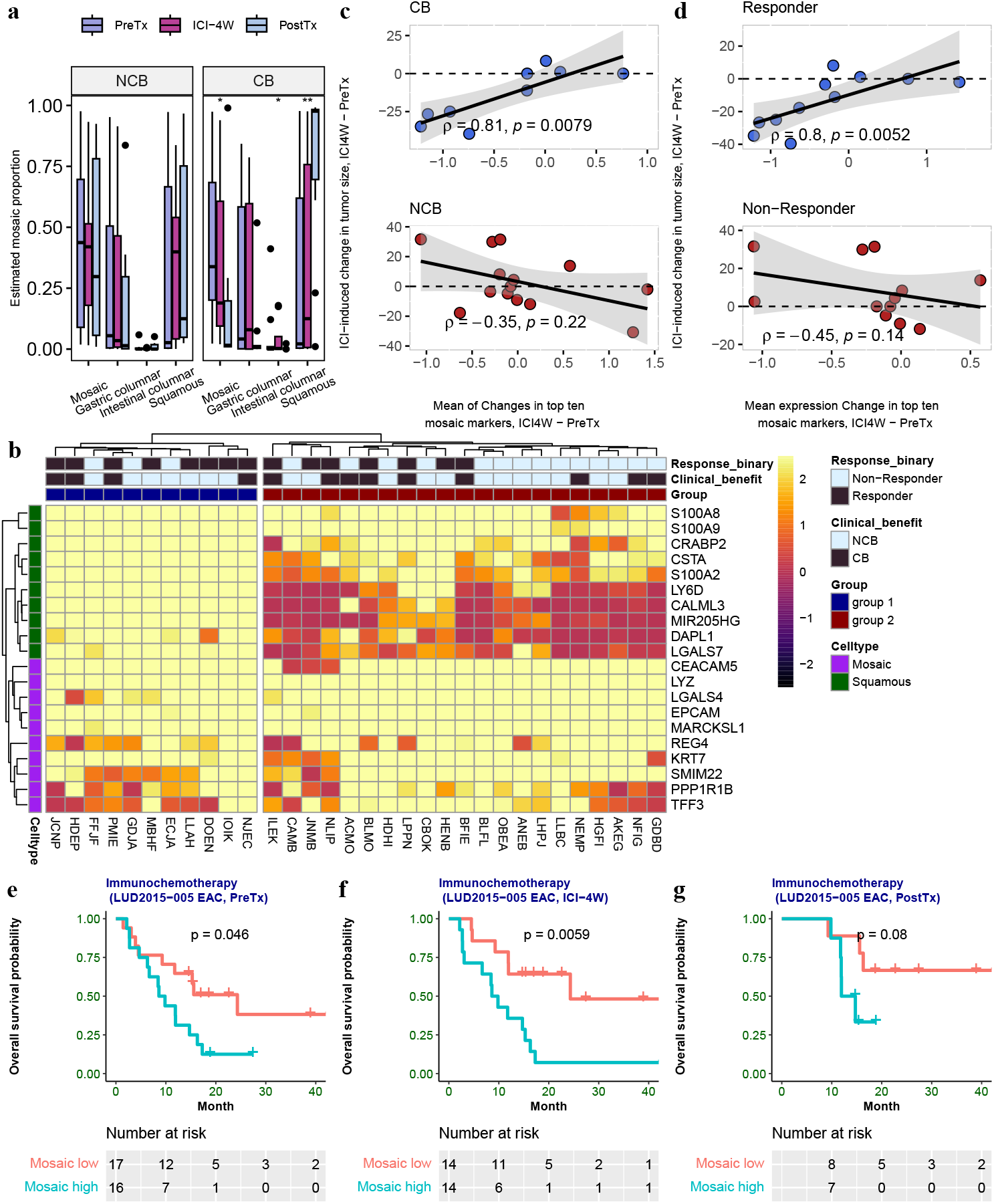
The mosaic-related signatures predict immunotherapy response and patient survival in esophageal adenocarcinoma. **a** Boxplots depicting CSsingle-estimated cell type proportions for patients with non-clinical benefit (NCB) and clinical benefit (CB), compared across pre-treatment (PreTx), during treatment (ICI-4W), and post-treatment (PostTx) phases. Statistical significance was assessed using the Kruskal-Wallis test. **b** Heatmap of log-TPM expression for top ten markers of MCCs and squamous cells in PreTx EAC tumors. Rows and columns were clustered using Euclidean distance and average linkage. **c**,**d** Scatter plots comparing ICI-induced mean expression changes of the top ten mosaic markers with ICI-induced tumor size changes in (**c**) CB vs. NCB patients and (**d**) responders vs. non-responders. Spearman correlation statistics are displayed. **e**,**f**,**g** Kaplan-Meier plots depict the overall survival (OS) of Mosaic-high and Mosaic-low groups from LUD2015-005 EAC patients at three time points (PreTx, ICI-4W, and PostTx), split by the cohort median. *p* value by log rank test.

Next, we evaluated whether PreTx expression of MCC and squamous markers could predict patient outcomes from immunochemotherapy. We performed hierarchical clustering of PreTx EAC tumor samples based on the expression profiles of the top ten MCC marker genes (REG4, TFF3, LGALS4, PPP1R1B, KRT7, EPCAM, CEACAM5, SMIM22, LYZ, and MARCKSL1) and the top ten squamous marker genes (S100A8, LY6D, MIR205HG, S100A9, S100A2, CSTA, DAPL1, CALML3, LGALS7, and CRABP2). Tumors were grouped into two clusters: group 1, characterized by higher squamous marker expression and lower MCC marker expression, and group 2, characterized by lower squamous marker expression and higher MCC marker expression (Fig.7b). Group 1 exhibited a higher response rate (8/11, 73%) compared to group 2 (7/21, 33%; Fig.7b). However, the CB rate in group 1 (4/11, 36%) was comparable to that in group 2 (9/21, 43%; Fig.7b). These results show that PreTx MCC and squamous marker expression predicts ICI response but not long-lasting CB.

We further examine whether changes in the expression of the MCC markers after four weeks of ICI treatment were associated with ICI outcomes. Analysis of the top ten MCC marker genes revealed that down-regulation of MCC signatures was associated with both the ICI response and the CB. Downregulation of the top ten MCC marker genes strongly correlated with ICI-induced tumor shrinkage in patients achieving CB (Fig.7c, top panel) or responding to ICI (Fig.7d, top panel). In contrast, no significant correlation was observed in patients without CB (Fig.7c, bottom panel) or ICI response (Fig.7d, bottom panel). These results indicate that changes in MCC marker expression during four weeks of ICI treatment may serve as a candidate biomarker for therapeutic efficacy, with potential utility in stratifying EAC patients likely to benefit from immunochemotherapy.

To investigate the prognostic role of the top ten MCC marker genes before, during, and after treatment, we stratified patients into mosaic-low and mosaic-high groups based on the cohort median at each of these time points. This classification effectively differentiated patient outcomes at each stage (Fig. 7e,f,g). A higher mosaic score was strongly associated with poorer overall survival (OS), highlighting its potential as a prognostic indicator of disease progression.

In summary, we revealed a dynamic relationship between MCCs and squamous cells during ICI treatment in EAC patients, suggesting MCC expression signatures as predictive and prognostic markers of ICI efficacy. Our findings reveal the critical role of MCC in the treatment of EAC and its potential as a biomarker to predict outcomes of immunochemotherapy, offering key insights into tumor epithelial plasticity to guide personalized immunotherapeutic strategies.

### CSsingle accurately estimates spatial distribution of cells from 10x Visium mouse brain cortex

With the rapid advancement of spatial transcriptomics (ST), we evaluated the ability of CSsingle to estimate the spatial distribution of cells using simulated ST data. To build the simulated ST data set, we initially used the CellTrek tool^74^ to align a well-annotated mouse scRNA-seq data set of the primary visual cortex from the Allen Brain Atlas with the corresponding 10x Visium ST data set derived from the region of the frontal cortex of mouse brain,^75^ thus generating a simulated ST data set with single cell resolution. Subsequently, we established 200 × 200 squares to serve as simulated spots and created spot-level expression profiles by aggregating gene expression data from multiple cells within each square. The coordinates of the first cell positioned within the square were designated as the spot’s location. To construct signature matrices, we used an independent scRNA-seq data set from the adult mouse primary visual cortex sequenced by Smart-seq2.^76^ Details of each data set are shown in Table 1.

ST is constrained by the limited detection of mRNA molecules within individual spatial spots, often due to low RNA capture efficiency. This sparsity obscures true gene expression signals, particularly for low-abundance transcripts, complicating tasks like deconvolution. CSsingle enhances ST deconvolution by detecting cell type enrichment at single-spot resolution rather than relying on pre-clustered regions, enabling finer-grained spatial mapping. The details are described in the “Methods” section.

We first applied CSsingle to estimate the cell type fractions of three main cell types in the cortex: GABAergic, glutamatergic, and non-neuronal cells, and compared its performance to that of the state-of-the-art ST deconvolution methods. The results demonstrated that the predictions of CSsingle were highly correlated with the actual cell type distributions (Pearson’s *R* = 0.98), outperforming seven other available methods (Pearson’s *R* = −0.02 − 0.89; Fig.8a and Fig.S17a). We further challenged CSsingle to predict the spatial distribution of the neuron sub-types. CSsingle’s prediction of the spatial distribution of neuron subtypes achieved the best performance, followed by SpatialDWLS^27^ and Seurat,^26^ all demonstrating a high level of consistency with the known locations of these subtypes across the cortex layers (Fig.8b and Fig.S17b). In contrast, the spatial distribution predicted by RCTD^29^ displayed ectopic mapping of L4 and L6 neurons to L5, while other four methods, including Redeconve,^31^ SpatialDecon,^30^ SPOTlight^28^ and SpatialDDLS^32^ failed to accurately reconstruct the layered structure. In general, CSsingle accurately mapped the three main cell types and nine subclusters in the cortex region, demonstrating its reliability in predicting cell type spatial distributions.

**Figure 8.**
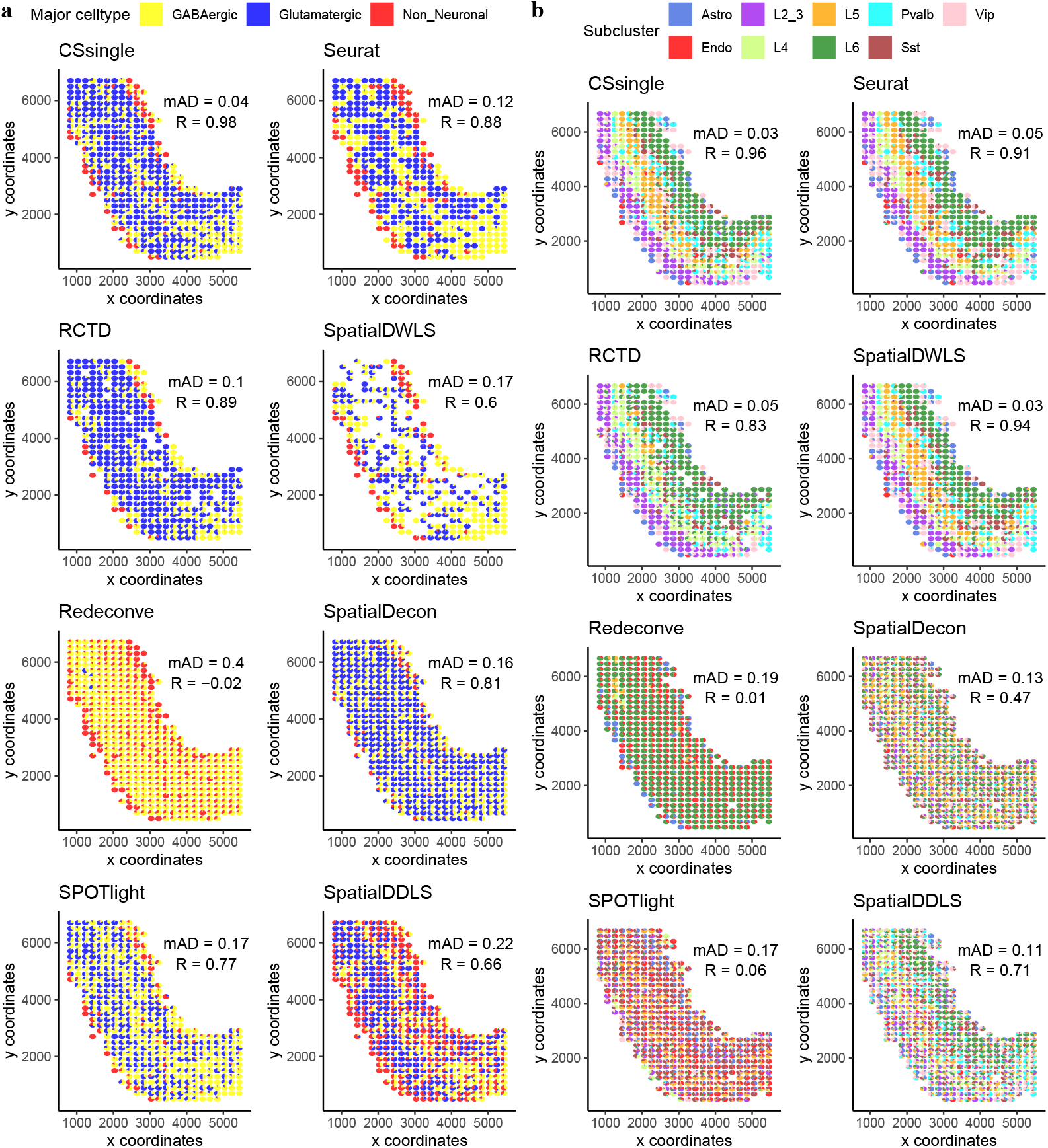
Decomposition benchmark in mouse brain tissue. Scatter pie plots showing the estimated proportions of major subtypes (**a**) and minor subtypes (**b**) at each spot.

In this study, we assessed whether the incorporation of cell size coefficients, estimated by ERCC spike-ins from single-cell reference data, improves the deconvolution accuracy of CSsingle in ST datasets. For comparison, parallel analyses were performed without cell size correction. Despite statistically significant differences in cell sizes between three main cell types and nine subclusters (p < 0.001, Kruskal-Wallis test; Fig.S18a,b), the deconvolution accuracy for CSsingle showed no significant difference with correction of cell size compared to uncorrected models (Fig.S18c,d). This contrasts with bulk RNA-seq, where size correction is critical. In 2D spatial transcriptomics, tissue sectioning can result in incomplete cells, as only a portion of a 3D cell is captured in the slice. This leads to partial RNA recovery, potentially underestimating gene expression and introducing noise. Furthermore, spatial complexity, such as partial cell overlaps and mixed cellular neighborhoods, overshadows RNA content biases in spatially resolved data.

### CSsingle reveals epithelial-to-mesechymal transition (EMT)/MET in acinar proliferation during human pancreas development

Previous research has demonstrated that during key stages of embryogenesis and organ development, cells in certain epithelia exhibit plasticity, transitioning between epithelial and mesenchymal states through the processes of epithelial-to-mesenchymal transition (EMT) and mesenchymal-to-epithelial transition (MET).^77^ Given this dynamic behavior, exploring the potential of CSsingle to identify and elucidate these transitions offers valuable opportunities to advance our understanding of these processes. Edward Olaniru and colleagues recently studied the human fetal pancreas at multiple developmental time points (12, 15, 18, and 20 postconception week [PCW]) using 10x Visium ST.^78^ We began by benchmarking CSsingle to evaluate its accuracy in estimating the spatial distribution of cell types using simulated ST data of the human pancreas. To construct the simulated ST dataset, we aligned the 10x Genomics scRNA-seq data of human adult pancreas^79^ with 10x Visium ST data from seven human fetal pancreas slides^78^ using CellTrek. We then generated 150 × 150 squares as simulated spots, preserving spatial context and cellular heterogeneity. To build the signature matrix, we used an independent scRNA-seq dataset of the human fetal pancreas at 12 PCW,^80^ generated using CEL-seq2. Benchmarking on seven simulated ST slides demonstrated that CSsingle achieved the highest correlation with actual cell-type distributions (Pearson’s *R* = 0.90), outperforming seven existing methods (Pearson’s *R* = 0.19̌0.88; Fig.S19). In particular, the correction for cell size (Fig.S20a) did not have a significant impact on precision (Fig.S20b), reinforcing its limited utility in spatial transcriptomics.

We next applied CSsingle to assign and decompose cell types in the real ST data set from the human pancreas.^78^ Since ground truth cell type labels are unavailable for real ST data, we assessed CSsingle’s performance by analyzing marker gene expression patterns and correlating its predictions with cell type enrichment scores. The results revealed that CSsingle accurately predicted the spatial distribution of cell types, with its predictions aligning well with known marker gene expression patterns (Fig. S21-S27) and exhibiting strong correlation with cell type enrichment scores (Fig.S28). To investigate spatial-temporal dynamics in cell type composition during pancreas development, we analyzed cell type abundances across four developmental stages (Fig.9a). Our analysis reveals that mesenchymal cells were predominant at 12 and 15 PCW, while acinar cells became the dominant population at 18 and 20 PCW. The abundance of mesenchymal cells decreased sharply from 24% at 15 PCW to 17% at 18 PCW, while the abundance of acinar cells increased from 19% at 15 PCW to 33% at 18 PCW (Fig.9b). The observed reduction in mesenchymal cells and expansion in acinar cells may reflect differences in proliferation rates or development loss of mesenchymal cells. Consistent with our findings, previous studies have shown that mesenchymal-epithelial interactions promote the proliferation and migration of the tissue epithelium.^81, 82^

**Figure 9.**
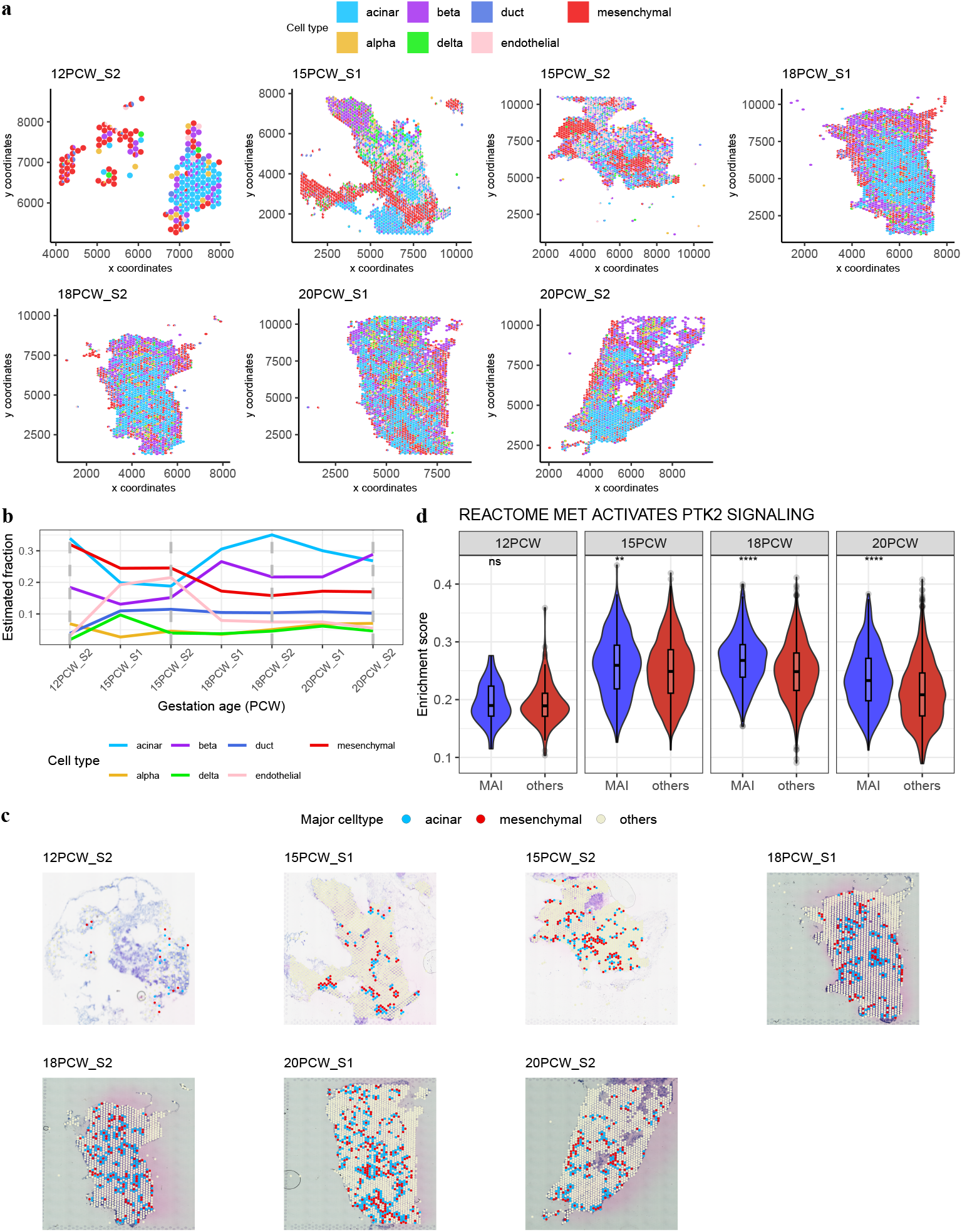
Spatial and molecular characterization of mesenchymal-acinar interactions (MAIs) estimated by CSsingle during pancreas development. **a** Spatial-temporal changes in CSsingle-estimated cell type composition during pancreas development. **b** Temporal changes in mesenchymal and acinar cell abundance during pancreas development. **c** CSsingle identified neighboring spots of mesenchymal and acinar cells for MAI investigation, revealing a significant increase at 18 and 20 PCW. **d** Reactome pathway analysis showed increased enrichment of MET-mediated activation of PTK2 signaling in MAIs compared to other spots, at 15, 18 and 20 PCW.

To further investigate the interaction between mesenchymal and epithelial cells, we used ST data to identify spatially adjacent spots that predominantly consist of mesenchymal or acinar cells, with a focus on mesenchymal-acinar interactions (MAI). These interactions were significantly elevated at 18 and 20 PCW (Fig.9c). To identify pathways associated with MAI, we performed a single sample gene set enrichment analysis (ssGSEA) using Hallmark and Reactome gene sets^83^ from the Molecular Signatures Database. Analysis of the Hallmark gene sets revealed a significant enrichment of the sets of target genes of EMT and MYC in MAI, with enrichment scores notably higher in MAIs compared to other spots at 15, 18, and 20 PCW (Fig.S29a,b). Furthermore, analysis of the Reactome pathways revealed that MET-mediated activation of the PTK2 (Focal Adhesion Kinase) signaling pathway was significantly enriched in MAIs compared to other spots at 15, 18, and 20 PCW (Fig.9d). These findings highlight the critical role of MAIs in the activation of pathways associated with the mesenchymal-epithelial transition, MYC-driven transcriptional regulation, cell adhesion and proliferation, highlighting their contribution to tissue remodeling during pancreatic development.

### Evaluation of signature matrix robustness

The utilization of a signature matrix not only improves computational efficiency but also significantly affects the accuracy of deconvolution. The construction of a signature matrix is a process of feature selection. To achieve this, we first identified differentially expressed genes between each cell type and all other cell types based on the thresholds of an FDR adjusted p-value of less than 0.01 and a *log*2 mean fold change greater than 0.25. Next, for each cell type, we ranked the differentially expressed genes by their p-values and selected the top *N* genes to build the signature matrix. In this study, the value of *N* ranged from 50 to 300, increasing by 50 at each step. We selected the optimal *N* for constructing a robust signature matrix by maximizing the Spearman correlation between the inferred and real bulk gene expression matrices (see Methods section). Traditionally, the stability of a signature matrix has been evaluated using the two norm condition number, and the signature matrix with the minimum condition number was retained.^15, 20^ Here, we explored whether these two strategies achieved consistent results by integrating CSsingle with a signature matrix generated from each. We systematically evaluated the deconvolution results across nine data sets, including HEK and Jurkat cell lines, pancreatic islets, PBMCs, and N-SCJ samples. The results showed that CSsingle exhibited comparable performance using both strategies for building the optimal signature matrix (Fig.S30), highlighting the robustness and flexibility of CSsingle in handling different matrix construction approaches.

Furthermore, since computational efficiency and deconvolution accuracy also depend on the step size used in generating the final signature matrix, we conducted an additional experiment. We assessed the runtime and deconvolution results as the step size decreased from 50 to 1 on five data sets. Across five data sets, we found that setting the step size to 1 did not necessarily lead to improved accuracy. However, setting the step size to 50 achieved comparable performance while significantly enhancing computational efficiency and robustness of deconvolution (Table S4).

## Discussion

Accurate decomposition of cell type mixtures is critical for studying cellular heterogeneity in clinical studies and spatial biology research. A common assumption prevalent in transcriptomic analysis, including cell type decomposition, is that most genes exhibit stable expression across cells, implying that RNA content remains constant across cell types. However, substantial evidence from many studies indicates frequent violations of this assumption.^84, 85^ Current bulk data deconvolution methods struggle to accurately decompose mixtures of cell types with significantly different cell sizes, which reflects variations in absolute RNA content between cell types. In this study, we demonstrated that cell size differences can significantly impact the accuracy of bulk data deconvolution. Bulk RNA-seq measures the average gene expression across a population of cells, which inherently includes contributions from cells of varying sizes. Larger cells contribute more RNA to the bulk sample, hence, by incorporating cell size coefficients, deconvolution algorithms can adjust for differences in RNA content between cell types, leading to more accurate estimates of their proportions in the bulk sample.

Here, we introduce CSsingle, a unified method developed to accurately decompose both bulk and spatial transcriptomic data into a predefined set of cell types, leveraging scRNA-seq or flow sorting reference data sets. Through comprehensive benchmarking and analysis of multiple simulated and real-world data sets, we highlight the significance of incorporating cell size coefficients into bulk data deconvolution methods. CSsingle uses absolute RNA content as a proxy for cell size and estimates cell sizes by leveraging ERCC spike-in controls, allowing researchers to empirically account for and eliminate systematic technical variations in RNA content quantification. When mixing cell types with significantly different cell sizes, incorporating cell size coefficients into the CSsingle method is crucial for achieving accurate bulk data deconvolution by normalizing RNA content biases. Using the influenza challenge data set as an example, we have demonstrated that CSsingle can accurately predict the proportions of neutrophils and lymphocytes, as well as the NLR from whole blood samples while incorporating cell size correction.

In this study, we also investigated the effect of cell size correction on deconvolution accuracy in ST data, aiming to determine whether accounting for RNA content differences improves cell type proportion estimates. Surprisingly, no significant differences in deconvolution accuracy were observed when cell size correction was used compared to uncorrected analyses. This result may be attributed to the dominant influence of incomplete cells captured by 2D slides and partial cell overlaps in ST data, which can obscure the impact of cell size differences. Additionally, the limited resolution of spatial platforms may render cell size adjustments inadequate at the spot level.

CSsingle also facilitates robust estimation of cell composition for mixtures and signature matrices derived from different sources. The iteratively reweighted least squares algorithm, integrated with robust initial estimates, efficiently addresses the technical and biological variations between individual bulk mixtures and the signature matrix. In both human pancreatic islet and PBMC data sets, we have demonstrated CSsingle’s versatility and robustness to accurately predict cell composition for both bulk and single-cell reference data obtained from various laboratory protocols and disease states, affirming the broad applicability of CSsingle across different single-cell sequencing methods.

In the context of spatial transcriptomics, CSsingle offers a powerful advantage by enabling the detection of enriched cell types at single-spot resolution rather than relying on pre-clustered regions, providing a high-resolution analysis of cellular composition within specific tissue locations. Prior knowledge of cell type enrichment guides spatial deconvolution by constraining plausible cell type combinations, preventing biologically implausible results and improving computational efficiency. Additionally, enriched markers amplify cell type-specific signals, mitigating ambiguities caused by low resolution or noise, which is especially critical for resolving sparse and noisy ST data. The enrichment at single-spot resolution captures subtle variations in cell type distribution that might be overlooked when analyzing broader tissue regions. By independently examining each spatial spot, CSsingle enhances the accuracy of deconvolution, facilitating detailed mapping of cellular interactions. For example, in human fetal pancreatic tissue, CSsingle provided valuable information on mesenchymal-acinar interactions at different developmental stages, demonstrating its ability to capture dynamic cellular interactions and developmental transitions. This fine-scale analysis demonstrates CSsingle’s potential to enhance our understanding of tissue architecture and cellular behavior, particularly in complex or heterogeneous microenvironments.

Esophageal cancer is the seventh leading cause of cancer-related mortality worldwide.^86^ Cancer immunotherapies have emerged as a promising neoadjuvant treatment strategy for advanced esophageal cancer.^87^ However, there are limited predictive biomarkers available to identify patients with gastroesophageal cancer who benefit the most from immunotherapy. In this study, we also performed a comprehensive and biologically significant validation of CSsingle using more than 700 normal and diseased samples of gastric esophageal tissue. In particular, we are the first to reveal the critical role of mosaic columnar cells (MCCs) in immunochemotherapy for patients with EAC. By analyzing large-scale bulk gene expression data from Barrett’s esophagus and esophageal adenocarcinoma, we reveal that MCCs are strikingly enriched in these conditions, while barely detectable in both normal esophageal tissue and esophageal squamous cell carcinoma (ESCC). CSsingle estimates reveal the predominant presence of MCC in esophageal intestinal metaplasia and adenocarcinoma, highlighting the specific association of MCC with the pathogenesis of EAC. Intriguingly, our findings suggest that MCCs may play a pivotal role in shaping tumor responses to ICI-based therapy in patients with EAC, positioning them as promising predictive biomarkers for therapeutic outcomes.

CSsingle addresses key challenges in the study of cellular heterogeneity by (i) providing accurate and robust decomposition of bulk and spatial transcriptomic data, (ii) applying cell size correction using ERCC spike-in controls to effectively correct for biases due to inherent differences in total RNA content across cell types, (iii) effectively handling technical and biological variations between individual mixtures and the signature matrix, and (iv) enhancing fine-scale analysis for spatial transcriptomic data.

## Method

### Construction of the signature matrices

Most current cell-type deconvolution techniques, which depend on a signature matrix composed of cell-type-specific GEPs, operate under the assumption that cells can be categorized into a predetermined set of types and the prevalent cell types within the bulk tissue are adequately reflected in the scRNA-seq data. To build a signature matrix from scRNA-seq data, we started with a read or UMI count matrix. We first summarized the gene counts of all cells assigned to the same cell type. The summation was followed by normalization based on total count and multiplication by a scale factor of 10^4^. This process produced a matrix of genes × cell types (denoted as 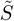), from which a submatrix (denoted as *S*) was derived by selecting a set of differentially expressed genes. The identification of differentially expressed genes was accomplished using the FindAllMarkers function with the default parameters of the Seurat R package (version 5.0.1). Specifically, we initially isolated differentially expressed genes using a log-scale threshold of ≥ 0.25-fold overexpression in a given cell population relative to all others. Subsequently, non-significant genes with a p-value larger than 0.01 (Wilcoxon Rank Sum test or likelihood-ratio test) were filtered out. Next, we ranked differentially expressed genes in ascending order by their p-values and selected the top *N* marker genes for each cell type as the most differentially expressed marker genes. Additionally, we excluded marker genes shared between two or more cell types. Finally, we generated multiple signature matrices by varying *N* from 50 to 200 with step 50 by default.

### Selecting the optimal signature matrix

To determine the optimal signature matrix, CSsingle is integrated with each signature matrix to estimate the cell type proportions. The optimal signature matrix *S*^*^ is selected as the candidate matrix whose inferred bulk/ST gene expression matrix has the highest Spearman correlation coefficient with the real bulk/ST gene expression matrix.

### CSsingle model

CSsingle is an iteratively reweighted linear regression model that decomposes the bulk or ST gene expression data into a set of predefined reference cell types to estimate cell abundances. In order to accurately estimate the cell type fraction, we designed a novel weighting scheme to properly adjust the contribution of each marker gene. We denote 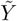 as a cell type mixture. Both 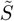 and 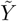 consist of column-normalized expression values, so that each cell type and cell type mixture have the same total count of 10^4^ across all genes shared between the scRNA-seq and cell type mixture. Additionally, we denote the truncated version of 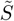 and 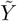, containing only *N* significant differential expressed marker genes, as *S* and *Y*, respectively. Let 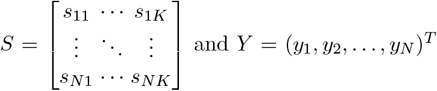 and *Y* = (*y*_1_, *y*_2_, …, *y*_*n*_)^*T*^. The deconvolution model for an observed cell type mixture within CSsingle is defined as follows:

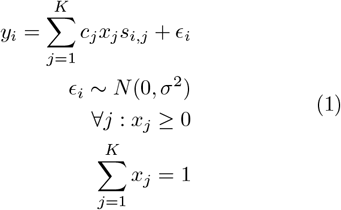

where *S* is an *N* × *K* signature matrix that contains GEPs for the *N* marker genes across *K* cell types, *Y* is an *N* × 1 vector representing a single bulk or spatial spot GEP for the same *N* marker genes, *X* = (*x*_1_, *x*_2_, …, *x*_*K*_)^*T*^ is a *K* × 1 vector containing the cell type composition from *Y, c*_*j*_ denotes the cell size coefficient of cell type *j*, and *ϵ* models measurement noise and other possible un-modeled factors. The constraints in (Eq.1) require the cell type fractions to be positive and to sum up to one.

Within CSsingle, we introduce the cell size factor *c*_*j*_ to account for differences in RNA content across cell types and their effect on the deconvolution problem. Cell size coefficients are estimated by using ERCC spike-in controls which allow absolute RNA expression quantification. Specifically, gene counts were nomalized by dividing by the upper-quartile (UQ; 75th percentile by default) of the ERCC spike-in counts to adjust for varying sequencing depths and other potential technical effects. Next, we calculated the absolute cellular RNA content by summing the normalized gene counts for each cell. We define the cell size coefficient *c*_*j*_ as the mean value of absolute cellular RNA contents in each cell of cell type *j*.

To minimize the technical and biological variation between individual cell type mixtures and the signature matrix, CSsingle introduces *w*_*i*_ by up-weighting marker genes with strong concordance and down-weighting genes with weak concordance between individual cell type mixtures and the signature matrix. To estimate *X* and 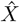, CSsingle minimizes the weighted squared error as follows:

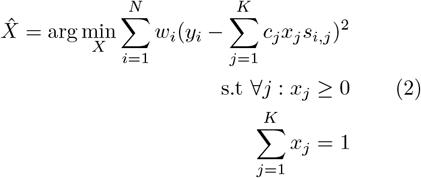

Here, the weights *w*_*i*_ are interdependent with both the residuals and the estimated coefficients, forming an iterative dependency loop. To address this, we employed an iterative approach known as iteratively reweighted least squares (IRIS) to solve the weighted least squares problem.

To achieve an efficient and robust estimation, it is vital to give careful consideration to the initial estimates. CSsingle takes advantage of the sectional linear relationship of the marker genes to generate an efficient set of initial estimates. We define ℳ^*j*^ as a finite set of marker genes for cell type *j*, where *j* = 1, 2, …, *K*. Let *N*_*j*_ = |ℳ^*j*^| represent the total number of marker genes for cell type *j*, and consequently, *N* = ∑_*j*_ *N*_*j*_. For each cell type *j*, CSsingle employs a linear regression model to fit the cell type mixture *Y* and each cell-type-specific GEPs within signature matrix *S* with its marker genes in log-scale using Eq.3. The goal is to find the best-fitting curve with a regression slope of one to a data set comprising *N*_*j*_ observations of cell-type-specific gene expression values 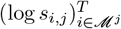, together with corresponding observations of the bulk or ST gene expression values of 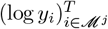 by minimizing the sum of the squares of the offsets of the points from the curve.

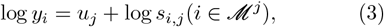

where *u*_*j*_ = log(*c*_*j*_*x*_*j*_) is the disturbance term for cell type *j* estimated from the above linear regression with a slope of one. We then define 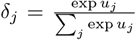, representing the estimated proportion of RNA content derived from cell type *j* in *Y*, where exp *u*_*j*_ = *c*_*j*_*x*_*j*_ represents the estimated total RNA content derived from cell type *j* in *Y*. Next, we use *δ*_*j*_ to generate the estimated cell-type mixture as *Y* ^*^ = (*t*_1_, *t*_2_, …, *t*_*N*_)^*T*^, where 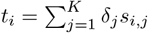. Finally, the initial weight for gene *i* is defined as:

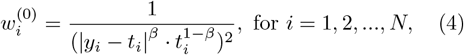

Let:

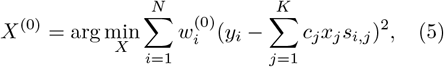

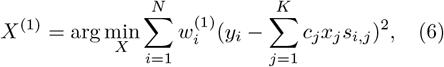

where 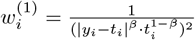 and 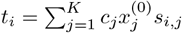,

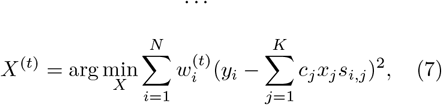

where 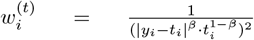 and 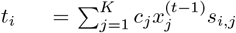.

The estimated coefficients converge when ∥*X*^(*t*)^ − *X*^(*t*−1)^∥ ≤ 0.01, and the optimal *X*^(*t*)^ is the final cell type composition estimated from the bulk data. We introduce *β* to balance the relative contributions of the difference between *y*_*i*_ and *t*_*i*_ and the magnitude of *t*_*i*_, with the goal of enhancing cross-platform performance and mitigating the influence of highly expressed genes in the least squares fitting procedure. The optimal *β*^*^ was selected from {0, 0.5, 1} to maximize the Spearman correlation between the inferred and real bulk/ST GEPs.

### Enhancing performance in microarray data and high technical variation scenarios

Given the major differences between RNA sequencing and microarray techniques, deconvolution might prove ineffective in scenarios where excessive technical variation exists. For microarray, background hybridization and probe saturation can impede the detection of transcripts at both low levels and high levels. In contrast, RNA sequencing enables the detection of a broader range of transcripts, including those with low abundance and high abundance.^88, 89^ We, therefore developed a strategy for handling bulk mixtures derived from microarrays, tailored for signature matrix generated from RNA sequencing, or vice versa. We introduced an upper bound *q*, which limits the maximum value that any weight can take on. The adjusted weights are defined as:

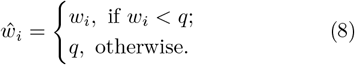

The upper bound *q* is selected as follows. The possible values for *q* are defined as the *τ* ^*th*^ quantile of the gene weights (*w*_1_, *w*_2_, · · ·, *w*_*N*_)^*T*^, where *τ* is selected from 0.01 to 1 with a step of 0.01. For each possible value of *q*, we obtain an estimate for *X*, denoted as 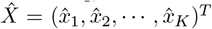, by minimizing the weighted squared error with respect to the adjusted weights 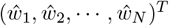:

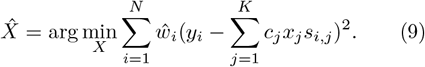

Subsequently, the inferred bulk/ST GEP 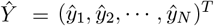 is defined as follows:

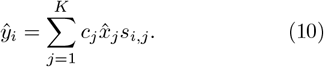

CSsingle calculates the Spearman correlation between the inferred and real bulk/ST GEPs, *Ŷ* and *Y*. The value of *q* corresponding to the maximum correlation coefficient is selected.

### Enrichment analysis for identifying cell types within a single spatial spot

Inspired by spatialDWLS,^27^ which uses PAGE^90^ to identify enriched cell types in clustered regions, we applied a modified approach that integrates multiple Z-score criteria to detect enriched cell types at the level of individual spatial spots. For each gene, the fold change is determined by the ratio of its expression at each spatial spot relative to the average expression across all spots. To establish a reference, the mean and standard deviation of fold changes across all genes are denoted as *µ* and *σ*, respectively. For each spatial spot, the mean fold change of marker genes for cell type *j* is calculated and denoted as *f*_*j*_. The *Z*-score for cell type *j* is then defined as follows:

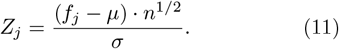

We identified enriched cell types per spatial spot using three *Z*-score-based strategies:

1. Selecting cell types with *Z*-scores *>* 0;
2. Identifying the cell type with the highest *Z*-score;
3. Normalizing *Z*-scores across spots (mean = 0, SD = 1) to ensure comparability by mitigating differences in *Z*-score distributions, and selecting cell types with normalized *Z*-scores *>* 0.

The results of all three methods were combined to determine the final set of enriched cell types per spatial spot.

### Construction of artificial bulk data sets

The aggregated counts for the simulated bulk data set are derived from a scRNA-seq data set, with the bulk counts computed as the sum of gene counts across all cells within the same sample. Specifically, for the scRNA-seq data set spanning fewer than five samples, we generate *t* bootstrap replicates, each of which matches the size of the original scRNA-seq data set by randomly sampling cells with replacement. Next, counts for the artificial bulk data set are generated from each bootstrap replicate by summing up gene counts from all cells within the same sample. For the artificial bulk data set in Fig.3b, we used *t* = 100. The actual cellular proportion of cell type *j* in the sample *s* is calculated by

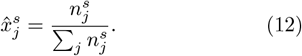

where 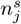 is the number of cells for cell type *j* in sample *s*.

### Functional enrichment analysis

The identification of marker genes for mosaic columnar cells (MCCs) was accomplished using the FindAllMarkers function of the Seurat R package. Specifically, we initially isolated differentially expressed genes using a log-scale threshold of ≥ 0.25-fold over-expression in a given cell population relative to all others. Subsequently, we ranked the differentially expressed genes in ascending order by their p-values and selected the top 25 marker genes for MCC. The R package clusterProfiler was used to identify and visualize enriched GO terms, highlighting significant terms in biological processes.

### Gene set variation analysis

Gene Set Variation Analysis (GSVA), implemented via the GSVA R package, was used to calculate enrichment scores for functional terms from the Hallmark and Reactome collections. Pathway gene sets were sourced from the MSigDB database (Reactome: c2.cp.reactome.v2024.1.Hs.symbols.gmt, Hallmark: h.all.v2024.1.Hs.symbols.gmt). Single-sample gene set enrichment analysis (ssGSEA) was employed using the GSVA R package to assess the activation level of the “TGF-*β* receptor signaling in the EMT” pathway, based on normalized enrichment scores. Additionally, the ssGSEA method was utilized to evaluate pathway activity in mesenchymal-acinar interacting cells within pancreas ST data.

### Statistical analysis

All statistical analyses were conducted using R (version 4.4.0; available at https://cran.r-project.org/). Specific statistical tests are detailed in the figures and their respective captions when applicable. A *p*-value threshold of 0.05 was used to determine statistical significance for all tests, unless otherwise specified. Statistical significance between two groups of samples was assessed using a two-sided Wilcoxon test and reported as follows: ^*ns*^*p* ≥ 0.05, ^*^*p <* 0.05,^**^ *p <* 0.01,^***^ *p <* 0.001, and ^****^*p <* 0.0001. The cumulative survival time was estimated via the Kaplan–Meier method, with the log-rank test from the R survminer package employed to assess survival curve disparities.

### Systematic evaluation of CSsingle and comparison against baseline methods

In this study, we benchmarked the performance of CSsingle against 11 existing methods for bulk data deconvolution: DWLS, BayesPrism, CIBERSORT, CIBER-SORTx, MuSiC, MuSiC2, NNLS, SCDC, BisqueRNA, CAMmarker, and EPIC. Further details regarding their implementation and specific parameters can be found in the respective original publications and GitHub repositories. All parameters were initialized to their default values, unless stated differently.

- **DWLS**. We downloaded DWLS from https://bitbucket.org/yuanlab/dwls/src/default/. The signature matrix was constructed using the hurdle model in the MAST R package. In the event of negative values, the estimated proportions were adjusted to zero. When the function solve.QP fails to find a solution due to inconsistent constraints, DWLS outputs errors.
- **BayesPrism**. The R package BayesPrism was downloaded from https://github.com/Danko-Lab/BayesPrism.git.
- **CIBERSORT**. CIBERSORT was run online (https://cibersortx.stanford.edu/runcibersortx.php). No batch correction was applied.
- **CIBERSORTx**. CIBERSORTx was executed online (https://cibersortx.stanford.edu/runcibersortx.php). Batch correction was applied to reduce cross-platform variance.
- **MuSiC**. The R package MuSiC was downloaded from https://github.com/xuranw/MuSiC. MuSiC was employed with parameter ‘cell_size’ set as NULL (default value, estimating cell size coefficients from data), while MuSiC* was employed with ‘cell_size’ estimated using ERCC spike-in controls.
- **MuSiC2**. The MuSiC2 functions are available within the R package MuSiC. MuSiC2 was designed for the deconvolution of multi-condition bulk data, we thus ran it only for multi-condition bulk data. MuSiC2 was employed with parameter ‘cell_size’ set as NULL (default value, estimating cell size coefficients from data), while MuSiC2* was employed with ‘cell_size’ estimated using ERCC spike-in controls.
- **NNLS**. The NNLS method is implemented in the R package MuSiC.
- **SCDC**. The R package SCDC was downloaded from http://meichendong.github.io/SCDC. During the quality control procedure, we set the parameter ‘qcthreshold = 0.7’.
- **BisqueRNA**. The R package BisqueRNA was downloaded from https://github.com/cozygene/bisque.
- **CAMmarker**. The R package debCAM was downloaded from https://bioconductor.org/packages/release/bioc/html/debCAM.html. The parameter ‘MGlist’ defines a list of vectors, each containing known markers for one cell type. These markers are chosen from the candidate signature matrix with the lowest condition number.
- **EPIC**. The R package EPIC was downloaded from https://github.com/Gfeller-Lab/EPIC. The parameter ‘sigGenes’ defines a character vector consisting of gene names chosen to serve as a signature for the deconvolution process. These genes are selected from the candidate signature matrix with the lowest condition number.

We further benchmarked the performance of CSsingle against 7 existing methods for spatial transcriptomics data deconvolution: Seurat, RCTD, SpatialDWLS, Redeconve, SpatialDecon, SPOTlight, and Spa-tialDDLS. Further details regarding their implementation and specific parameters can be found in the respective original publications and GitHub repositories. All parameters were initialized to their default values, unless stated differently.

- **Seurat**. Deconvolution was performed using the FindTransferAnchors and TransferData functions from the Seurat (https://satijalab.org/seurat/) package.
- **RCTD**. The R package spacexr was downloaded from https://github.com/dmcable/spacexr. We set the parameter ‘doublet_mode’ to ‘full’ in the run.RCTD function.
- **SpatialDWLS**. Deconvolution was performed using the runDWLSDeconv functions from the Giotto (https://rubd.github.io/Giotto_site/) package. We used the signature matrix with the lowest condition number.
- **Redeconve**. The R package Redeconve was downloaded from https://github.com/ZxZhou4150/Redeconve.
- **SpatialDecon**. The R package BisqueRNA was downloaded from https://github.com/Nanostring-Biostats/SpatialDecon/.
- **SPOTlight**. The R package SPOTlight was downloaded from https://github.com/MarcElosua/SPOTlight.
- **SpatialDDLS**. The SpatialDDLS method is implemented using the R package SpatialDDLS (https://github.com/diegommcc/SpatialDDLS).

### Assessment of deconvolution performance

We assessed the performance of various deconvolution methods using Pearson’s correlation coefficient (R), root mean squared deviation (RMSD), and mean absolute deviance (mAD) as evaluation metrics. These metrics are calculated using the following equations:

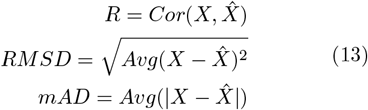

where *X* and 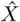 are actual and estimated cell type proportions, respectively.

## Data availability

All data analyzed in this study are publicly available through online sources. Accession numbers and reference links to all data sources are presented in Table 1.

## Code availability

The CSsingle method is implemented in an R package called ‘CSsingle’. Both the source code and instructions are available at https://github.com/wenjshen/CSsingle.

## Acknowledgements

This work was supported by Guangdong Basic and Applied Basic Research Foundation (2023A1515030154) to W.S., and the NIGMS Maximizing Investigators’ Research Award (MIRA) R35 GM146960 to X.M.Z..

