## Appendix for "CSsingle: A Unified Tool for Robust Decomposition of Bulk and Spatial Transcriptomic Data Across Diverse Single-Cell References"

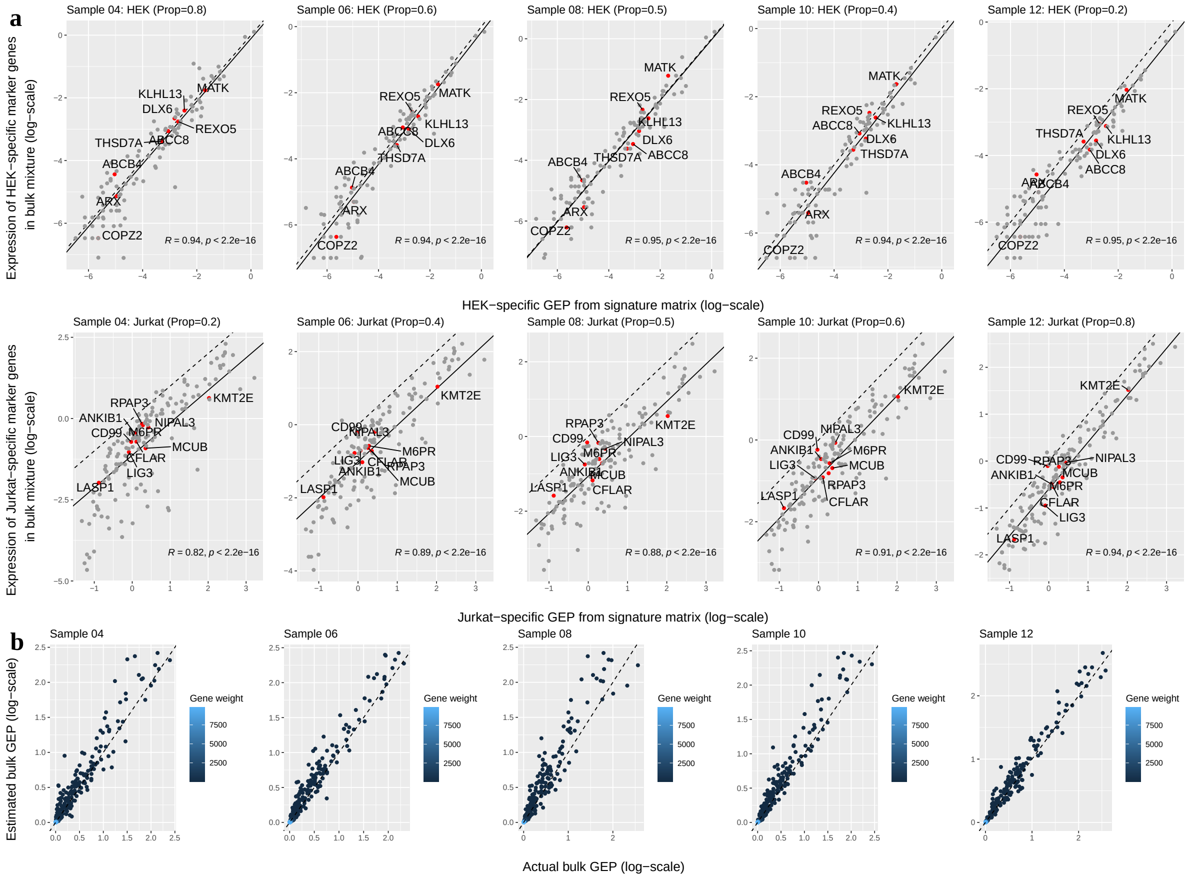


Figure S1: **Sectional linear relationship between individual bulk mixtures of HEK and Jurkat cells and the signature matrix. a** Linear regression with a slope of one for individual bulk mixtures and the cell type-specific GEPs for samples 04, 06, 08, 10, and 12. Top row: Linear regression between mean expression levels of HEK-specific marker genes in HEK-specific GEPs and their expression in each bulk mixture. Bottom row: Linear regression between mean expression levels of Jurkat-specific marker genes in Jurkat-specific GEPs and their expression in each bulk mixture. The dashed line in each plot represents the line of $y = x$. The signature matrix was constructed by selecting the top 150 marker genes for each cell type. Gene symbols of the top 10 most significant marker genes were plotted. **b** Scatter plots comparing estimated and actual bulk GEPs, colored by gene weights. The dashed line in each plot represents the line of $y = x$.

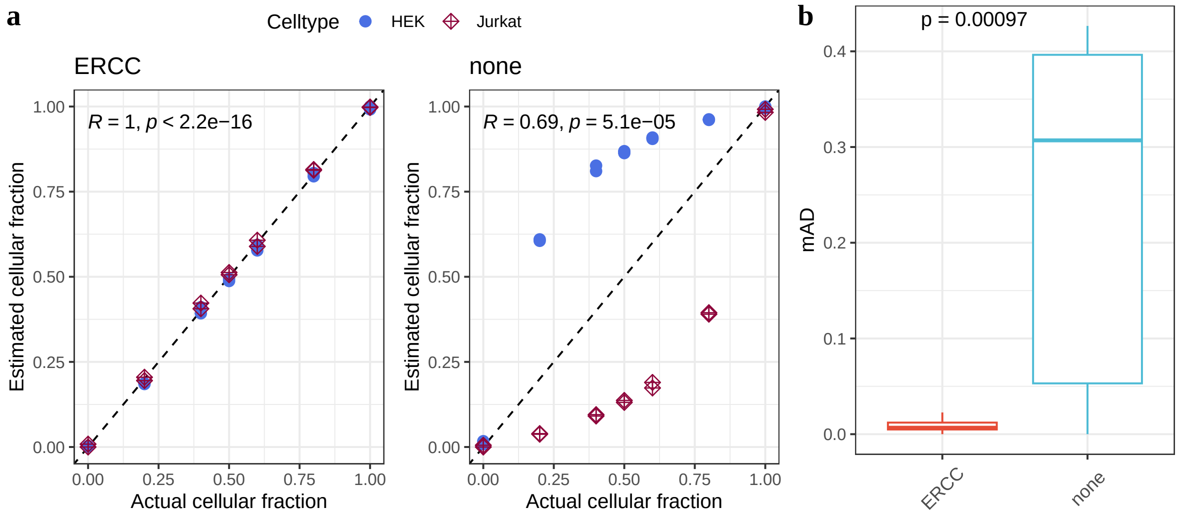


Figure S2: **CSsingle deconvolution performance in HEK and Jurkat cell mixtures: with versus without cell size correction. a** Plots show the Pearson correlation between estimated and actual cell type proportions with (left) and without (right) cell size correction. **b** Comparison of deconvolution performance with versus without cell size correction in terms of mAD.

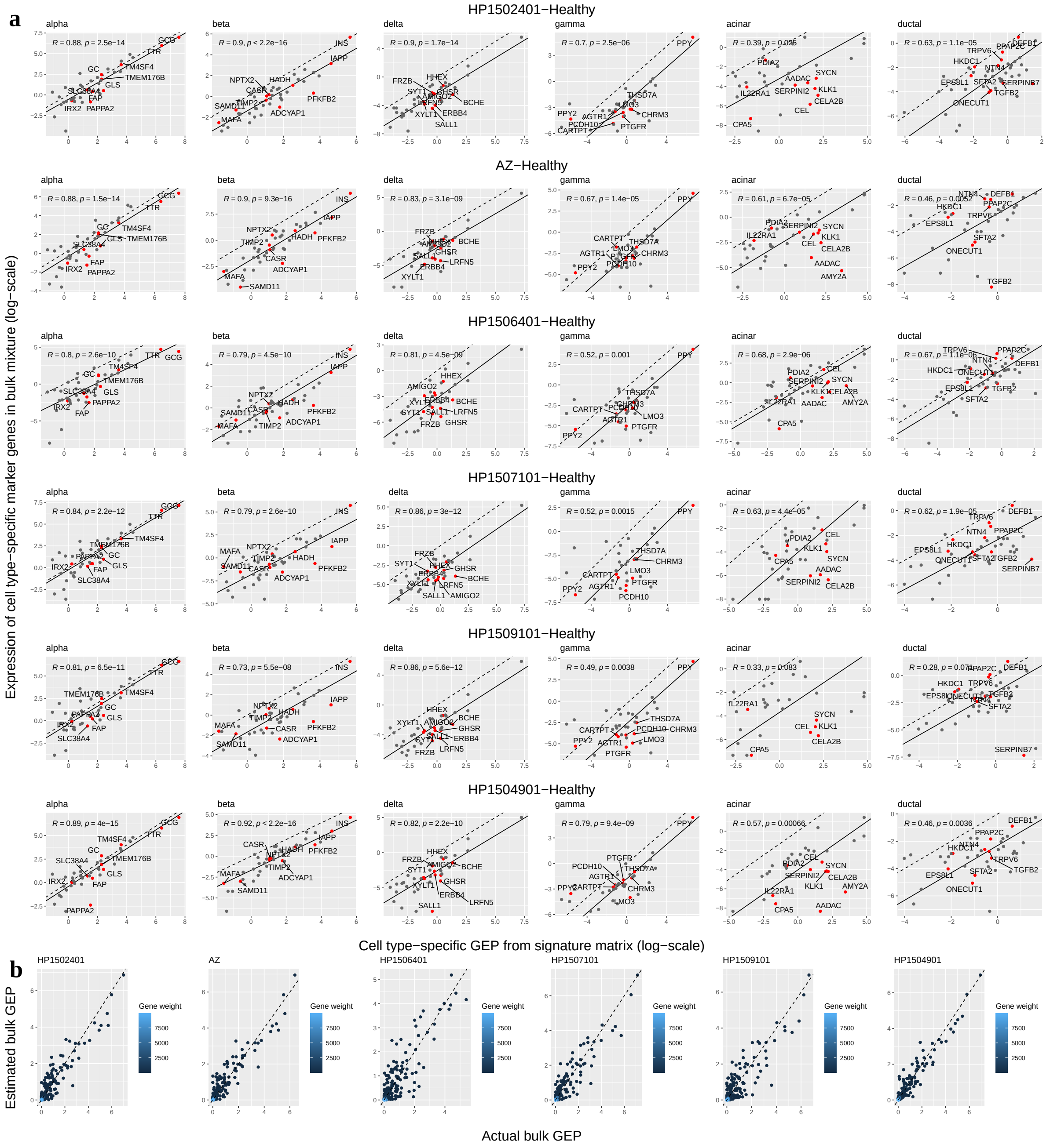


Figure S3: **Sectional linear relationship between individual bulk mixtures and the signature matrix for healthy samples in the human pancreatic islet data set. a** Linear regression with a slope of one for individual bulk mixtures and the cell type-specific GEPs for healthy pancreatic islet samples. The dashed line in each plot represents the line of $y = x$. The signature matrix was constructed by selecting the top 150 marker genes for each cell type. Gene symbols of the top 10 most significant marker genes were plotted. **b** Scatter plots comparing estimated and actual bulk GEPs, colored by gene weights. The dashed line in each plot represents the line of $y = x$.

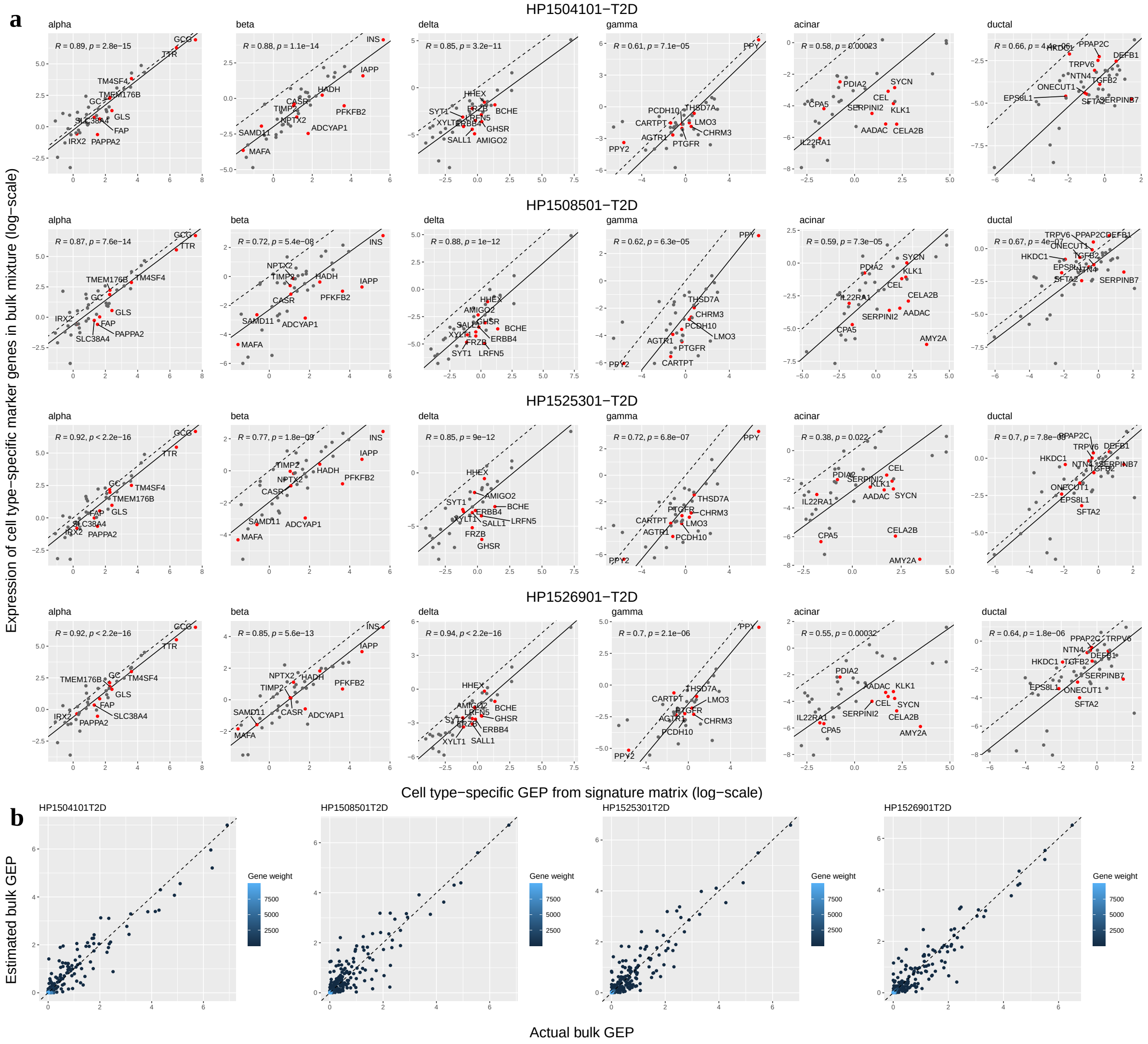


Figure S4: **Sectional linear relationship between individual bulk mixtures and the signature matrix for T2D samples in the human pancreatic islet data set. a** Linear regression with a slope of one for individual bulk mixtures and the cell type-specific GEPs for T2D pancreatic islet samples. The dashed line in each plot represents the line of $y = x$. The signature matrix was constructed by selecting the top 150 marker genes for each cell type. Gene symbols of the top 10 most significant marker genes were plotted. **b** Scatter plots comparing estimated and actual bulk GEPs, colored by gene weights. The dashed line in each plot represents the line of $y = x$.

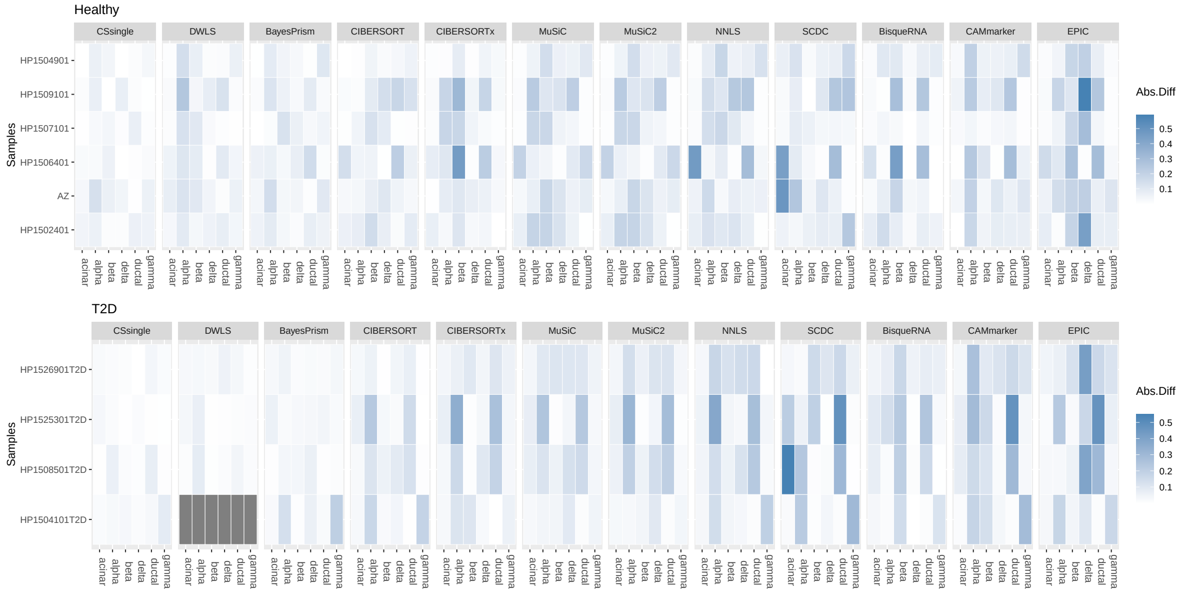


Figure S5: **Decomposition benchmark in human pancreatic islet tissue.** Benchmarking results are shown in heatmaps, displaying the mean absolute deviation (mAD) between true and estimated cell type proportions for healthy (top panel) and T2D (bottom panel) samples. Darker colors represent higher mAD values.

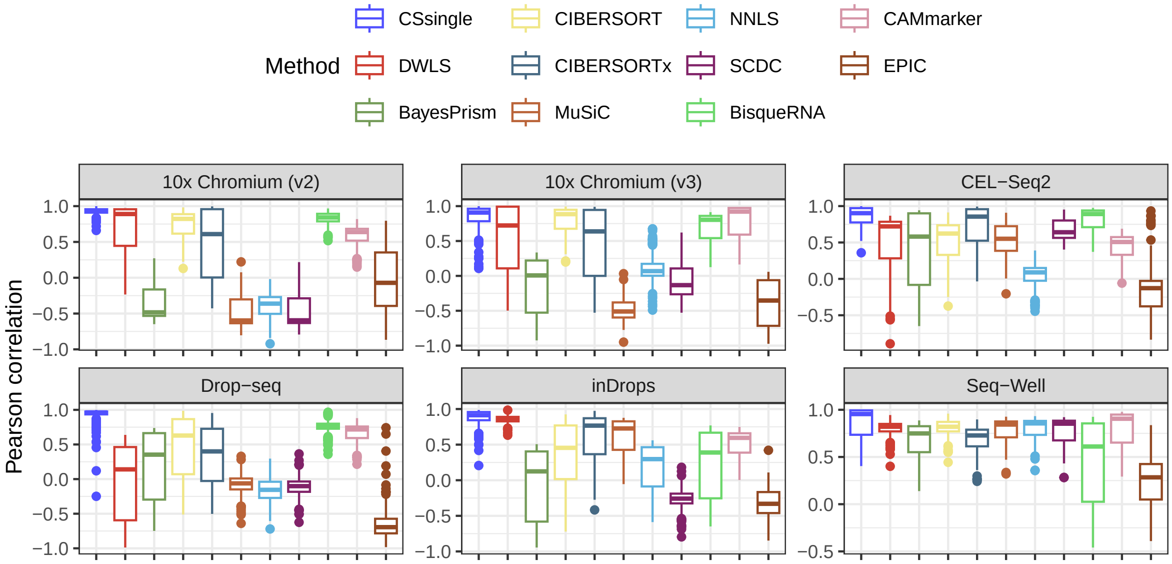


Figure S6: **Decomposition benchmark in human PBMC.** Benchmarking of deconvolution accuracy in terms of Pearson correlation. Data are depicted using boxplots, where the center line indicates the median, the box boundaries signify the upper and lower quartiles, and the whiskers span the full range of maximum and minimum values.

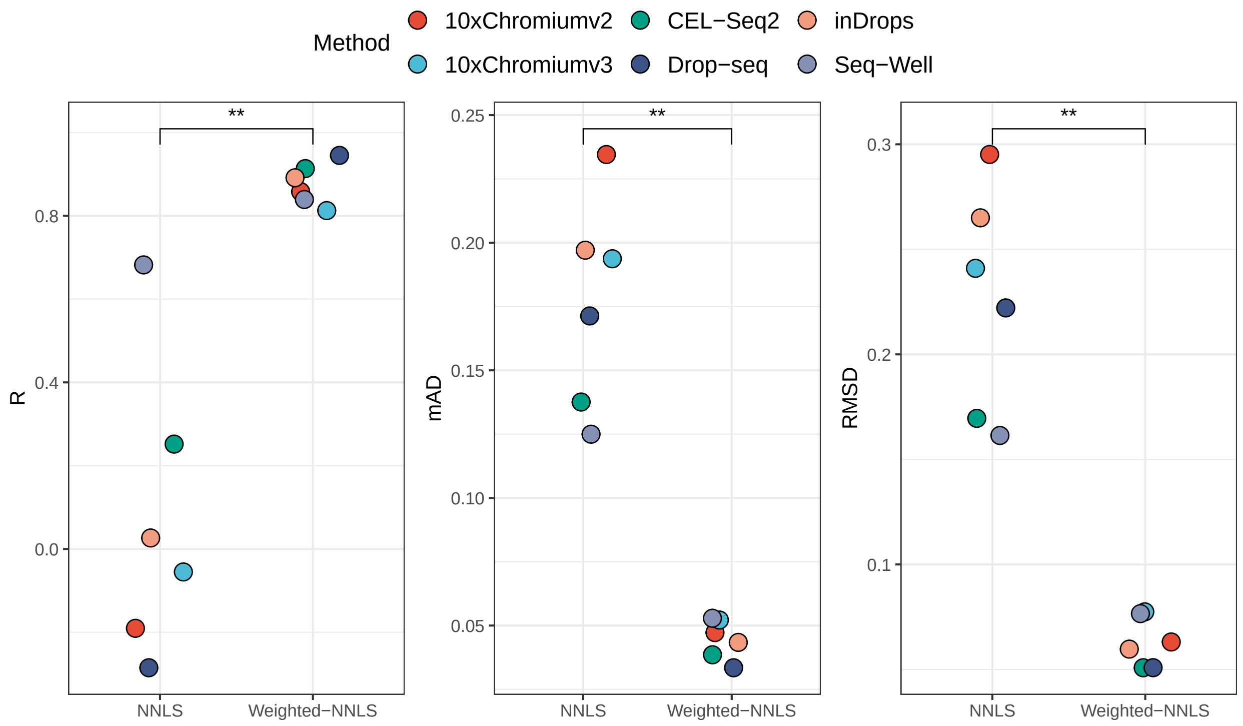


Figure S7: **Comparison of the performance of CSsingle initialized with two different weighting strategies.** Left panel: constant gene weights (NNLS) and Right panel: more weights on strong concordant genes and less weights on weak concordant genes (weighted-NNLS). Reported ‘R’ corresponds to Pearson correlation and $p$-values indicate the significance of these correlations.

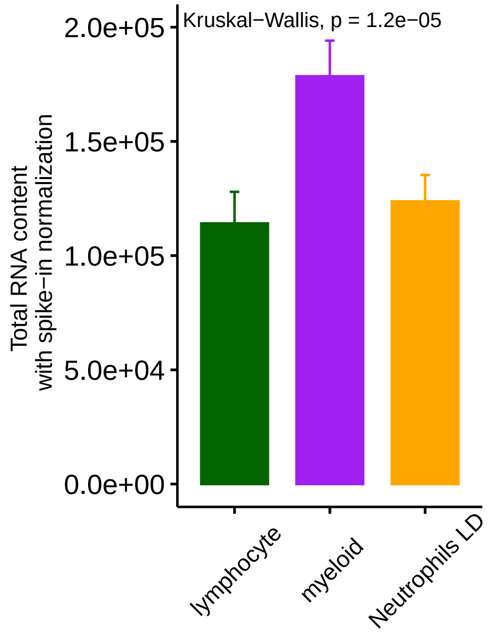


Figure S8: **Comparison of the estimated cell sizes of three large immune cell groups.** Statistical significance was assessed using the Kruskal-Wallis test.

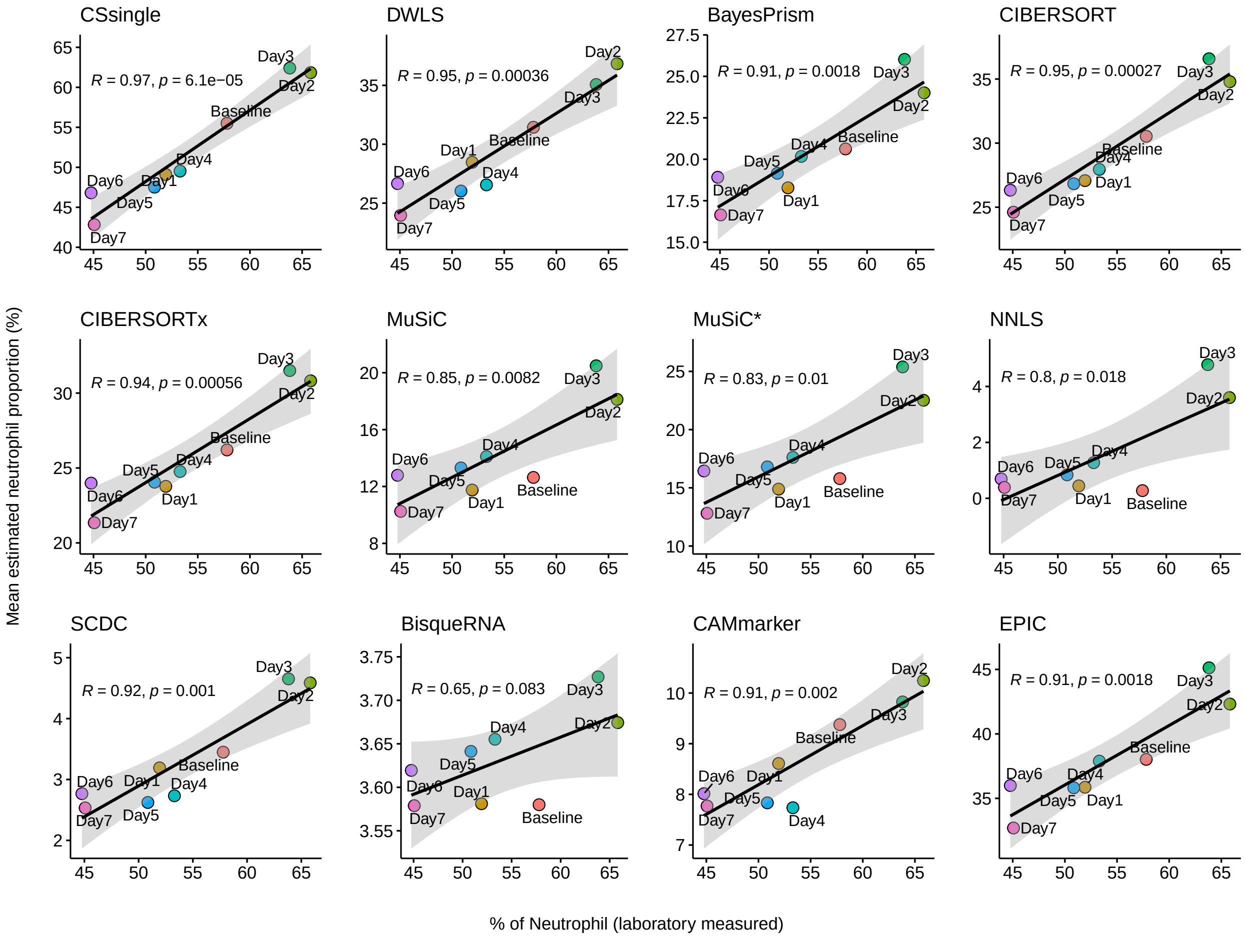


Figure S9: **Correlation between laboratory measured and estimated proportions of neutrophil in SI group of influenza H3N2.** Data points are labeled by days post-inoculation, with the baseline denoting pre-inoculation and day 1 marking the day of inoculation. Reported 'R' corresponds to Pearson correlation and $p$-values indicate the significance of these correlations.

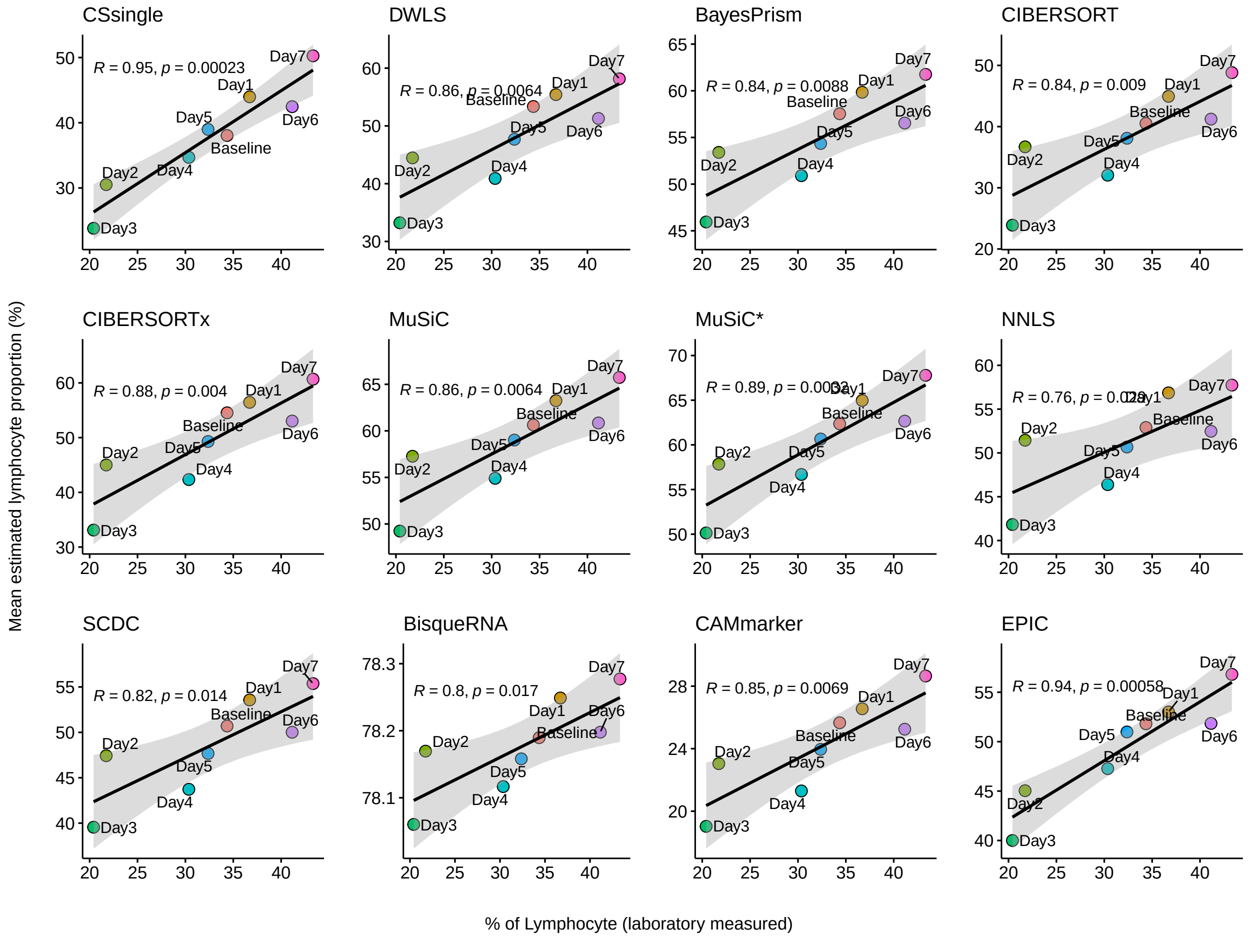


Figure S10: **Correlation between laboratory measured and estimated proportions of lymphocyte in SI group of influenza H3N2.** Data points are labeled by days post-inoculation, with the baseline denoting pre-inoculation and day 1 marking the day of inoculation. Reported ‘R’ corresponds to Pearson correlation and $p$-values indicate the significance of these correlations.

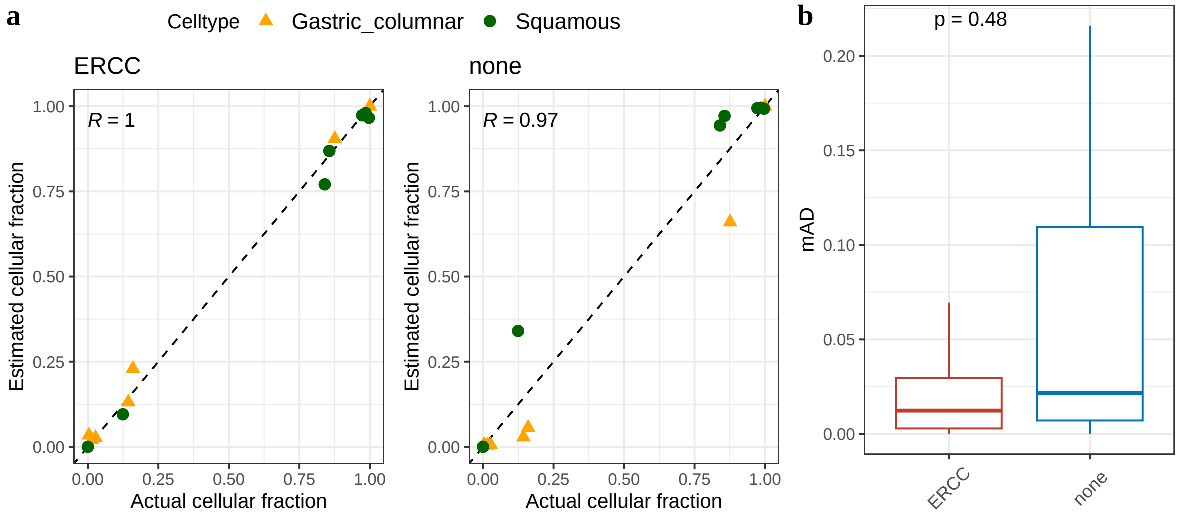


Figure S11: **CSsingle deconvolution performance in human normal squamous-columnar junction (N-SCJ) tissue: with versus without cell size correction. a** Plots show the Pearson correlation between estimated and actual cell type proportions with (left) and without (right) cell size correction. **b** Comparison of deconvolution performance with versus without cell size correction in terms of mAD.

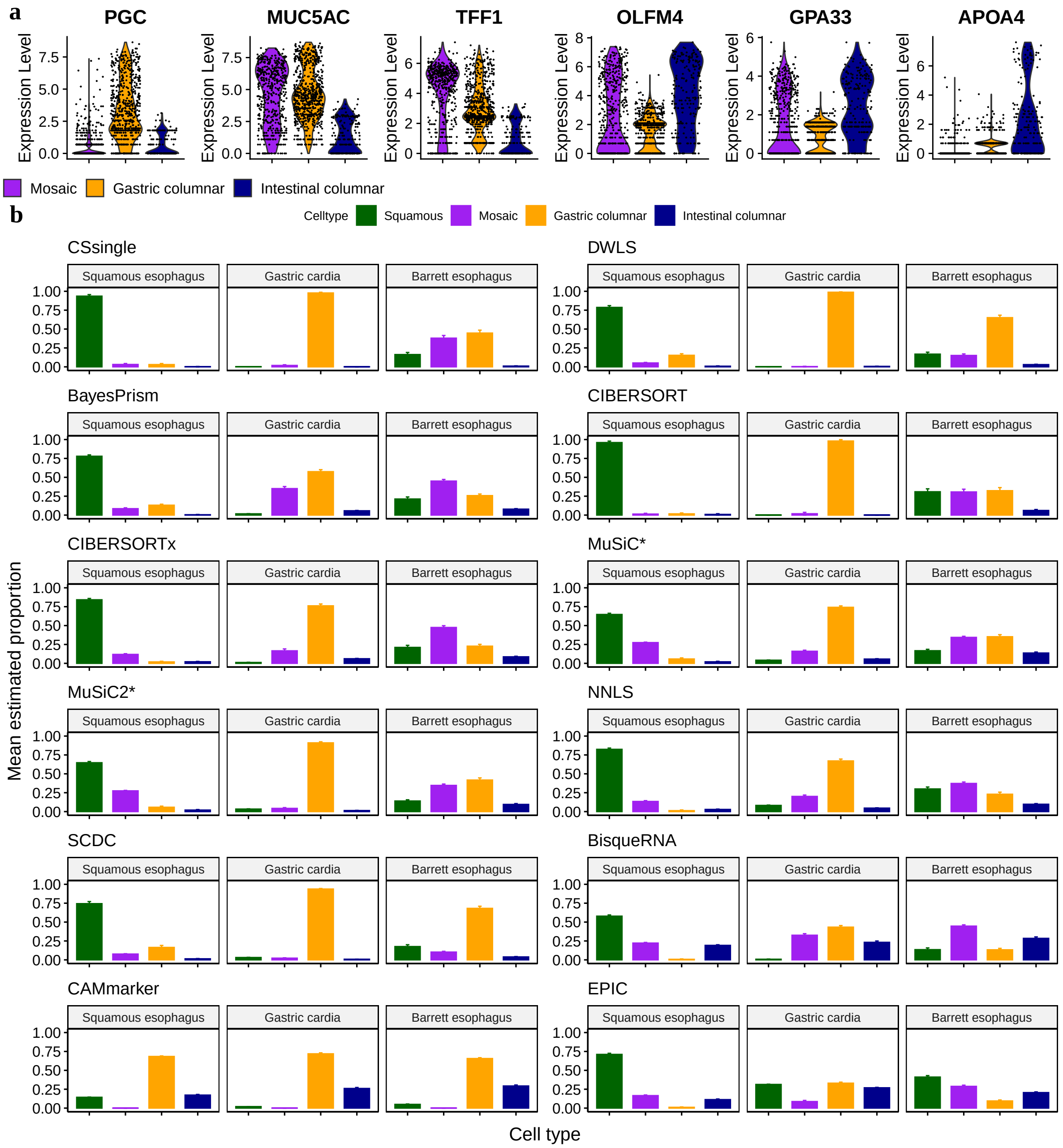


Figure S12: **Decomposition benchmark in human Barrett's esophagus tissue. a** Violin plots of marker gene expression in mosaic, gastric, and intestinal columnar cells. **b** Comparison of decomposition estimates between CSsingle and other methods for 233 epithelium samples derived from NE, NGC and BE. Each color represents a cell cluster. Error bars represent standard deviation.

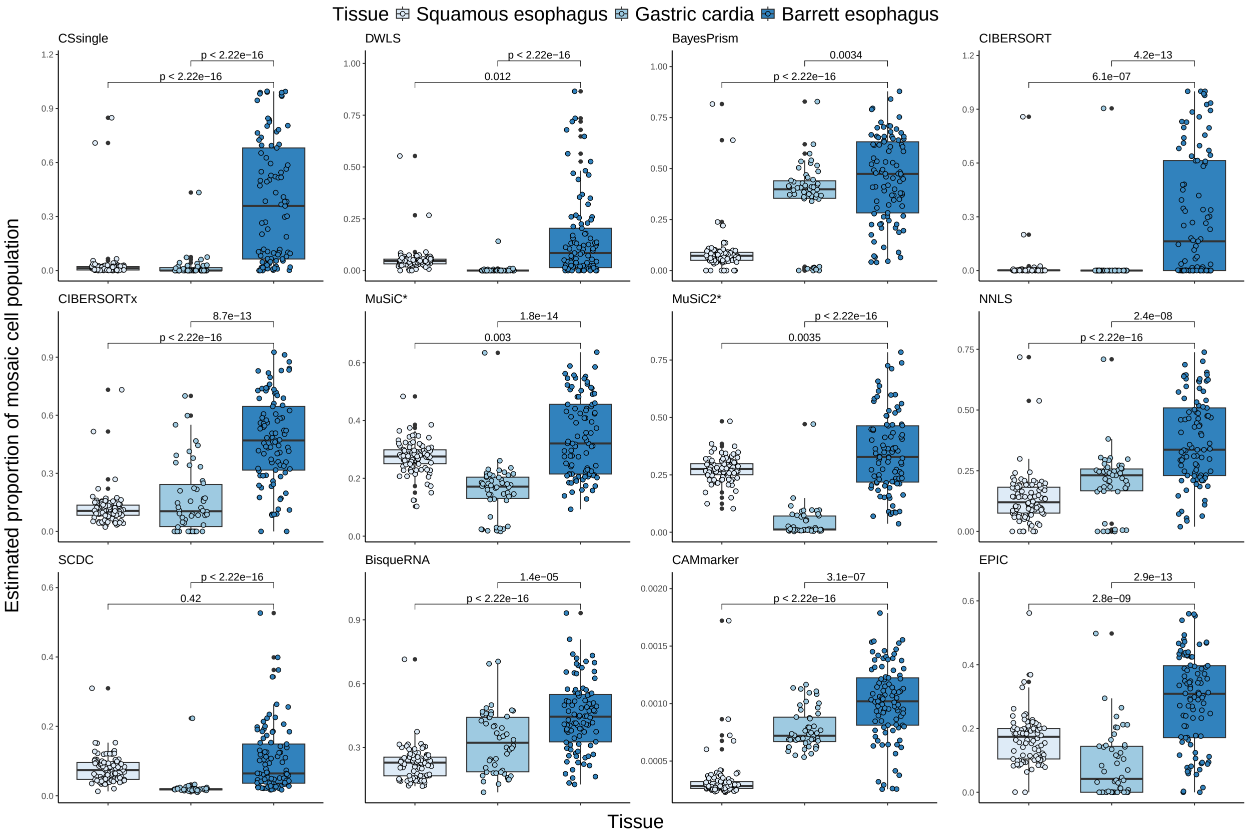


Figure S13: **Comparison of the estimated proportions of mosaic columnar cell (MCC) in NE, NGC, and BE.** The $p$-values indicate the significance of the one-sided Wilcoxon-test. A $p$-value less than 0.05 is considered statistically significant, indicating MCC is significantly increased in proportion in BE compared with the proportions in NE or NGC.

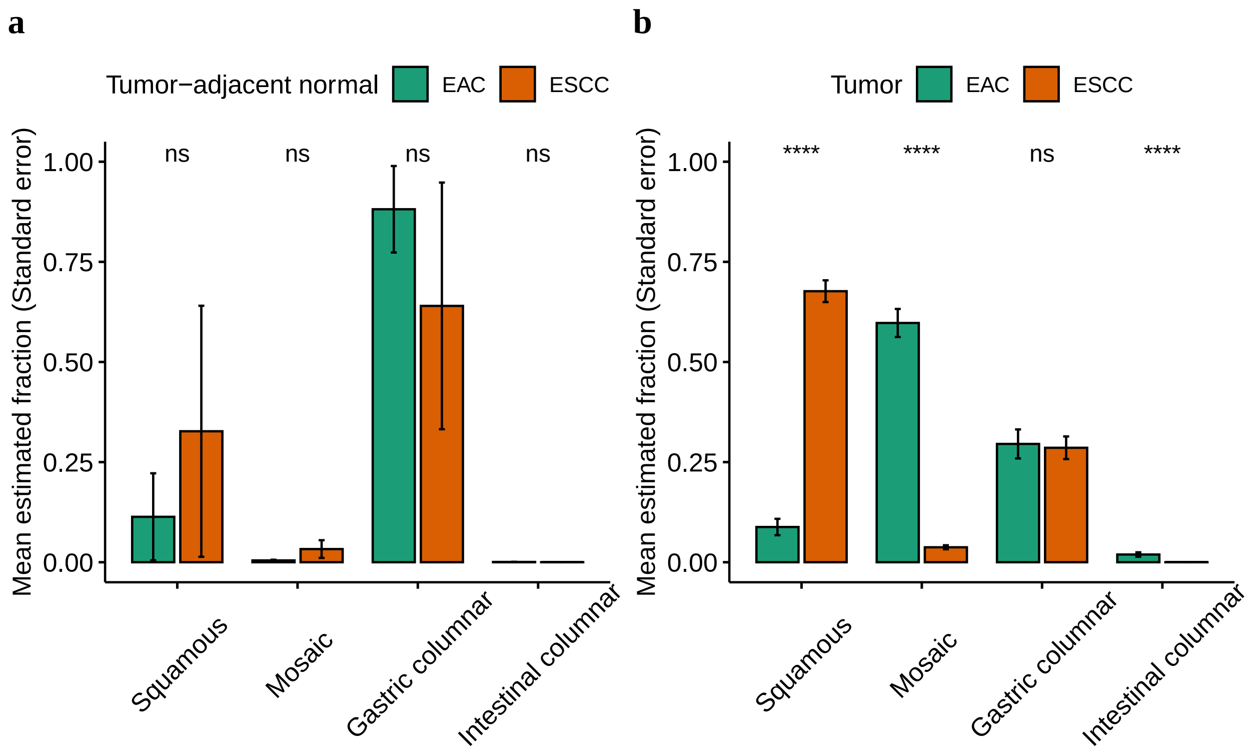


Figure S14: **Application of CSsingle to deconvolve oesophageal carcinoma biopsies.** The relative abundance of four epithelial cell populations in tumor-adjacent normal (**a**) and tumor (**b**) samples in TCGA-ESAD cohort. Error bars represent standard deviation. Statistical significance between ESCC and EAC samples was assessed using a two-sided Wilcoxon test and indicated as follows: $^{ns}p \geq 0.05$, $^{*}p < 0.05$, $^{**}p < 0.01$, $^{***}p < 0.001$, and $^{****}p < 0.0001$.

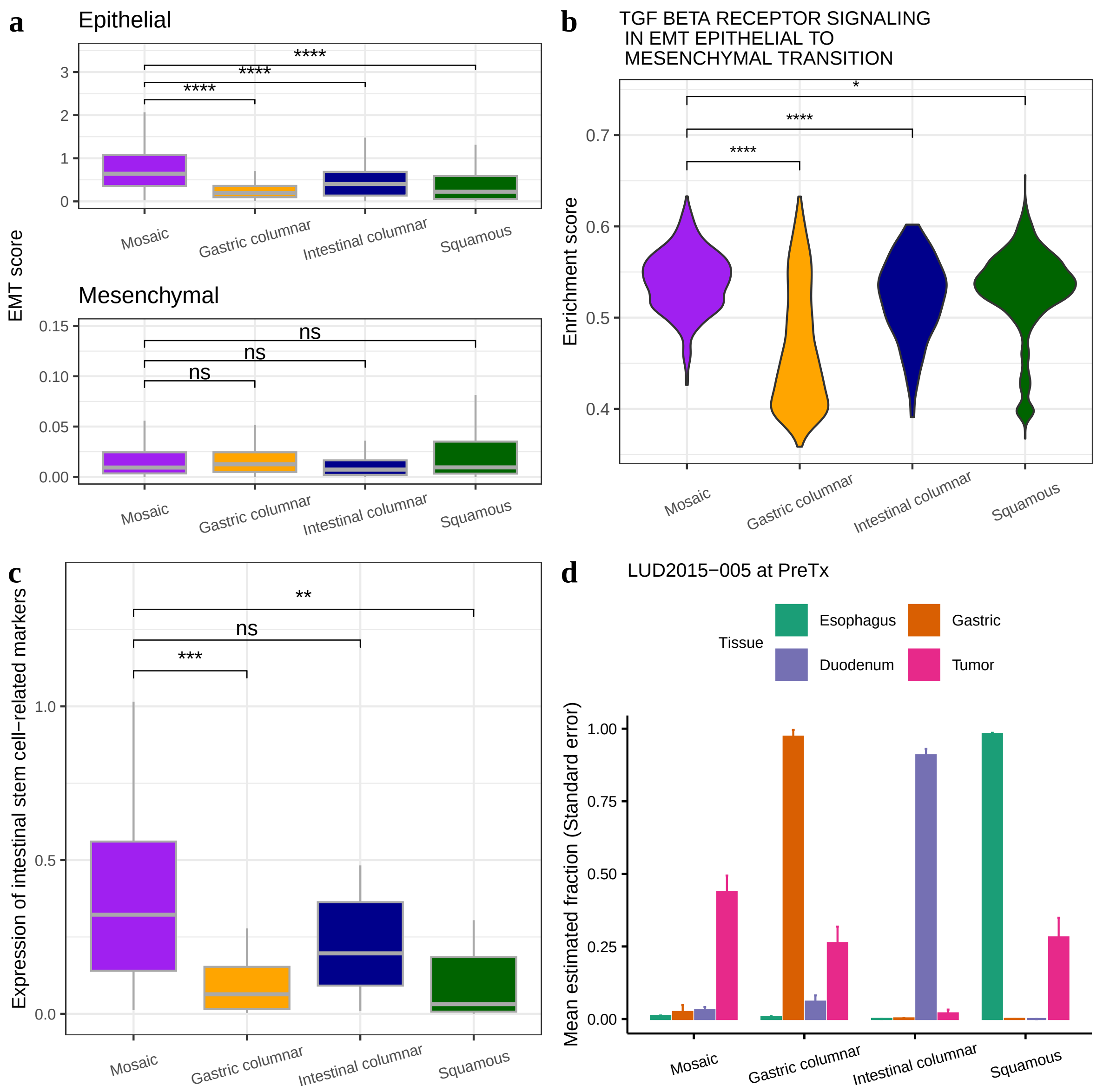


Figure S15: **Comparison of the cell composition estimates between EAC tumor and paired normal gastrointestinal tissue biopsies (esophagus, gastric, and duodenum) collected pre-ICI treatment (PreTx).** **a** Box plot of EMT scores for epithelial (top) and mesenchymal (bottom) markers across four epithelial cell populations. **b** Violin plot of enrichment scores in TGF $\beta$ receptor signaling in EMT across four epithelial cell populations. **c** Box plot comparing the expression of 18 intestinal stem cell-related marker genes across four epithelial cell populations. Statistical significance between mosaic and other three cell populations was assessed using a two-sided Wilcoxon test and indicated as follows: $^{ns}p \geq 0.05$, $^{*}p < 0.05$, $^{**}p < 0.01$, $^{***}p < 0.001$, and $^{****}p < 0.0001$. **d** The relative abundance of four epithelial cell populations in EAC tumor and normal tissues from LUD2015-005 cohort, colored by tissue of origin. Error bars represent standard deviation.

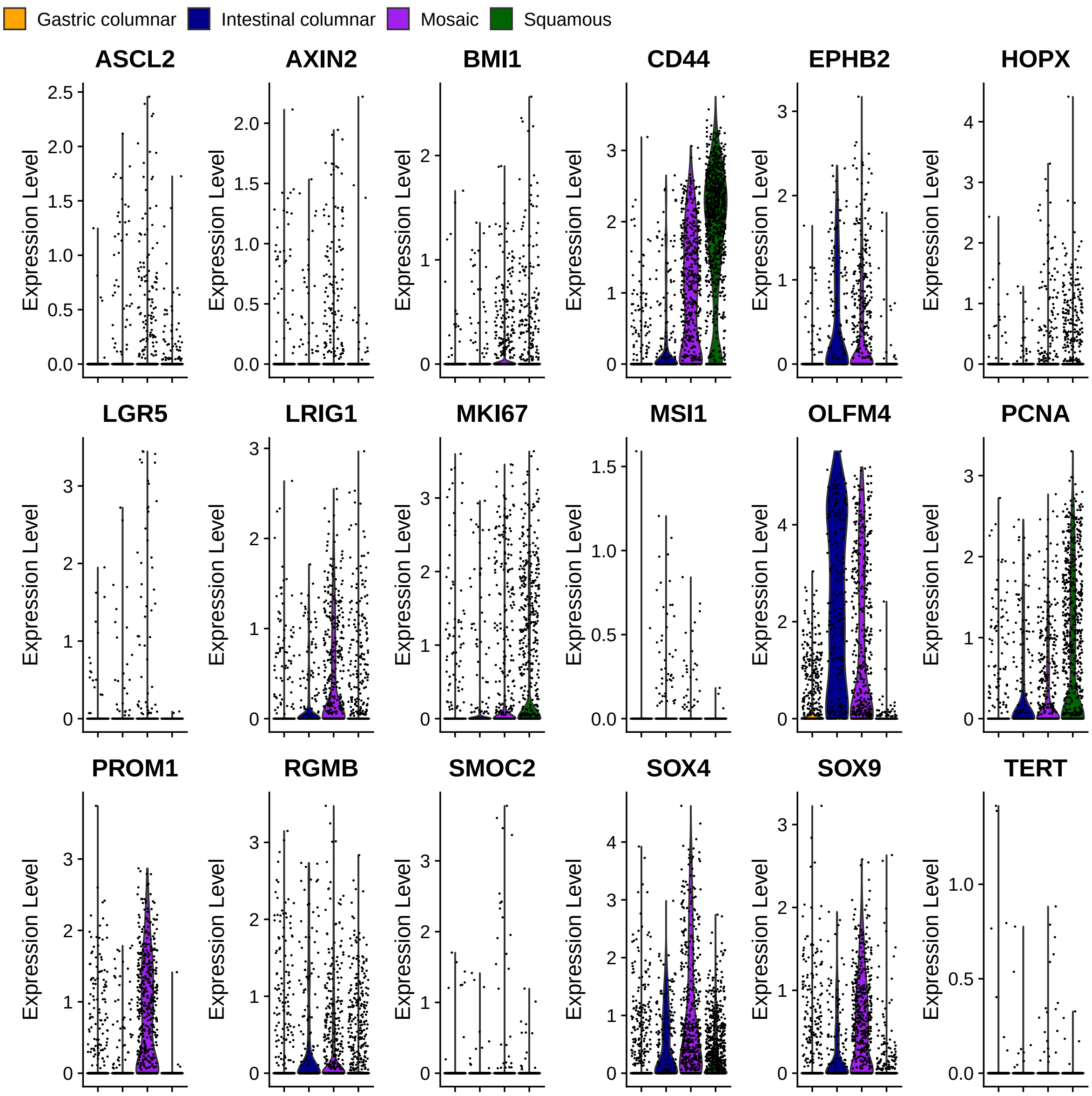


Figure S16: **Violin plots of the expression of 18 intestinal stem cell-related marker genes across gastric, intestinal, and mosaic columnar, and squamous cells.**

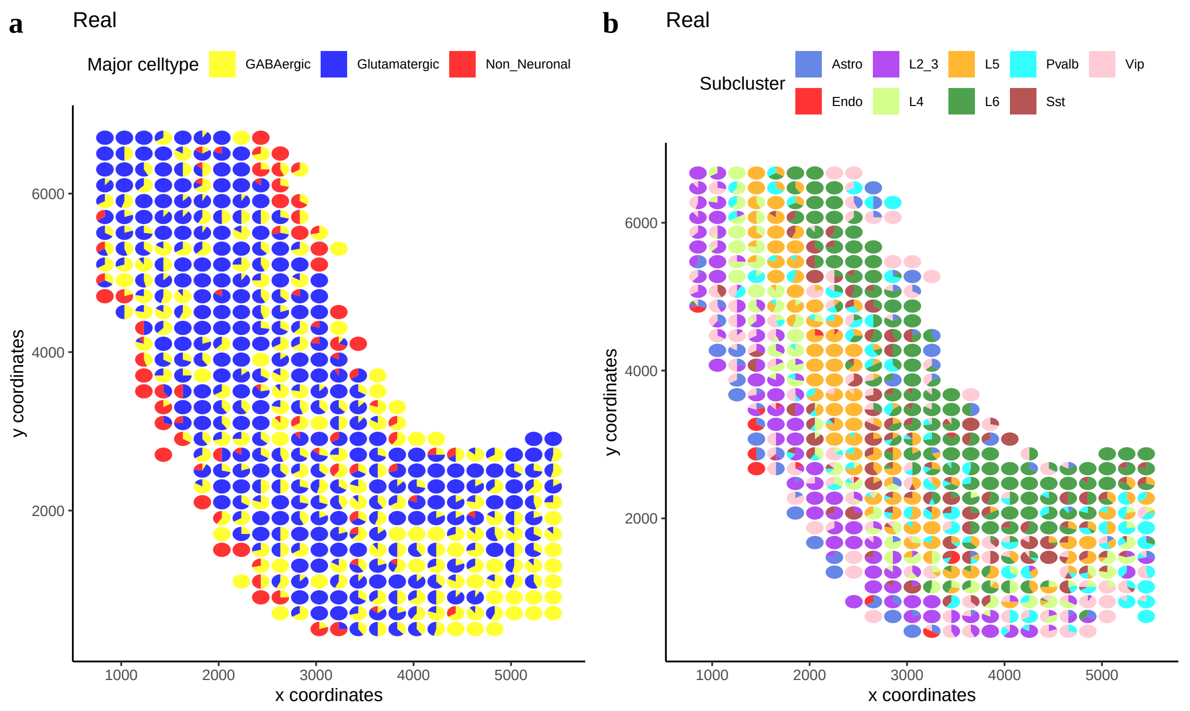


Figure S17: **Scatter pie plots showing the real proportions of major (a) and minor (b) subtypes at each spot in simulated ST data of mouse brain tissue.**

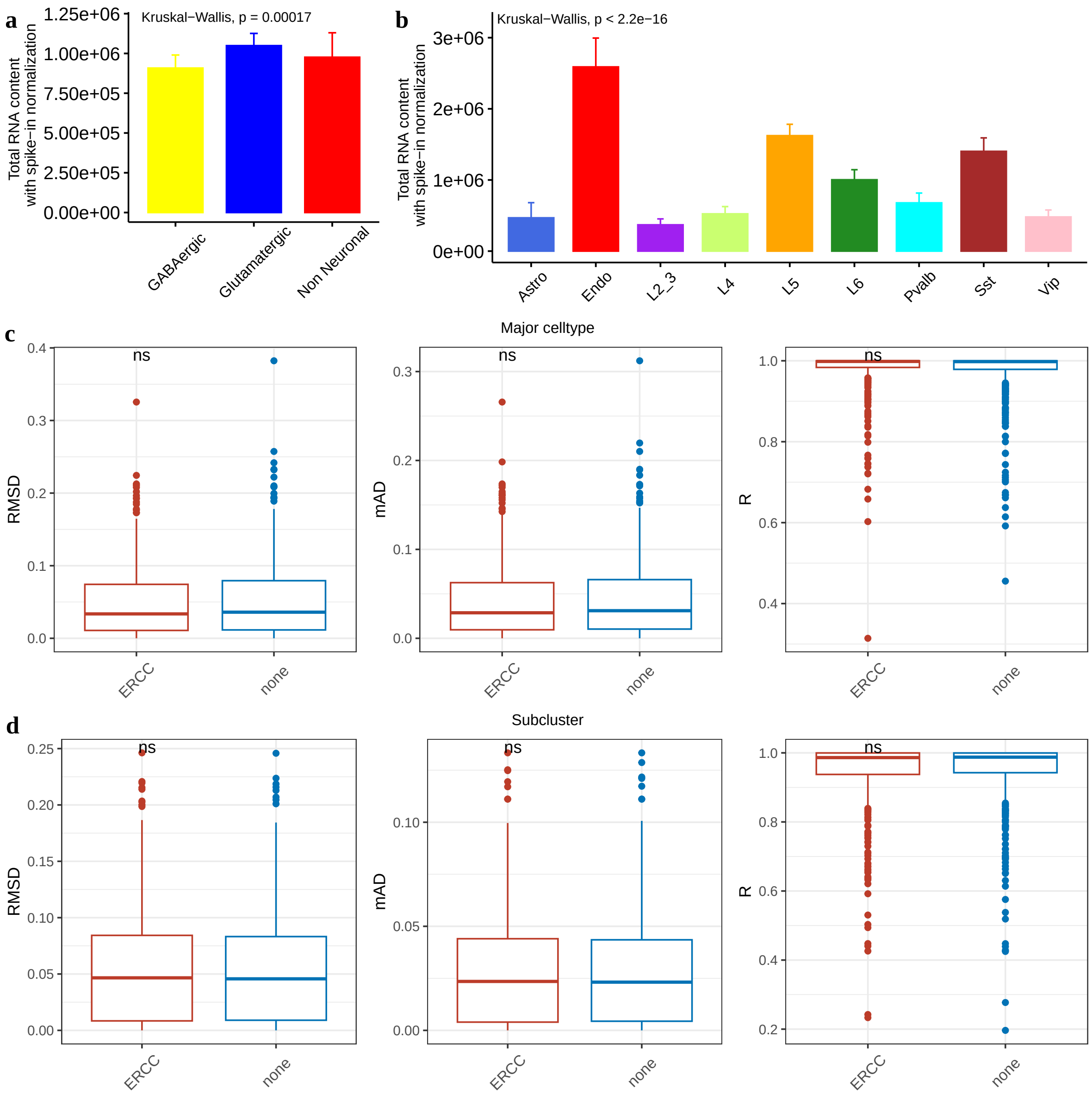


Figure S18: **CSsingle deconvolution performance in simulated ST data of mouse brain tissue: with versus without cell size correction.** (**a,b**) Comparison of the estimated cell sizes for three major (**a**) and nine minor (**b**) subtypes. Statistical significance was assessed using Kruskal-Wallis test. (**c,d**) Performance comparison for major (**c**) and minor (**d**) subtypes: with versus without cell size correction. Statistical significance was assessed using a two-sided Wilcoxon test and indicated as follows: $^{ns}p \geq 0.05$.

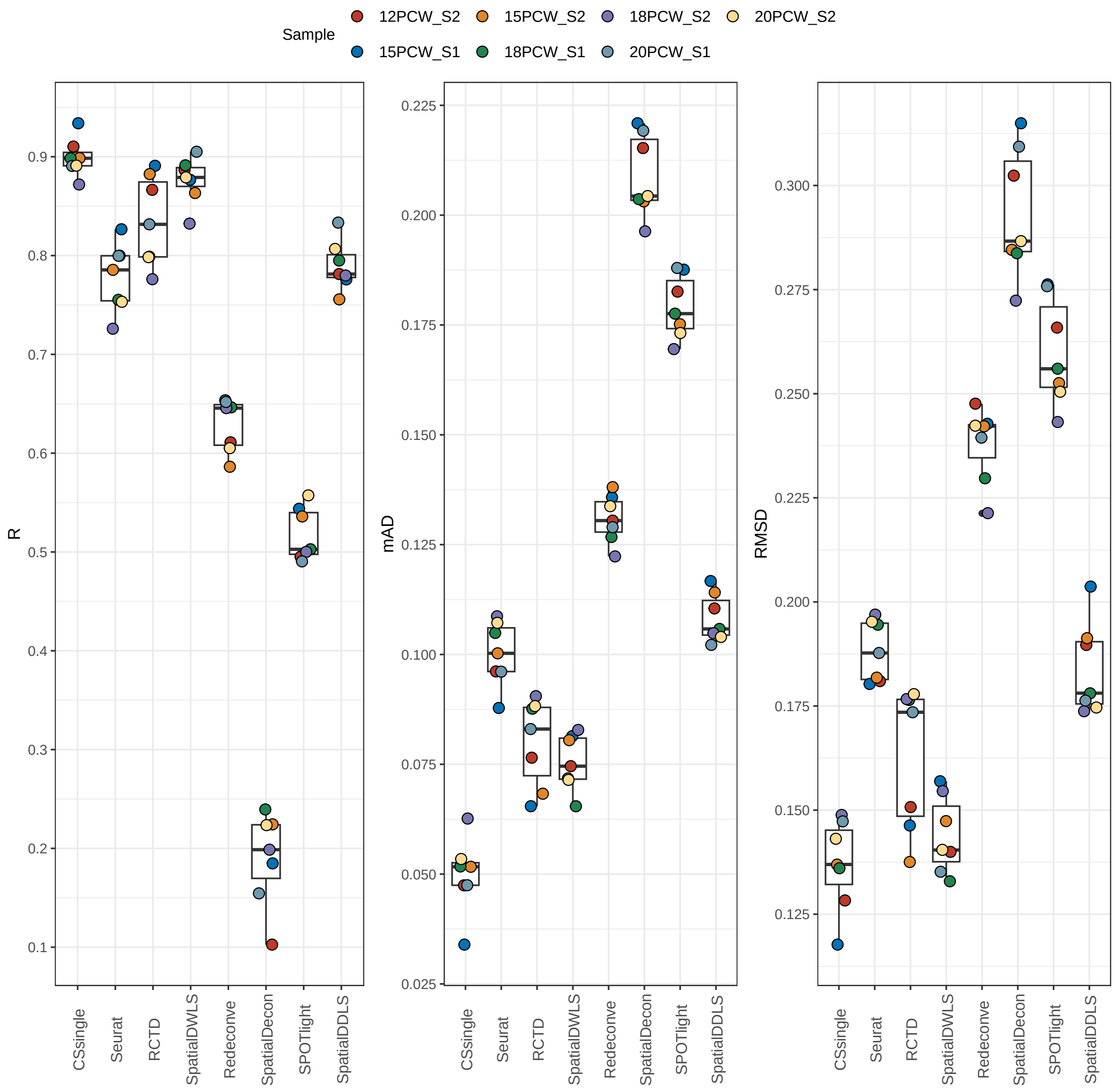


Figure S19: **Decomposition benchmark in simulated ST data of human pancreas.**

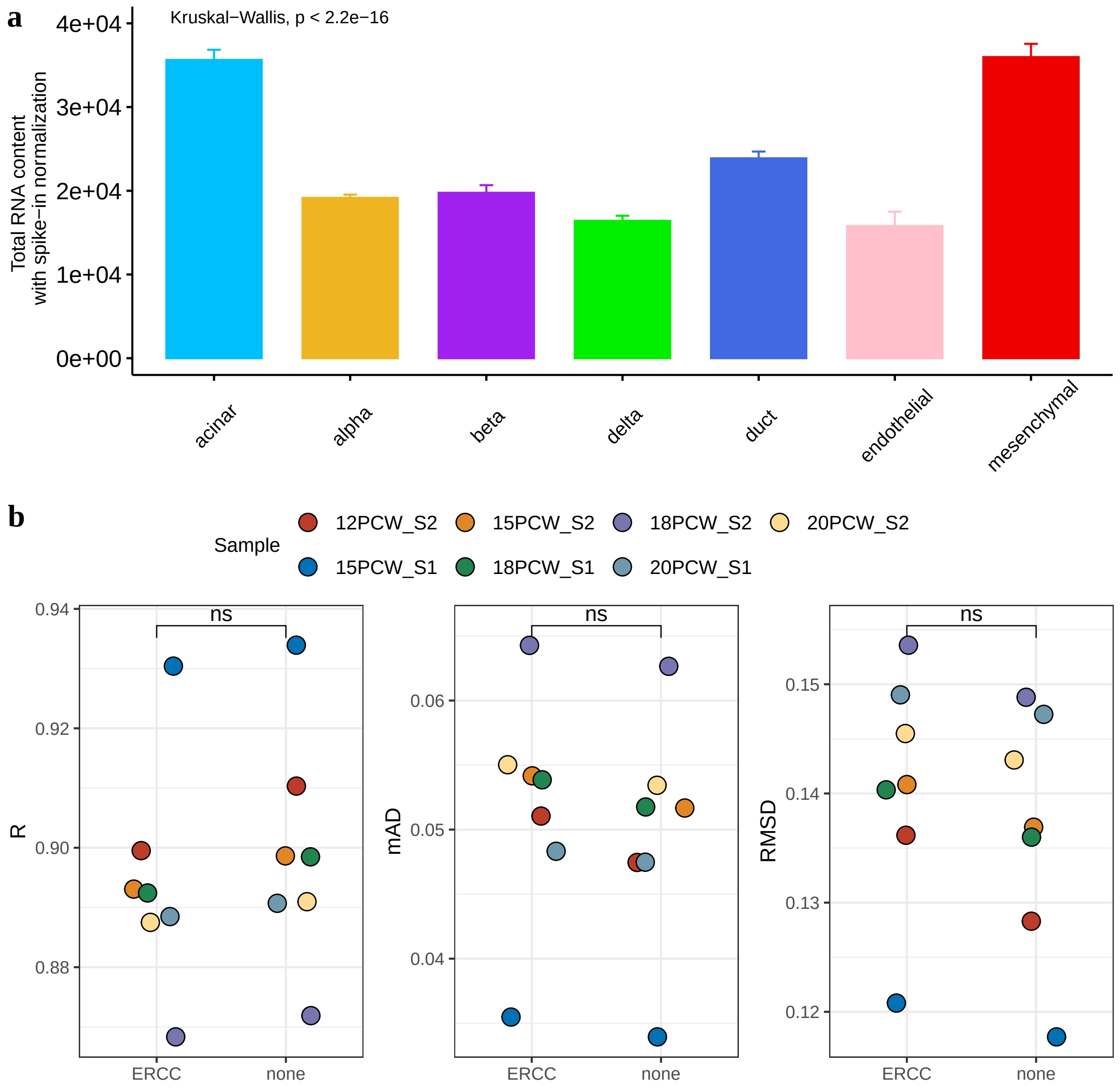


Figure S20: **CSsingle deconvolution performance in simulated ST data of human pancreas: with versus without cell size correction.** (**a**) Comparison of the estimated cell sizes for seven cell types. (**b**) Comparative evaluation of deconvolution performance: with versus without cell size correction. Points are color-coded by sample origin.

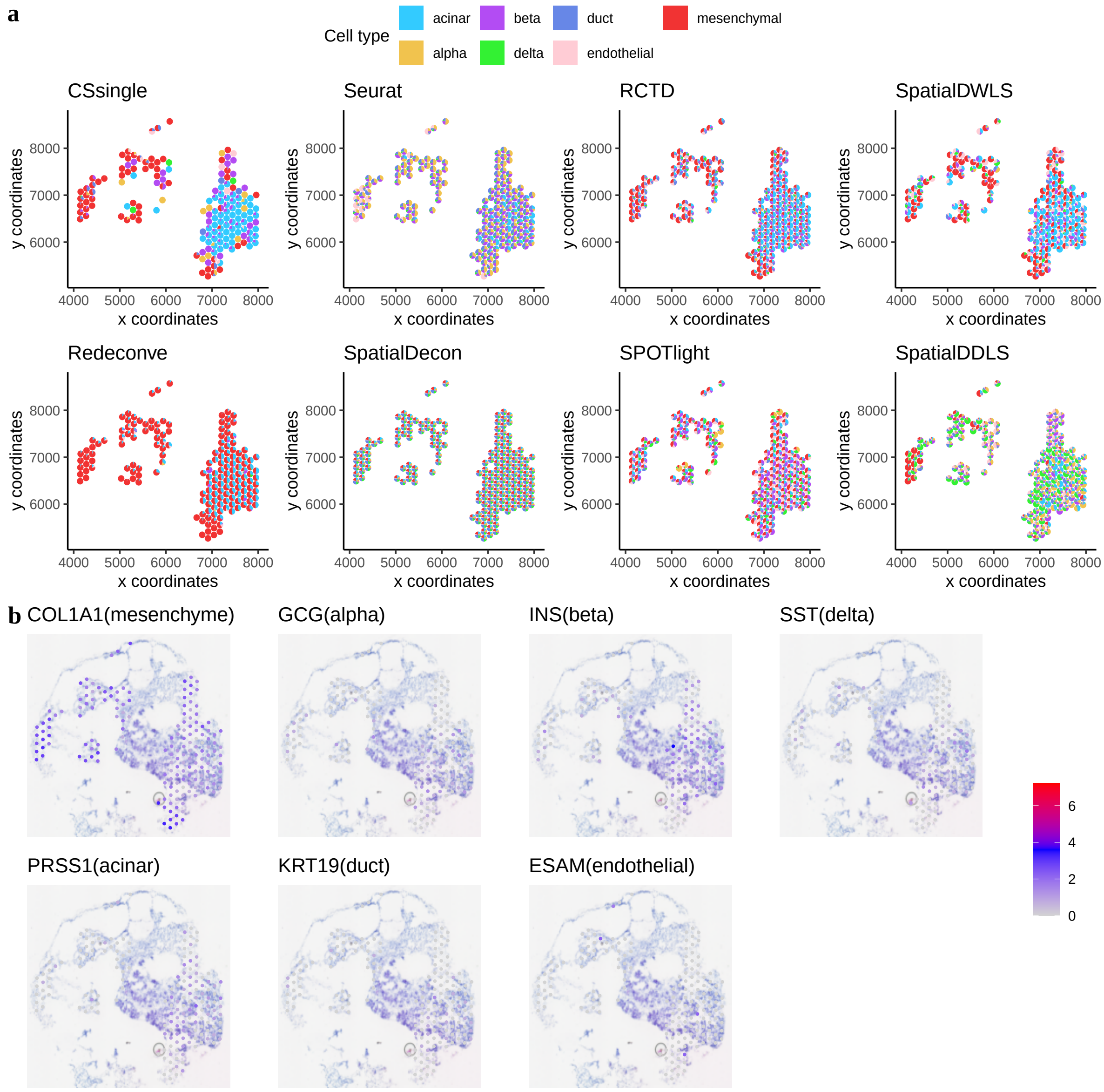


Figure S21: **Decomposition benchmark on tissue section 2 from human fetal pancreas at 12 PCW.** **a** Scatter pie plots showing the estimated cellular composition. **b** Feature plots showing some selected marker genes that were used to annotate the cell types (listed in parentheses).

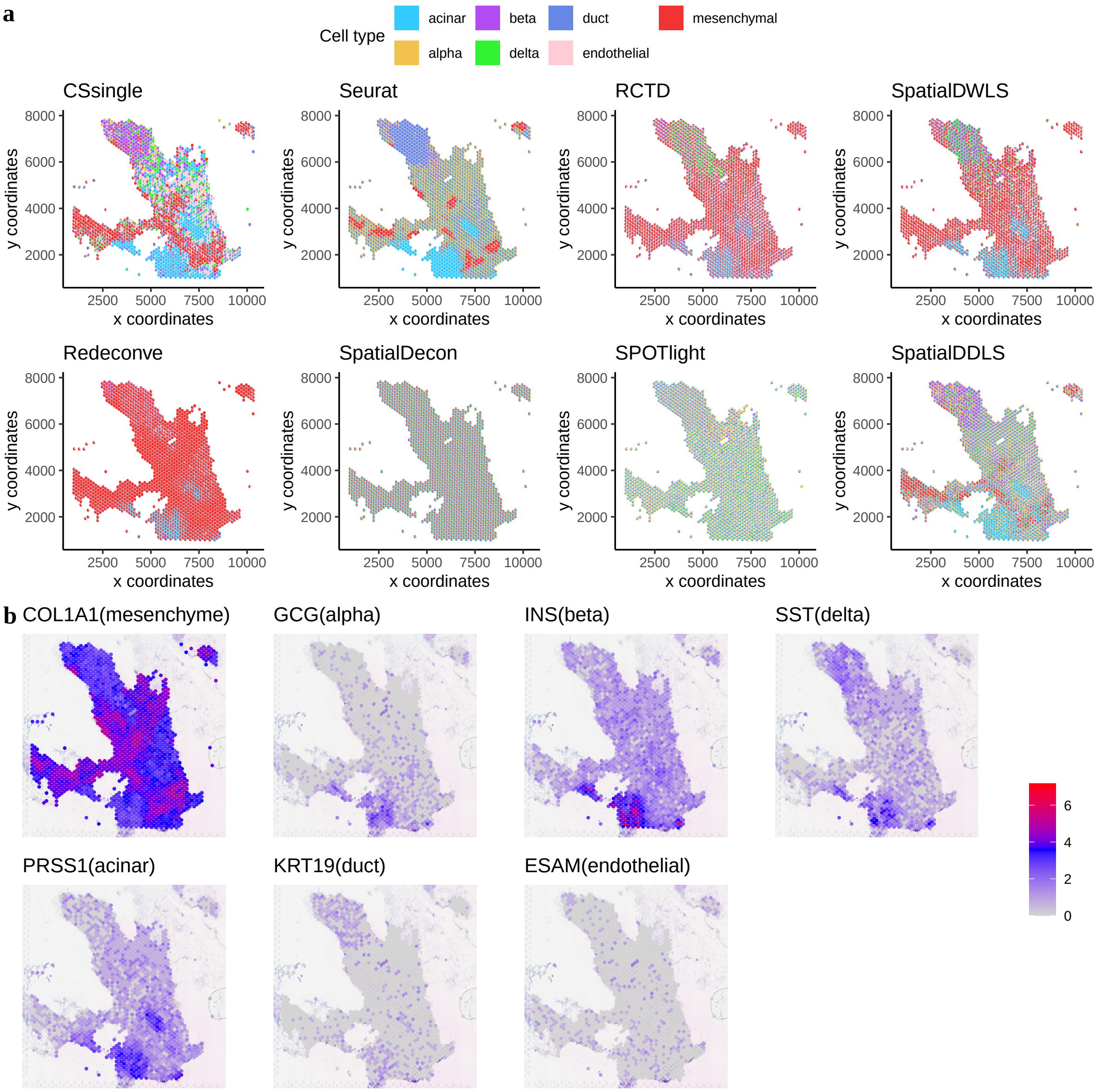


Figure S22: **Decomposition benchmark on tissue section 1 from human fetal pancreas at 15 PCW.** **a** Scatter pie plots showing the estimated cellular composition. **b** Feature plots showing some selected marker genes that were used to annotate the cell types (listed in parentheses).

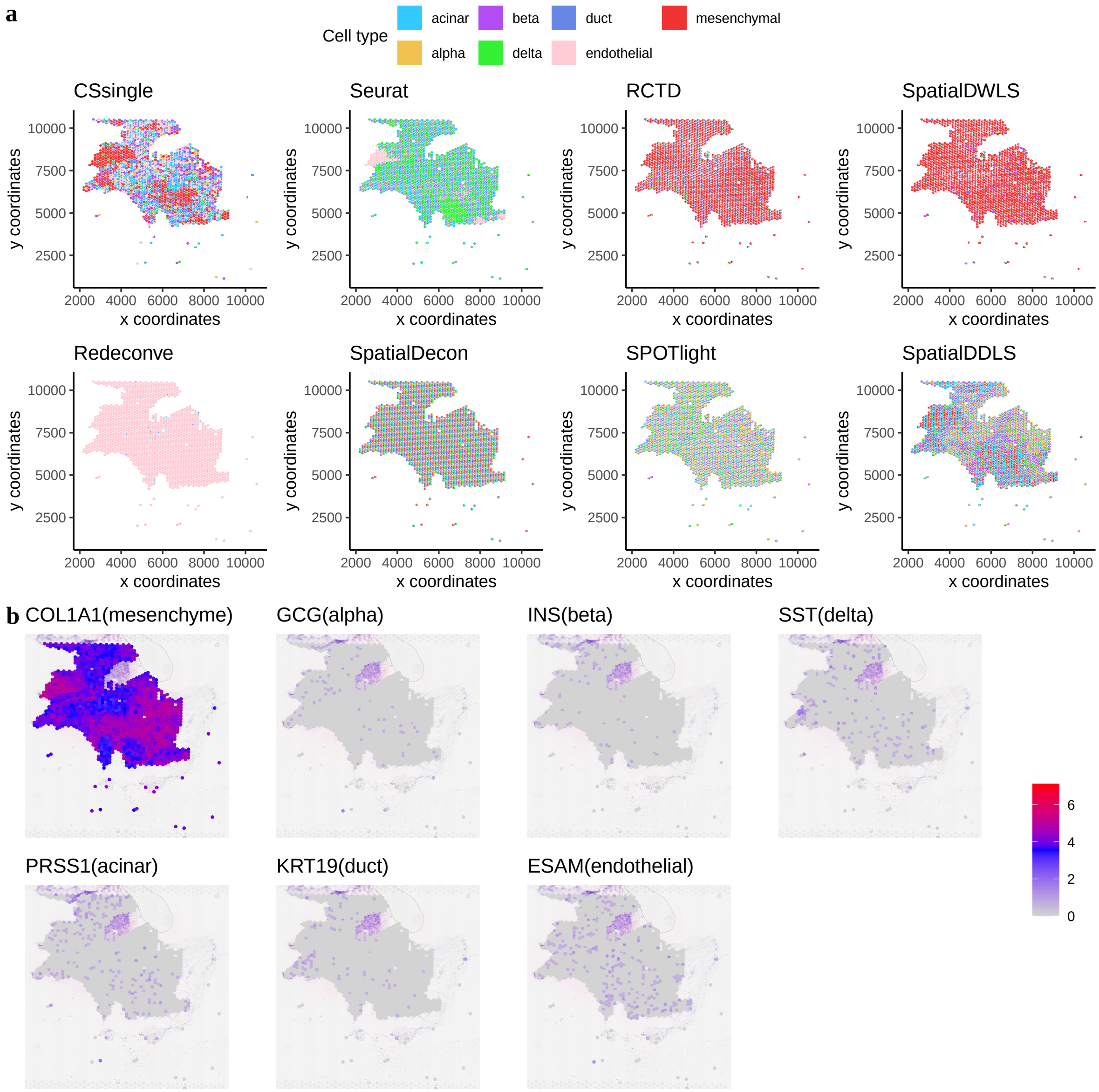


Figure S23: **Decomposition benchmark on tissue section 2 from human fetal pancreas at 15 PCW.** **a** Scatter pie plots showing the estimated cellular composition. **b** Feature plots showing some selected marker genes that were used to annotate the cell types (listed in parentheses).

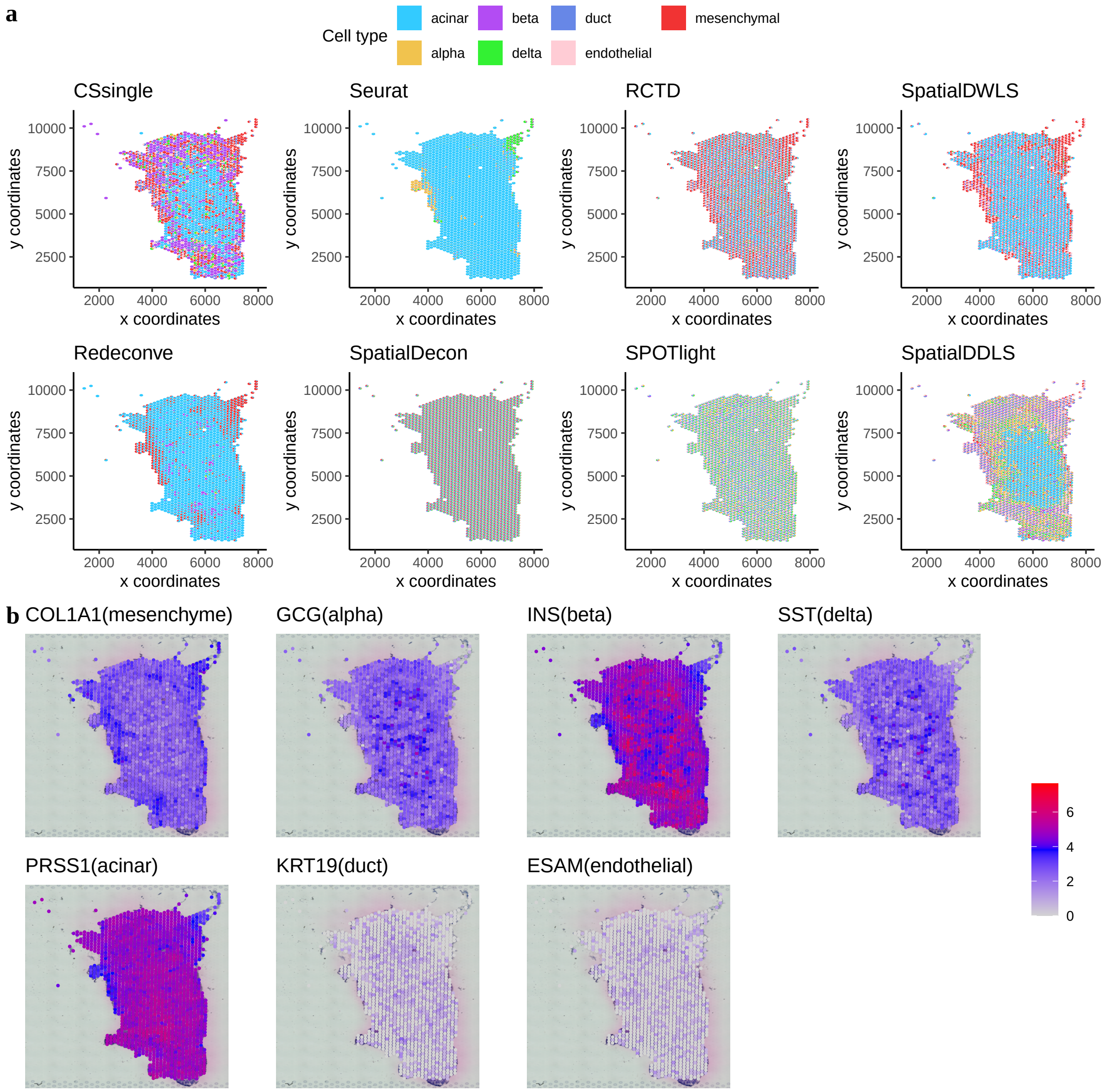


Figure S24: **Decomposition benchmark on tissue section 1 from human fetal pancreas at 18 PCW.** **a** Scatter pie plots showing the estimated cellular composition. **b** Feature plots showing some selected marker genes that were used to annotate the cell types (listed in parentheses).

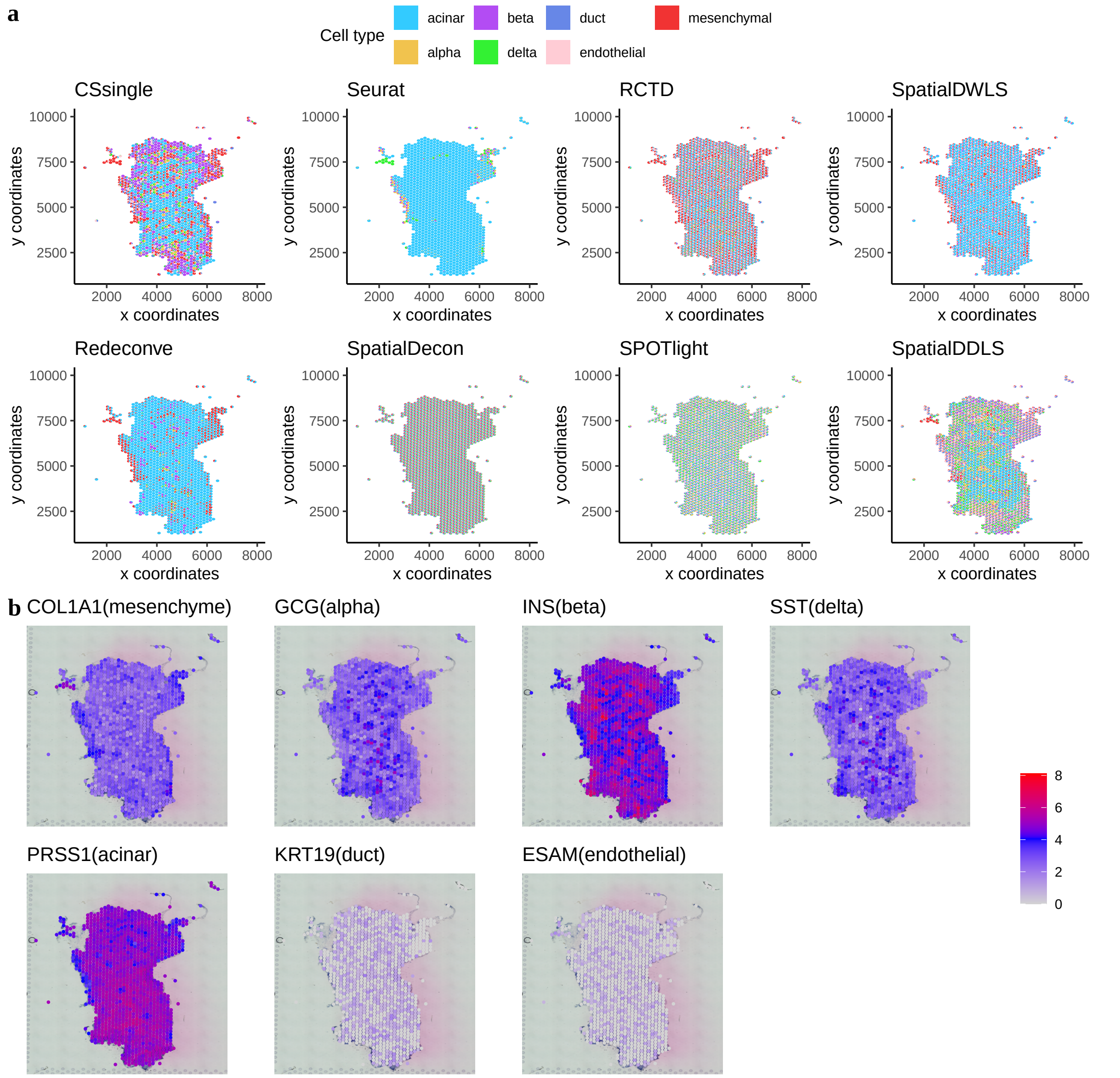


Figure S25: **Decomposition benchmark on tissue section 2 from human fetal pancreas at 18 PCW.** **a** Scatter pie plots showing the estimated cellular composition. **b** Feature plots showing some selected marker genes that were used to annotate the cell types (listed in parentheses).

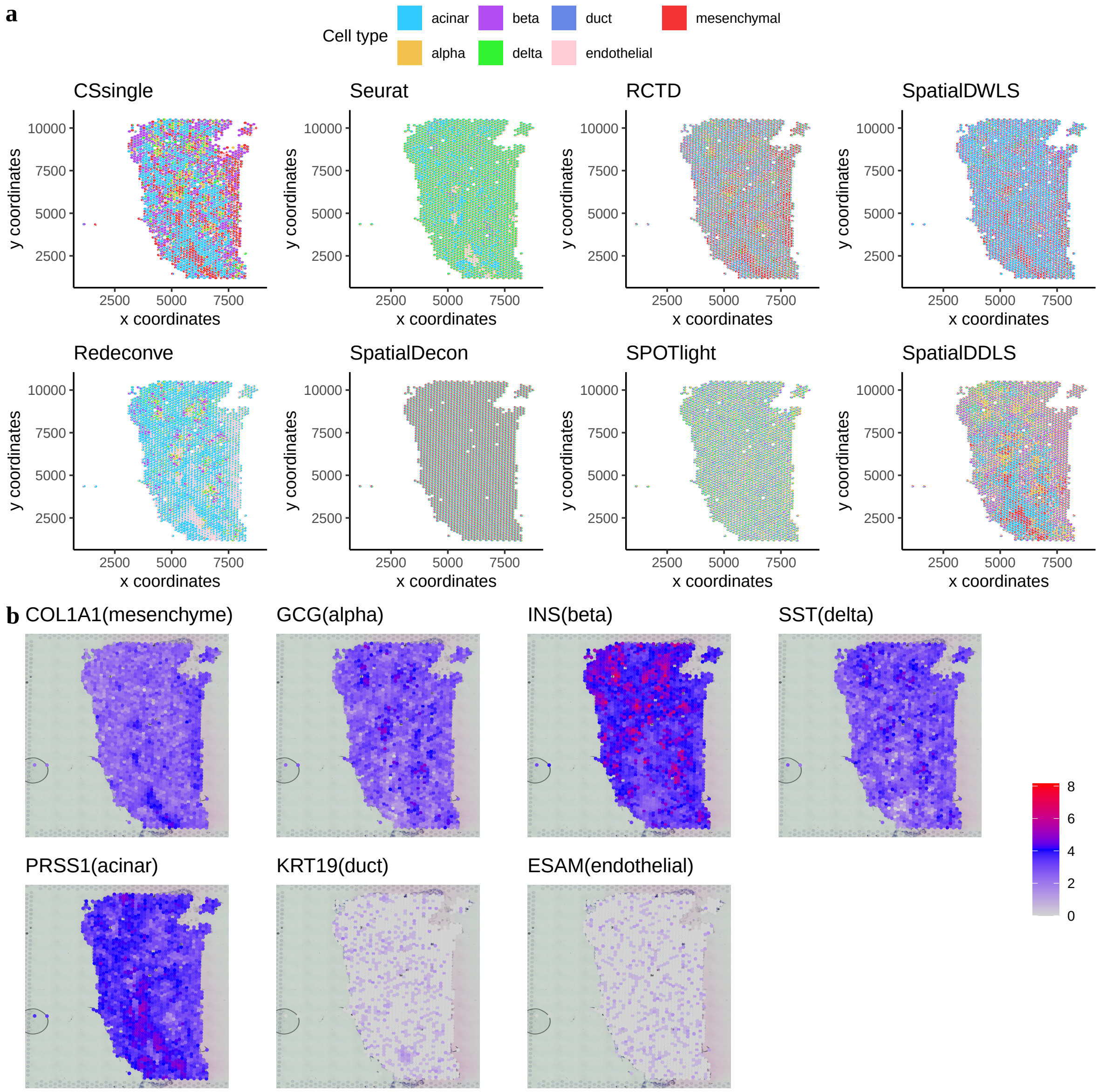


Figure S26: **Decomposition benchmark on tissue section 1 from human fetal pancreas at 20 PCW.** **a** Scatter pie plots showing the estimated cellular composition. **b** Feature plots showing some selected marker genes that were used to annotate the cell types (listed in parentheses).

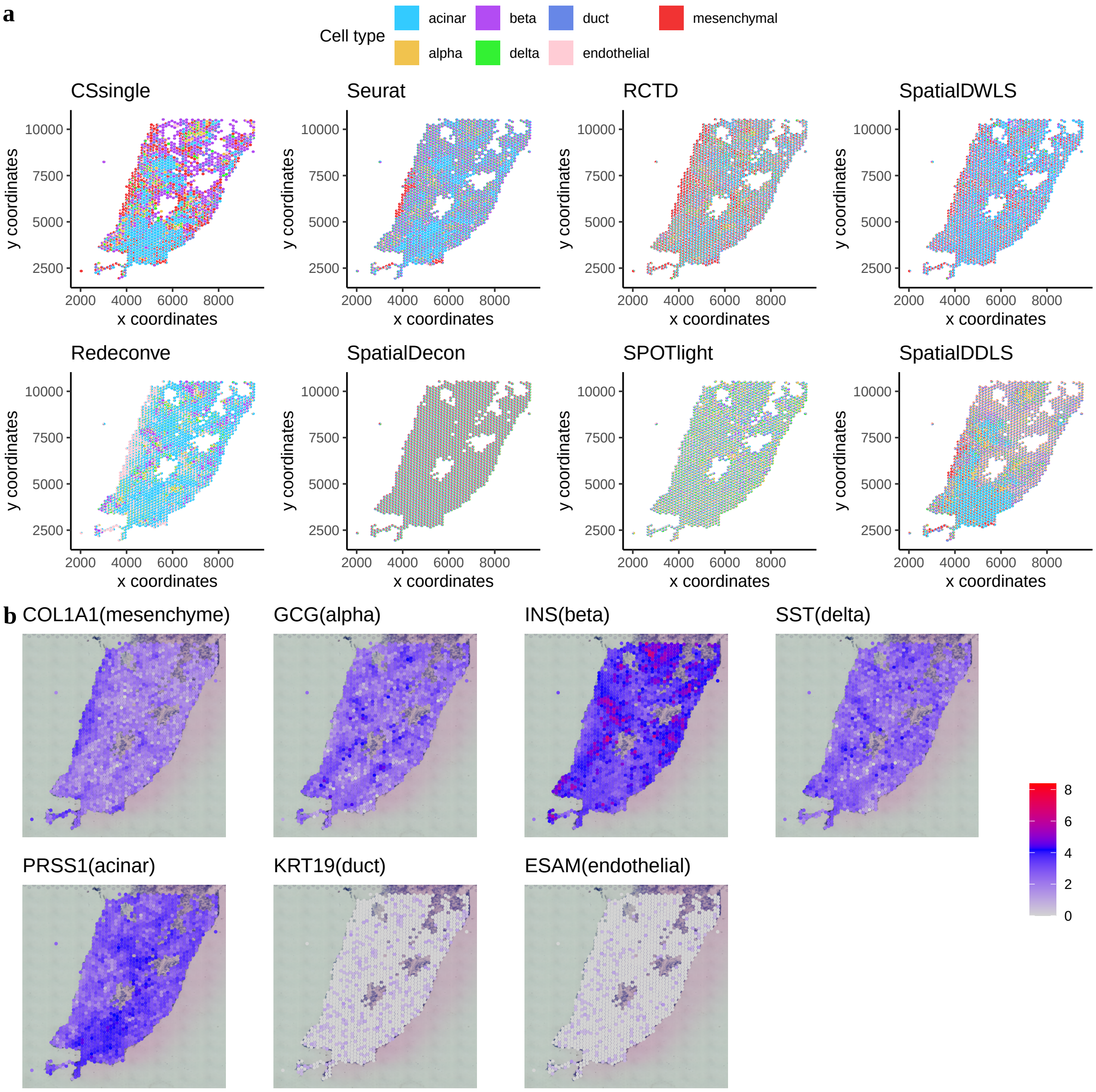


Figure S27: **Decomposition benchmark on tissue section 2 from human fetal pancreas at 15 PCW.** **a** Scatter pie plots showing the estimated cellular composition. **b** Feature plots showing some selected marker genes that were used to annotate the cell types (listed in parentheses).

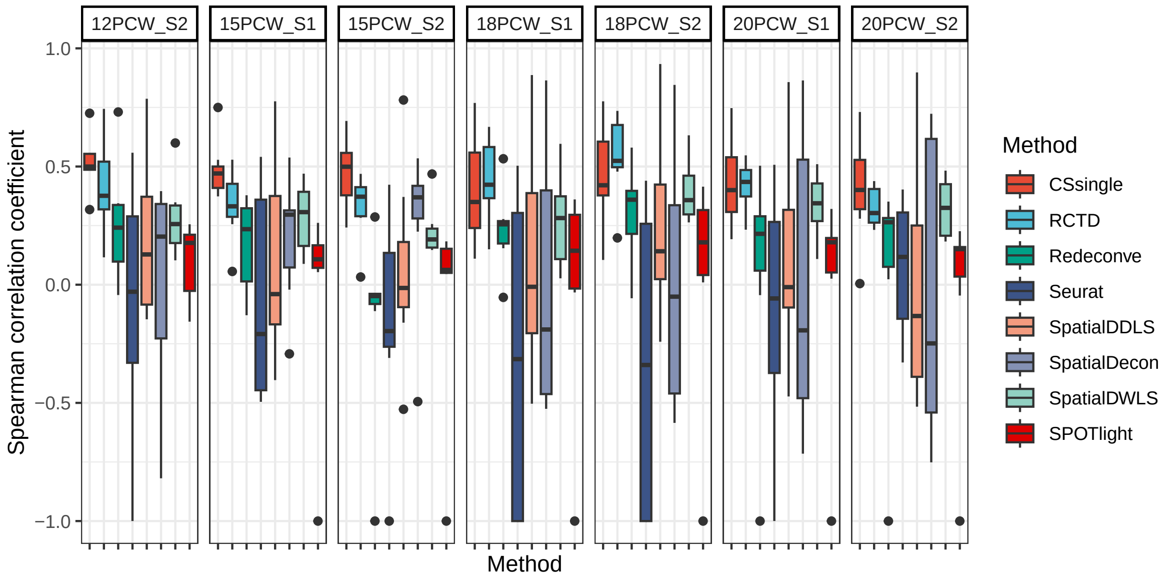


Figure S28: **Comparison of Spearman correlations between estimated cell type proportions and cell type enrichment scores.**

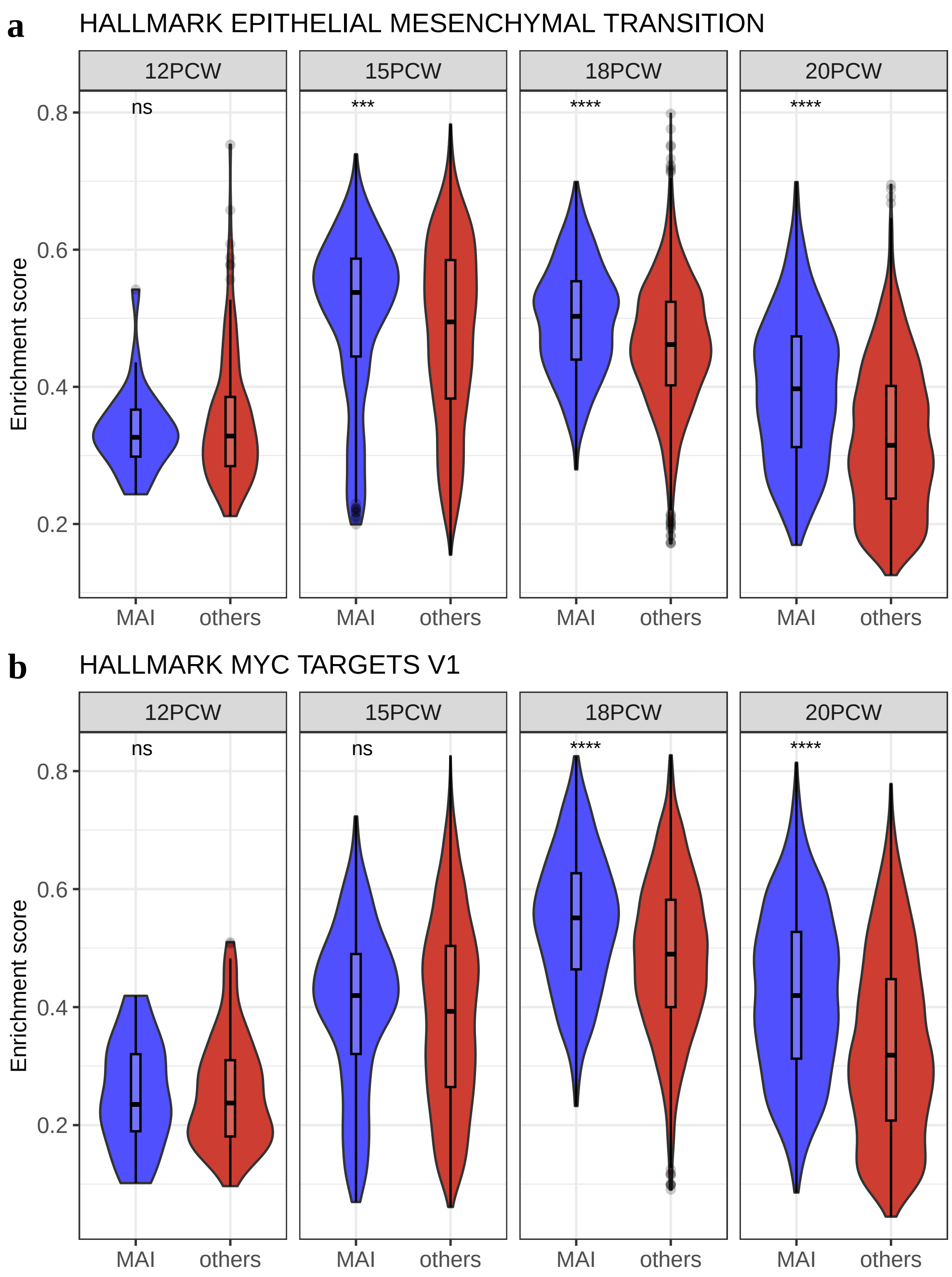


Figure S29: **Comparison of enrichment scores for Hallmark gene sets enriched in mesenchymal-acinar interactions (MAIs) versus other spots.** Enrichment of Hallmark gene sets for epithelial-mesenchymal transition (**a**) and MYC targets (**b**) in MAIs compared to other spots.

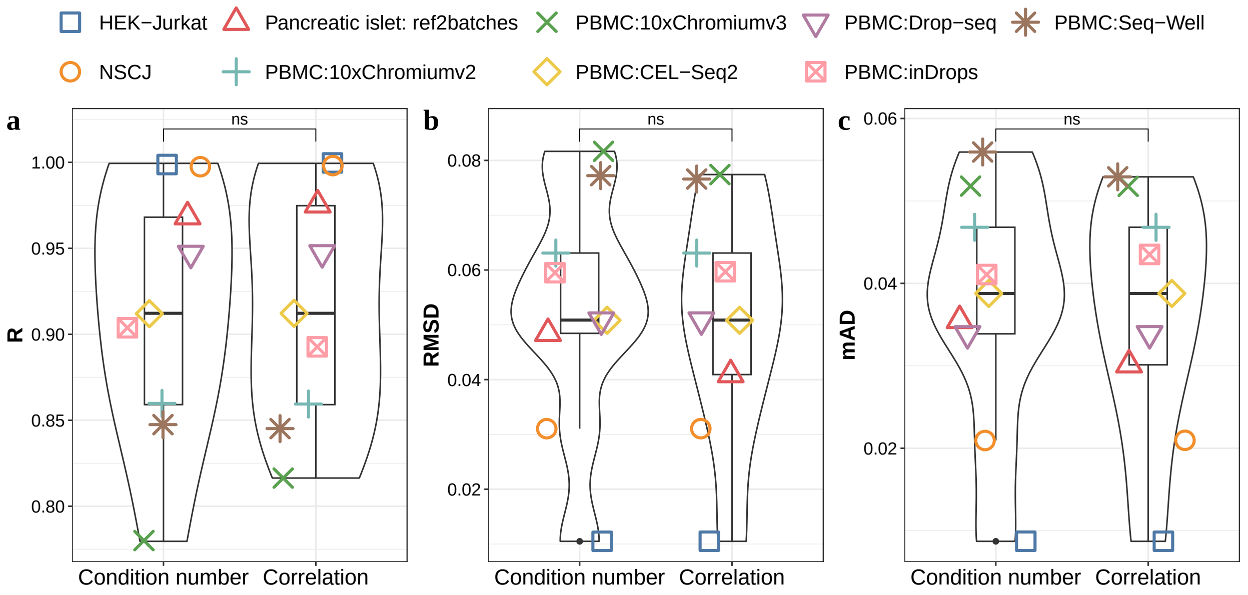


Figure S30: **Comparisons of the deconvolution results by integrating CSsingle with the signature matrix generated by different strategies: condition number versus spearman correlation.** The evaluation plot depicting Pearson correlation (**a**), root mean square deviation (**b**) and mean absolute deviation (**c**). Multiple signature matrices (six in total) were created by varying the number of marker genes from 50 to 300 with a step of 50 for each cell type. Nine data sets were denoted by different shapes and colors.

Table S1: Decomposition benchmark in human pancreatic islet tissue in terms of root mean square deviation (RMSD), mean absolute deviation (mAD) and Pearson correlation (R).

| Method | RMSD | mAD | R |
|---|---|---|---|
| CSsingle | 0.041 | 0.030 | 0.975 |
| DWLS | 0.071 | 0.052 | 0.952 |
| BayesPrism | 0.066 | 0.051 | 0.938 |
| CIBERSORT | 0.089 | 0.067 | 0.878 |
| CIBERSORTx | 0.125 | 0.084 | 0.755 |
| MuSiC | 0.112 | 0.090 | 0.855 |
| MuSiC2 | 0.120 | 0.094 | 0.844 |
| NNLS | 0.141 | 0.105 | 0.793 |
| SCDC | 0.178 | 0.120 | 0.583 |
| BisqueRNA | 0.122 | 0.082 | 0.798 |
| CAMmarker | 0.145 | 0.108 | 0.799 |
| EPIC | 0.189 | 0.137 | 0.522 |

Table S2: Performance of CSsingle and other methods in the human PBMC. The proportions of cell type in the bulk data are shown in parentheses. The RMSE, mAD and R values were averaged across six different scRNA-seq methods. The smallest RMSD and mAD, and the largest R values for each cell type are highlighted in bold.

| **B cells(0.152)** | RMSD | mAD | R | **Monocytes(0.152)** | RMSD | mAD | R |
|---|---|---|---|---|---|---|---|
| CSsingle | **0.02** | **0.02** | **0.98** | CSsingle | **0.03** | **0.02** | 0.94 |
| DWLS | 0.04 | 0.04 | **0.99** | DWLS | 0.1 | 0.09 | 0.96 |
| BayesPrism | 0.06 | 0.06 | **0.99** | BayesPrism | 0.14 | 0.13 | 0.96 |
| CIBERSORT | 0.09 | 0.08 | 0.95 | CIBERSORT | 0.07 | 0.06 | 0.87 |
| CIBERSORTx | 0.07 | 0.07 | 0.95 | CIBERSORTx | 0.08 | 0.07 | 0.92 |
| MuSiC | 0.14 | 0.13 | 0.61 | MuSiC | 0.1 | 0.09 | 0.83 |
| NNLS | 0.14 | 0.13 | -0.19 | NNLS | 0.22 | 0.2 | 0.69 |
| SCDC | 0.13 | 0.12 | 0.64 | SCDC | 0.08 | 0.07 | 0.8 |
| BisqueRNA | 0.06 | 0.05 | 0.56 | BisqueRNA | 0.09 | 0.09 | 0.05 |
| CAMmarker | 0.06 | 0.06 | 0.99 | CAMmarker | **0.03** | 0.03 | **0.97** |
| EPIC | 0.16 | 0.13 | 0.58 | EPIC | 0.46 | 0.4 | 0.54 |

| **T cells CD4+(0.220)** | RMSD | mAD | R | **NK cells(0.092)** | RMSD | mAD | R |
|---|---|---|---|---|---|---|---|
| CSsingle | **0.04** | **0.03** | 0.37 | CSsingle | 0.08 | 0.07 | 0.89 |
| DWLS | 0.12 | 0.11 | -0.39 | DWLS | **0.05** | **0.04** | 0.96 |
| BayesPrism | 0.16 | 0.15 | 0.32 | BayesPrism | 0.28 | 0.26 | **0.99** |
| CIBERSORT | 0.12 | 0.11 | -0.47 | CIBERSORT | 0.1 | 0.08 | 0.77 |
| CIBERSORTx | 0.16 | 0.13 | -0.56 | CIBERSORTx | **0.05** | **0.04** | 0.96 |
| MuSiC | 0.17 | 0.17 | -0.04 | MuSiC | 0.38 | 0.37 | 0.33 |
| NNLS | 0.27 | 0.26 | 0.02 | NNLS | 0.22 | 0.2 | 0.55 |
| SCDC | 0.15 | 0.15 | 0.24 | SCDC | 0.42 | 0.41 | 0.58 |
| BisqueRNA | 0.08 | 0.07 | **0.59** | BisqueRNA | 0.07 | 0.06 | 0.94 |
| CAMmarker | 0.12 | 0.12 | 0.52 | CAMmarker | 0.08 | 0.06 | 0.98 |
| EPIC | 0.21 | 0.21 | -0.03 | EPIC | 0.22 | 0.19 | 0.11 |

| **T cells CD8+(0.384)** | RMSD | mAD | R |
|---|---|---|---|
| CSsingle | **0.09** | **0.08** | 0.77 |
| DWLS | 0.13 | 0.12 | 0.9 |
| BayesPrism | 0.19 | 0.17 | 0.91 |
| CIBERSORT | 0.13 | 0.11 | **0.96** |
| CIBERSORTx | 0.14 | 0.12 | **0.96** |
| MuSiC | 0.3 | 0.28 | -0.1 |
| NNLS | 0.32 | 0.29 | -0.61 |
| SCDC | 0.24 | 0.22 | -0.08 |
| BisqueRNA | 0.11 | 0.09 | 0.53 |
| CAMmarker | 0.14 | 0.11 | 0.79 |
| EPIC | 0.35 | 0.32 | 0.04 |

Table S3: Top 25 marker genes for mosaic columnar cells. Genes were sorted by p-values in ascending order.

| p_val | avg_log2FC | pct.1 | pct.2 | p_val_adj | cluster | gene |
|---|---|---|---|---|---|---|
| 6.1E-280 | 3.038162 | 0.979 | 0.219 | 1.1E-275 | Mosaic | TFF3 |
| 1.7E-273 | 3.19257 | 0.979 | 0.246 | 3.1E-269 | Mosaic | REG4 |
| 8.0E-244 | 2.305866 | 0.977 | 0.289 | 1.5E-239 | Mosaic | LGALS4 |
| 2.3E-225 | 1.395139 | 0.841 | 0.117 | 4.3E-221 | Mosaic | PPP1R1B |
| 7.7E-221 | 1.501688 | 0.696 | 0.047 | 1.4E-216 | Mosaic | KRT7 |
| 9.4E-208 | 1.426637 | 0.969 | 0.389 | 1.7E-203 | Mosaic | EPCAM |
| 1.2E-205 | 1.151341 | 0.882 | 0.194 | 2.3E-201 | Mosaic | S100P |
| 5.1E-205 | 2.130895 | 0.749 | 0.084 | 9.4E-201 | Mosaic | CAMK2N1 |
| 6.0E-204 | 1.977627 | 0.915 | 0.208 | 1.1E-199 | Mosaic | MARCKSL1 |
| 1.5E-200 | 3.286145 | 0.797 | 0.11 | 2.8E-196 | Mosaic | SERPINA1 |
| 3.1E-200 | 3.783393 | 0.714 | 0.063 | 5.7E-196 | Mosaic | CEACAM5 |
| 4.0E-200 | 2.540482 | 0.828 | 0.132 | 7.2E-196 | Mosaic | CREB3L1 |
| 8.1E-198 | 2.819529 | 0.816 | 0.131 | 1.5E-193 | Mosaic | AGR3 |
| 1.6E-197 | 2.047543 | 0.952 | 0.268 | 3.0E-193 | Mosaic | LYZ |
| 2.3E-190 | 2.027108 | 0.923 | 0.278 | 4.3E-186 | Mosaic | GMDS |
| 3.6E-188 | 2.432044 | 0.774 | 0.108 | 6.7E-184 | Mosaic | AOC1 |
| 3.1E-184 | 2.752314 | 0.735 | 0.089 | 5.6E-180 | Mosaic | PROM1 |
| 1.4E-180 | 1.547915 | 0.801 | 0.147 | 2.5E-176 | Mosaic | ERN2 |
| 1.7E-180 | 1.097357 | 0.816 | 0.177 | 3.1E-176 | Mosaic | HNF4A |
| 4.1E-180 | 3.872445 | 0.551 | 0.019 | 7.4E-176 | Mosaic | LEFTY1 |
| 1.6E-179 | 3.336712 | 0.754 | 0.116 | 2.9E-175 | Mosaic | ATP2C2 |
| 4.8E-179 | 2.510736 | 0.816 | 0.149 | 8.8E-175 | Mosaic | FAM3D |
| 3.3E-178 | 1.854474 | 0.778 | 0.131 | 6.0E-174 | Mosaic | MUC13 |
| 1.1E-177 | 1.424082 | 0.884 | 0.217 | 2.0E-173 | Mosaic | SLC44A4 |
| 4.8E-176 | 1.893564 | 0.909 | 0.235 | 8.8E-172 | Mosaic | ELAPOR1 |

Table S4: Runtime efficiency and deconvolution accuracy for different step sizes used in the signature matrix construction. Multiple signature matrices were created by varying the number of marker gens from 50 to 300 with step 50 (six in total) or 1 (251 in total) for each cell type. All datasets were run on a 2.59 GHz Intel Xeon Processor with 256 GB of RAM and 32 cores. R's time module was used to obtain runtime measurements in seconds.

| | | | step size = 50 | | | | | step size = 1 | | |
|---|---|---|---|---|---|---|---|---|---|---|
| Datasets (# samples) | RMSD | mAD | R | Optimal # | Runtime(s) | RMSD | mAD | R | Optimal # | Runtime(s) |
| 10xChromiumv2 | 0.06 | 0.05 | 0.86 | 100 | 6.21 | 0.07 | 0.05 | 0.82 | 69 | 225.85 |
| 10xChromiumv3 | 0.08 | 0.05 | 0.82 | 100 | 5.7 | 0.08 | 0.05 | 0.82 | 84 | 223.95 |
| CEL-Seq2 | 0.05 | 0.04 | 0.91 | 50 | 6.38 | 0.05 | 0.04 | 0.91 | 50 | 241.34 |
| Drop-seq | 0.05 | 0.03 | 0.95 | 50 | 5.54 | 0.05 | 0.04 | 0.94 | 66 | 215.84 |
| inDrops | 0.06 | 0.04 | 0.89 | 100 | 5.54 | 0.06 | 0.05 | 0.88 | 81 | 206.28 |
| Seq-Well | 0.08 | 0.05 | 0.84 | 100 | 4.91 | 0.08 | 0.05 | 0.85 | 70 | 184.89 |
